# Phase separation behavior of TDP-43 governs its protein interactome and regulation of alternative splicing

**DOI:** 10.64898/2026.04.06.716630

**Authors:** Yelyzaveta Zadorozhna, Federico Uliana, Emanuele Zippo, Anke Busch, Niklas Kretschmer, Simone Mosna, Yongwon Suk, Jiaxuan Chen, Martina Hallegger, Carla Schmidt, Lukas Stelzl, Dorothee Dormann

## Abstract

TDP-43 is a nuclear RNA-binding protein that regulates RNA metabolism, including alternative splicing. Its aggregation is a major pathological hallmark of several neurodegenerative diseases. TDP-43 undergoes phase separation (PS) and this condensation behavior may be linked to aggregate formation. Whether and how PS governs TDP-43’s RNA regulatory functions remains poorly understood.

Here we utilized rationally designed mutations in the TDP-43 low complexity domain to tune TDP-43 PS, yielding a panel of TDP-43 variants with reduced propensity to form condensates (“PS-deficient”), and a panel forming irreversible, undynamic condensates (“solid-like”) *in vitro* and in cells. Two complementary interactomics approaches identified PS-dependent interactions between TDP-43 and key RNA regulatory factors, including splicing regulators and the RNA helicase UPF1, which show increased interactions with solid-like variants. Our results highlight that TDP-43 PS regulates RNA and protein homeostasis by modulating a subset of TDP-43-dependent alternative splicing events and by reshaping interactions with RNA regulatory factors.

## Introduction

TDP-43 is a nuclear RNA-binding protein (RBP) known to regulate many aspects of RNA metabolism, including alternative splicing, transcription, biogenesis of various micro-RNAs and long non-coding RNAs and alternative polyadenylation, while the smaller cytoplasmic pool of the protein is involved in the regulation of translation and mRNA transport in neurons^1^.

TDP-43 is implicated in several neurodegenerative disorders, including amyotrophic lateral sclerosis (ALS), frontotemporal dementia (FTD)^2,3^, up to 60% of Alzheimer’s disease cases^4,5^, and Limbic-predominant Age-related TDP-43 Encephalopathy (LATE)^6^. In these pathological conditions, TDP-43 is lost from the nucleus and accumulates in cytoplasmic inclusions in neurons and glial cells of affected brain regions, resulting in a nuclear loss-of-function^2,7,8^. This loss-of-function is further exacerbated by cytoplasmic TDP-43 aggregates, which progressively sequester TDP-43 and further deplete it from its physiological nuclear localization^9,10^. This causes disturbances of its essential RNA regulatory functions^9,10^, including altered splicing of neuron-essential genes^11–14^, alterations in translation^15^, changes in polyadenylation and mRNA stability^16–19^. Additionally, TDP-43 inclusions were found to co-aggregate with other cellular proteins, e.g. nuclear transport machinery components^20,21^, RNA processing factors, e.g. splicing and translation factors^22–24^ and UPF1, an RNA helicase involved in non-sense mediated decay (NMD)^16^, as well as protein quality control components^25,26^ and stress granule-associated proteins^27–29^. Some of these proteins also interact with TDP-43 under physiological conditions^30–33^, suggesting that TDP-43 may sequester some of its physiological interactors into pathological aggregates, possibly causing additional proteome perturbations and functional deficits.

How TDP-43 aggregates form is still not well understood. It has been proposed that a first step could be coalescence of TDP-43 into physiological or stress-induced biomolecular condensates, such as stress granules^34–36^, followed by intra-condensate demixing and TDP-43 aggregation^37^. *In vitro*, purified TDP-43 undergoes phase separation (PS) at physiological concentrations^38^, and in cells it is found in numerous nuclear and cytoplasmic condensates, including paraspeckles, Cajal bodies^39,40^ and neuronal transport granules^41^. TDP-43 PS is strongly mediated by its intrinsically disordered C-terminal low complexity domain (LCD). In particular, aromatic residues, such as tryptophans and phenylalanines, but also aliphatic residues and an alpha-helix in the conserved region (CR)^42–46^ contribute to a multivalent interaction network and drive PS. Additionally, oligomerization via the folded N-terminal domain is necessary for condensation of the full-length protein^38,40,47^.

To what extent PS of TDP-43 is necessary for its regulatory roles in the cell is not well understood. Generally, PS appears to be relevant for several DNA/RNA regulatory processes, including DNA damage repair^48,49^, transcription^50–57^, splicing ^58^, and translation^59^. For TDP-43, distinct PS phenotypes have been shown to dictate its RNA-binding repertoire and affect its roles in polyadenylation and autoregulation^60,61^, to increase global translation^62^, and to affect RNP granule transport for local translation in neurons^41,63^. Although these studies provided first evidence of the importance of TDP-43’s condensation behavior for cellular function, a system-wide view of the consequences of altered TDP-43 PS is still missing.

Here, we utilized a panel of TDP-43 variants with rational amino acid substitutions in the C-terminal LCD to bidirectionally tune TDP-43 PS through different types of mutations. Using *in vitro* assays with purified proteins we show that our designed TDP-43 mutants, on the one hand, include variants that have reduced condensation propensity combined with more dynamic condensate properties (“PS-deficient”), and, on the other hand, include variants that form amorphous condensates with lower dynamicity and reduced reversibility (“solid-like”). Utilizing cell lines with inducible expression of these different TDP-43 PS variants at near physiological levels and in the absence of endogenous TDP-43, we demonstrate that these PS properties are preserved in cells. Using immunoprecipitation and co-condensation assays in cellular lysate coupled to mass spectrometry, we find that solid-like TDP-43 variants exhibit enhanced interactions with several proteins involved in RNA binding and splicing regulation. One such example is UPF1, for which our data indicate that sequestration into solid-like TDP-43 condensates compromises the protein’s RNA regulatory function. Altogether, our data suggest that TDP-43 phase separation regulates RNA processing, and thus the proteome, in two ways. First, TDP-43 condensation affects alternative splicing of some of its target genes, leading to alterations in mRNA and protein levels. Second, differential TDP-43 condensation alters its interaction with other RNA processing proteins and potentially modulates their functionality, thereby causing broad indirect effects on cellular mRNA and protein levels.

## Materials and Methods

### Molecular dynamics simulations

Molecular dynamics (MD) simulations were performed using the CALVADOS3 coarse-grained model^64^. In this framework, each residue was represented by a single bead, connected by harmonic bonds, and solvent effects, including ionic strength (fixed at 150 mM), were treated implicitly through effective pairwise interactions. In particular, van der Waals interactions were modeled using the Ashbaugh-Hatch potential (a modified Lennard-Jones with attractive part scaled down based on the hydropathy of each specific amino acid) and the salt-screened electrostatics through a Debye-Hückel potential. Within folded regions, beads were positioned at the center of mass of each residue rather than at the Cα atom, and the structures were conserved using elastic networks, as described by Cao et al.^64^. Disordered regions, instead, were modeled as flexible chains.

Protein structures were obtained from AlphaFold2 predictions downloaded in January 2025^65^. Folded domains were defined based on domain annotations from the TED (The Encyclopedia of Domains) database (downloaded in August 2025)^66^.

All simulations were performed using Langevin dynamics with an integration time step of 0.01 ps and carried out using the python package HOOMD-blue v. 3.8.184^67^ together with GSD v. 2.8.1 for trajectory storage and analysis.

Simulation codes and the HOOMD-blue plugin for the Ashbaugh-Hatch pair potential are available on the following GitHub page: https://github.com/ezippo/hoomd3_phosphorylation, https://github.com/ezippo/ashbaugh_plugin ^68^.

#### In silico generation of TDP-43 condensates

The condensates in slab geometry were prepared by placing 100 chains of TDP-43 variants in a large cubic simulation box (35 * 35 * 35 nm), which was shrunk to a side length of 23 nm and then expanded in the z-direction to 150 nm. Periodic boundary conditions were applied in all directions. Initial chain positions and orientations were randomly assigned. Simulations were thermalized (1 µs) and run at a temperature between 250 and 290 K, for 5 × 10 integration steps, corresponding to 5 µs of simulated time (2 500 frames). For the computation of the density profiles, the center of mass of the condensate was centered in the simulation box at each frame and particle z-coordinates were binned at a resolution of 0.1 nm. For each z-position bin, particle counts were averaged over the trajectory, divided by the bin volume, and normalized to account for different protein length.

#### Calculation of contacts between TDP-43 and interaction pairs

20 random proteins from the list of TDP-43 interactors and 20 proteins from the dataset that were not identified as interactors (controls) were selected. In total the simulations have been performed on 40 interaction pairs.

For each interaction pair, we ran a 1×10^8^ integration steps (1 µs, 2 000 frames) long simulation with one TDP-43 chain and one interaction pair in a cubic box of 40 nm side length at 250 K. Intermolecular contacts between TDP-43 and its pairs were computed by pairwise minimum-image conversion under periodic boundary conditions. For each residue pair, a contact was defined when the inter-residue distance was smaller than the mean of the CALVADOS Lennard-Jones σ parameters (representing the diameters) of the two interacting amino acids.

Contact frequencies were calculated from residue–residue interaction datasets and normalized for partner protein length, and the number of analyzed frames. Statistical analysis was performed on normalized contact frequencies for both control and interaction datasets (Extended Data Fig. 4D).

Binding propensity was additionally quantified as the fraction of simulation frames containing at least one intermolecular contact (bound state). Apparent Kd was quantified as follows: 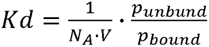 ^69^, statistical significance was assessed using a Wilcoxon rank-sum test (Extended Data Fig. 4C).

#### Simulation of the interaction between one chain of UPF1 and TDP-43 condensates

To simulate the interaction of UPF1 with TDP-43 condensates, one chain of UPF1 or albumin (ALB, control) was placed outside a previously prepared TDP-43 *in silico* condensate (1:100 UPF1 or ALB:TDP-43 ratio). Simulations were thermalized (1 µs) and run at 270 K, for 5 × 10 integration steps, corresponding to 5 µs of simulated time (2 500 frames). For each frame of the trajectory, the center of mass of the condensate was centered in the simulation box and particle z-coordinates were binned at resolution of 0.1 nm. For each z-position, bin counts were summed and normalized by the total number of particles of the corresponding species, yielding relative density distributions along the condensate axis. Contacts were calculated as detailed in the “Calculation of contacts between TDP-43 and interaction pairs” section.

### Cloning of constructs for bacterial expression

pJ4M TDP-43-MBP-His_6_ backbone^38^ (Addgene Plasmid #104480) was used to introduce mutations into the WT TDP-43 sequence for bacterial expression of recombinant proteins. Mutations were introduced using restriction cloning of synthetic DNA gBlocks (IDT) containing desired mutations. pJ4M TDP-43-MBP-His_6_ 12D/12A and G376V variants have been described earlier^70,71^. The mutations introduced into the distinct TDP-43 variants are listed in Supplementary Table 1.

### Cloning of constructs for the generation of mammalian HeLa Flp-In T-REx cell lines

pcDNA5-FRT-TO backbone (Invitrogen) expressing N-terminally myc-tagged TDP-43 WT^71^ was used to introduce PS-altering mutations. The TDP-43 sequence within this backbone is resistant to a TDP-43-specific siRNA (see “siRNA treatments”). The following mutations were introduced to confer siRNA resistance: c.1209 A>T, c.1212 C>T, c.1218 T>C, c.1221 T>A. Mutations spanning multiple residues (3W, ΔCR, 5R) were introduced as synthetic DNA gBlocks (IDT) via BsrGI and XhoI restriction sites. The G376V point mutation was introduced using site-directed mutagenesis with the Q5 polymerase and primers containing the desired mutation. pcDNA5-FRT-TO-myc-TDP-43-12D and pcDNA5-FRT-TO-myc-TDP-43-12A variants were generated earlier^71^.

pcDNA5-FRT-TO constructs containing EGFP-tagged TDP-43 WT, 12D, 12A, ΔCR and 5R were generated using Gibson cloning. pcDNA5-FRT-TO-EGFP-G376V was generated using site-directed mutagenesis. 3W mutations were first introduced into the pEGFP-C1-hTDP-43 vector^72^, and afterwards EGFP-3W insert was PCR-amplified and inserted into the pcDNA5-FRT-TO backbone.

### Expression and purification of TDP-43-MBP-His_6_ recombinant proteins

Recombinant TDP-43-MBP-His_6_ WT and its variants were purified according to Wang et al.^73^ with minor modifications. Briefly, proteins were expressed in BL21-DE3 Rosetta2 *E. coli*. Protein expression was induced with 0.5 mM IPTG and performed overnight at 16°C. Bacterial cell pellets were lysed in 50 mM Tris pH 8.0, 1 M NaCl, 10 mM imidazole, 10% glycerol, 4 mM β-mercaptoethanol, supplemented with aprotinin, leupeptin hemisulfate and pepstatin at 1 μg/μl each, as well as RNAse A and lysozyme (100 μg/ml each). Lysed material was sonicated and clarified by centrifugation. Protein purification was performed using Ni-NTA agarose beads (Qiagen) and elution was done using the lysis buffer containing 300 mM imidazole. Next, oligomeric and cleavage product species were removed by size exclusion chromatography on the Hiload 16/600 Superdex 200 pg column (GE Healthcare) in SEC buffer (50 mM Tris pH 8.0, 300 mM NaCl, 5% glycerol, 2 mM TCEP). Fractions of interest were concentrated and stored in the SEC buffer.

### Fluorescent labeling of TDP-43-MBP-His_6_

To enable fluorescent detection of TDP-43 condensates *in vitro*, TDP-43-MBP-His_6_ proteins were labeled at cysteine residues using Alexa Fluor 488 C5 maleimide (Thermo Fisher) at a low labeling efficiency (2 - 4%) to avoid potential effects of the dye on TDP-43 condensation ability. Protein was incubated with the dye at 100:7 (protein:dye) molar ratio for 2 hours at room temperature (RT). Excess dye was removed by 4 sequential washes using the Amicon Ultra protein concentration columns (30 kDa MWCO, Merck Millipore). For assays where several TDP-43 variants were compared side by side, a maximum of 1% difference in labeling efficiency between protein variants was allowed.

### Phase separation assays with purified proteins

Prior to all experiments, proteins were centrifuged at 20 000 g for 10 minutes at 4°C to remove any preformed aggregates/insoluble protein. Phase separation assays were performed in phase separation buffer (20 mM HEPES pH 7.5, 150 mM NaCl, 2 mM DTT). TDP-43 condensation was induced by incubating TDP-43-MBP-His_6_ (WT or variants) at the indicated concentration with TEV protease (4 μM final concentration) and reactions were incubated for the indicated amount of time (15 - 60 minutes) at RT. Phase separation reactions for brightfield microscopy and FRAP experiments were set up in µ-Slide 18-Well-Flat IbidiTreat microscope chambers (Ibidi 81826), for fluorescence microscopy in 384-well plates pre-coated with 10% Pluronic F-127, and for all other assays in Eppendorf tubes.

### Brightfield microscopy: Imaging of unlabeled TDP-43 condensates

Imaging of condensates formed by unlabeled TDP-43 was performed on a Zeiss Axio Observer 7 microscope in the phase contrast mode using a Plan-Apochromat 63x/1.4 Oil Ph3 objective, an Axiocam 705 monocamera and the Zeiss Zen software.

### Confocal microscopy: Imaging of fluorescently labeled TDP-43 condensates

Prior to the phase separation reaction setup, 384-well plates were coated with 10% Pluronic F-127 for 1 hour at RT and washed with water. Phase separation reactions were performed as described above and imaged using a Revvity Opera Phenix high content confocal microscope (IMB Mainz Microscopy Core Facility) equipped with the Revvity Harmony analysis software. Imaging was performed using the 63x water immersion objective, NA 1.15, binning 1. Alexa 488 imaging channel was used (excitation at 488 nm, emission range: 500 - 550 nm, time set to 60 ms, power set to 80%).

### Condensate reversibility assay

Phase separation reactions were set up in test tubes using 5 μM TDP-43-MBP-His_6_ as described in the “Phase separation assay with purified proteins” section. After the addition of TEV protease, reactions were incubated at RT for 15 or 60 minutes. After the incubation, the “1:1” reactions were immediately centrifuged at 20 000 g for 15 minutes, 4°C. Dilution reactions (“1:5” and “1:10”) were diluted with phase separation buffer (20 mM HEPES pH 7.5, 150 mM NaCl, 2 mM DTT) 1:5 and 1:10 respectively, incubated for 15 minutes at RT to enable condensate dissolution, and centrifuged at 20 000 g for 15 minutes, 4°C. Proteins from supernatant and pellet fractions were subsequently separated using SDS-PAGE, followed by western blotting with a rabbit anti-TDP43 antibody diluted 1:1 000 (Proteintech, 80001-RR). western blot signal was quantified as % soluble, where 100% indicates the sum of pellet and supernatant signal from undiluted 1:1 samples, and the signal coming from the 1:1, 1:5 and 1:10 supernatant samples is represented as a percentage of that value.

### C_sat_ (saturation concentration) estimation

For each experiment, phase separation reactions were set up in triplicate using 1 μM TDP-43-MBP-His_6_ as described in the “Phase separation assay with purified proteins” section. After the addition of TEV protease, reactions were incubated at RT for 60 minutes and subsequently centrifuged at 20 000 g for 15 minutes, 4°C. Obtained pellet and supernatant fractions were analyzed using SDS-PAGE, followed by SYPRO Ruby protein staining. Briefly, gels were fixed in 50% EtOH, 10% (v/v) acetic acid for 30 minutes at RT. Afterwards, SYPRO Ruby solution was applied overnight. Finally, gels were destained in 10% EtOH, 10% acetic acid (v/v) for 30 minutes at RT and visualized. To obtain C_sat_ values, pellet and supernatant signals were quantified. Assuming the sum of pellet and supernatant signal is 1 μM TDP-43, a corresponding concentration value was assigned for the signal in the supernatant alone, indicating an approximate saturation concentration. Each experimental replicate was conducted using three independent phase separation reactions.

### Stable cell line generation and cell culture

Parental HeLa Flp-In T-REx host cell line (gift of Christian Behrends, LMU Munich) was grown in Dulbecco’s Modified Eagle Medium (DMEM), supplemented with 10% Tet-System Approved fetal bovine serum (Tet-FBS, Thermo Fisher Scientific, A4736401), 10 μg/ml gentamycin and 100 μg/ml Zeocin in a humidified incubator at 37°C and 5% CO_2_. To generate HeLa Flp-In T-REx cell lines with inducible expression of a single copy of myc-or EGFP-tagged TDP-43 (WT or variants), HeLa Flp-In T-REx host cells were transfected with a pcDNA5-FRT-TO vector containing the TDP-43 variant of interest and a pOG44 Flp recombinase vector (Invitrogen) at a 1:10 ratio (pcDNA5:pOG44) using Lipofectamine 2 000 according to manufacturer’s instructions. After transfection, cells were placed into selection medium (DMEM supplemented with 10% Tet-FBS, 10 μg/ml gentamycin, 150 µg/ml hygromycin B and 10 µg/ml blasticidin). Once an antibiotic-resistant stable population was established, cells were FACS-sorted using a Bigfoot cell sorter (Invitrogen, IMB Mainz Flow Cytometry core facility) using the 100 µm nozzle at 30 psi based on forward and side scatter profiles and seeded individually for clonal growth. HeLa Flp-In T-REx EGFP-TDP-43 cell lines were induced with 1 μg/ml of doxycycline prior to FACS and sorted based on EGFP fluorescence intensity. Individual clones were screened for homogeneity by Opera high-throughput imaging based on EGFP signal or using an anti-myc immunofluorescence staining (mouse 9E10 antibody, IMB Protein Production core facility). Total levels of each expressed TDP-43 variant were also confirmed by western blotting using a mouse anti-TDP-43 antibody (diluted 1:2 000, Proteintech 60019-2-Ig). The amount of TDP-43 expressed was further adjusted by titrating the concentration of doxycycline (see below).

Established cell lines were grown in DMEM supplemented with 10% Tet-FBS, 10 μg/ml gentamycin, 150 µg/ml hygromycin B and 10 μg/ml blasticidin. The expression of TDP-43 variants was induced by the addition of doxycycline for 24 hours. The following concentrations were used for individual myc-TDP-43-expressing cell lines: 0.002 μg/ml (WT, 3W, ΔCR), 0.008 μg/ml (5R), 0.03 μg/ml (12A), 0.5 μg/ml (G376V) and 10 μg/ml (12D). For EGFP-TDP-43-expressing cell lines, the following doxycycline concentrations were used: 0.05 μg/ml (WT), 0.1 μg/ml (ΔCR, G376V), 0.5 μg/ml (12D, 5R, 12A), 1 μg/ml (3W).

The parental HeLa Flp-In T-REx host cell line was used to generate cell lysates for the co-condensation experiments.

### siRNA treatments

Cells were transfected with a non-targeting siRNA pool (ON-TARGETplus non-targeting control pool, Dharmacon) or a custom TDP-43-specific siRNA (sequence: 5’-GGCUCAAGCAUGGAUUCUAAGUCUU-3’, Biomers) at a final concentration of 20 nM using Lipofectamine RNAiMax (Invitrogen) according to manufacturer’s instructions. Cells were analyzed at 72 hours post-transfection.

### Heat stress treatments

To induce heat shock (HS), cells were placed in a humidified incubator at 43°C and 5% CO_2_ for 1 hour. Afterwards, cells underwent recovery in normal growth conditions (37°C, 5% CO_2_) for 3 hours.

### Immunofluorescence staining

Cells were fixed using 3.7% formaldehyde in PBS for 8 minutes and permeabilized with 0.5% Triton X-100 in PBS for 5 minutes. Blocking to prevent unspecific binding of antibodies was done for 30 minutes at RT using blocking buffer (5% donkey serum in PBS with 0.1% Tween-20). Cells were incubated with primary antibodies diluted in the blocking buffer for 1.5 hours (anti-myc mouse 9E10 antibody diluted 1:100, IMB Protein Production core facility), followed by 3 washes using PBS with 0.1% Tween-20. Secondary antibody (anti-mouse conjugated to Alexa Fluor 488, Invitrogen, A21202) was applied in blocking buffer for 1 hour at RT, followed by another 3 washes in PBS with 0.1% Tween-20. Nuclei were stained with 0.5 μg/ml DAPI in PBS (Sigma-Aldrich, D9542) for 5 minutes.

### Preparation of cell lysates for western blotting and full proteome analysis

Cells were washed with ice-cold PBS and collected using a cell scraper. Collected material was centrifuged at 500 g for 5 minutes at 4°C, and the pellets were lysed with ice-cold RIPA buffer (50 mM Tris-HCl pH 8, 150 mM NaCl, 1% NP-40, 0.5% sodium deoxycholate, 0.1% SDS) for 15 minutes on ice. Cells were then centrifuged at 15 000 g for 15 minutes at 4°C, and the supernatant containing cellular proteins was used to quantify protein levels using the BCA assay.

### SDS-PAGE and western blotting

For SDS-PAGE, lysates were supplemented with Laemmli buffer (62.5 mM Tris pH 6.8, 10% glycerol, 2% SDS, 0.025% bromophenol blue, 1% β-mercaptoethanol) and boiled for 5 minutes at 95°C. Proteins were separated on a 10% SDS-PAGE gel, followed by a wet transfer onto a 0.2 μm nitrocellulose membrane (Amersham Protran, Cytiva). Blocking was done in 5% milk in TBST for 30 minutes, followed by an overnight incubation with the indicated primary antibody. Afterwards, the membrane was washed 3 times with TBST and incubated with a species-specific secondary antibody conjugated to an IRDye 800CW or 680RD (Li-COR) for 1 hour at RT. Membrane visualization was done using the Odyssey M system (Li-COR). Quantification of the western blot signal was done using the Image Studio Lite (LI-COR). Antibodies used for western blotting: rabbit anti-TDP-43 (Proteintech, 80001-RR), mouse anti-TDP-43 (Proteintech, 60019-2-Ig), rabbit anti-GAPDH (Proteintech 60004-1-Ig), rabbit anti-UPF1 (Sigma-Aldrich, HPA019587).

### Confocal fluorescence microscopy: imaging of immunofluorescence-stained cells

To assess TDP-43 nuclear foci counts, cells were seeded onto 96-well plates and after immunofluorescence staining, were imaged using the Revvity Opera Phenix high content confocal microscope (IMB Mainz microscopy core facility). Imaging was performed using the 63x water immersion objective, NA 1.15, binning 2. The following settings were applied for each imaging channel: DAPI (excitation at 405 nm, emission range: 435 - 480nm, time set to 100 ms, power set to 50%), GFP (excitation at 488 nm, emission: 500 - 550 nm, time set to 200 ms, power set to 100%).

Quantification of nuclear TDP-43 foci was performed using the Revvity Harmony analysis software. Cell nuclei were segmented based on the nuclear EGFP- or myc-TDP-43 signal to ensure all cells expressing enough of the protein of interest are included. Cells on image borders were omitted. Spots were identified using the spots segmentation function, which was tuned to ensure segmentation of the foci that specifically represent TDP-43. For cells that had undergone heat shock (1 hour HS + 3 hours recovery), this parameter was adjusted to detect brighter foci formed by TDP-43.

### Fluorescence recovery after photobleaching (FRAP)

All FRAP experiments were performed using the STELLARIS 8 FALCON (Leica Microsystems) confocal microscope equipped with the LAS X imaging platform. The images were obtained using a HC PL APO CS2 63x/1.4 oil objective for all experiments.

#### FRAP of in vitro-formed TDP-43 condensates

For FRAP experiments, TDP-43 condensates were formed from fluorescently labelled TDP-43-MBP-His_6_ protein as described in the “Phase separation assay with purified proteins” section. PS-deficient TDP-43 variants (3W, ΔCR, 12D) were analyzed at 40 μM protein concentration, and solid-like TDP-43 variants (5R, 12A, G376V) were analyzed at 15 μM protein concentration. As a control, TDP-43 WT condensates were analyzed in the same concentration regime. Bleaching experiments were performed between 15 and 30 minutes after TEV cleavage for solid-like variants, and between 15 and 45 minutes after TEV cleavage for PS-deficient variants.

For pre-bleach and post-bleach imaging, 476 nm excitation line from White Light Laser and HyD S3 detector in counting mode (emission band 500 - 739 nm) were used. Half-bleaches were done within the selected region of interest in “zoom in” mode using an OPSL488 laser at 488 nm excitation and laser intensity of 25%. Five pre-bleach frames were acquired, followed by photobleaching, then 50 post-bleach frames in “minimize mode”, and 30 additional frames at 2-second intervals. The pixel size for all images corresponds to 60.18 nm (scan format 512 x 512 px).

#### FRAP of EGFP-TDP-43-containing nuclear foci in T-REx Flp-In HeLa cells

Cells were grown on 8-well chambered slides (Ibidi, 80826) and subjected to TDP-43 knockdown using siRNA (72 hours total), doxycycline induction (24 hours total) and, where appropriate, heat stress treatment as described above. FRAP experiments were conducted on live cells at 37°C and 5% CO_2_.

For pre-bleach and post-bleach imaging, 476 nm excitation line from White Light Laser and HyD S3 detector in counting mode (emission band 500 - 550 nm) were used. Photobleaching was done using the 488 nm excitation line from White Light Laser at intensity of 100%. For non-stressed cells, regions of interest were photobleached using the “zoom in” mode, while for heat-shocked cells a point bleach was used. Five pre-bleach frames were acquired, followed by photobleaching, then 40 post-bleach frames in “minimize mode”, and 10 additional frames at 2-second intervals. The pixel size for all images corresponds to 72.2 nm (scan format 512 x 512 px).

#### Analysis of FRAP data

All images were prepared using Fiji distribution of ImageJ^74^. Mean fluorescence intensity was quantified in three regions of interest (ROI1 – bleached region, ROI2 – whole condensate/cell nucleus, ROI3 – background). Further data normalization (double) and quantification were done using the easyFRAP software^75^. Obtained curves were graphed using GraphPad Prism 9, and significance was determined based on area under the curve (AUC) values for each curve obtained.

### Affinity Purification coupled with Mass Spectrometry (AP-MS)

#### Antibody crosslinking to protein G Sepharose for immunoprecipitation

Prior to immunoprecipitation experiments, crosslinking of the mouse anti-myc antibody (clone 9E10, IMB Protein Production core facility) to protein G Sepharose was performed according to Gersten & Marcgalonis^76^.

Beads were equilibrated in binding buffer (20 mM HEPES pH 7.5, 150 mM NaCl) and incubated with the anti-myc antibody for 1 hour at RT in a 2:1 antibody:beads ratio. Beads were extensively washed with borate buffer pH 9.0, and crosslinked using 20 mM dimethyl pimelimidate (DMP) for 30 minutes at RT. The reaction was quenched using 0.2 M ethanolamine, and the beads were washed and stored in IP buffer (50 mM Tris pH 8.0, 150 mM NaCl).

#### Affinity purification of myc-tagged TDP-43 and MS preparation

HeLa Flp-In T-REx cells expressing a single copy of N-terminally myc-tagged TDP-43 WT or variants were subjected to TDP-43 knockdown (KD), and 48 hours later, the expression of myc-TDP-43 variants was induced with doxycycline for 24 hours as described in “siRNA treatments” and “Stable cell line generation and cell culture” sections.

For each affinity purification experiment, cells were harvested and processed for lysis and affinity purification of the bait protein as previously described by *Uliana et al.*^77^. As a control, non-induced HeLa Flp-In T-REx myc-TDP-43 WT cells with a TDP-43 knockdown were processed in parallel. The experiment was performed with three independent biological replicates. Cell pellets were collected, frozen, and subsequently lysed in 1500 µl of lysis buffer (0.5% NP-40, 50 mM Tris-HCl (pH 8.0), 150 mM NaCl, and 2.5 mM MgCl_2_, supplemented with protease inhibitors for 20 minutes on ice. Lysates were sonicated using the Bioruptor Pico (Diagenode) for 1 cycle of 45 seconds “on” and 45 seconds “off”. Lysates were then treated with Benzonase (50 U/mL) for 20 minutes at 10°C. Insoluble material was removed by centrifugation at 20 000 g for 20 minutes at 4°C. The cleared supernatant was incubated overnight at 4°C with 50 μl of beads conjugated to 100 µg of an anti-myc antibody. Beads were then extensively washed with lysis buffer with and without detergent (total wash volume: 5 ml). Beads with bound proteins were transferred to a 10 kDa MWCO centrifugal filter unit (Vivacon 500, Sartorius) and processed according to the FASP protocol^78^. Beads in solution were centrifuged at 8 000 g until dry and the proteins bound were denatured and reduced in 8 M urea and 5 mM TCEP prepared in 50 mM ammonium bicarbonate for 30 minutes and alkylated with 10 mM iodoacetamide for 30 minutes in the dark. Samples were washed three times with 25 mM ammonium bicarbonate and digested with 1 µg sequencing-grade trypsin (Promega) for 16 hours at 37°C. Proteolysis was quenched by the addition of 5% formic acid and peptides were recovered by centrifugation, purified through C18 StageTip columns according to the manufacturer’s instructions (Affinisep), dried in a vacuum concentrator, and resuspended in 20 µl of 0.1% formic acid and 2% acetonitrile prior to mass spectrometry analysis.

#### MS acquisition

LC-MS/MS analysis was performed on a Q Exactive Plus Hybrid Quadrupole-Orbitrap Mass Spectrometer (Thermo Fisher) coupled to Dionex Ultimate 3000 RSLCnano System liquid chromatography system. Peptides were loaded on a commercial trap column (μ-Precolumn C18 PepMap100, C18, 300 μm I.D., 5 μm particle size) and separated using a reverse phase column Thermo PepMap100 C18 (50 cm length, 75 μm inner diameter, 35 μm particle size) across a gradient from 8 to 25% in 120 minutes (buffer A: 0.1% (v/v) formic acid; buffer B: 0.1% (v/v) formic acid, 95% (v/v) acetonitrile). The data-dependent acquisition (DDA) mode was set to perform one MS1 scan followed by a maximum of 20 scans for the top 20 most intense peptides with MS1 scans (R = 70 000 at 400 m/z, AGC = 3 × 10^6^ and maximum IT = 64 ms), HCD fragmentation (NCE = 28%), isolation windows (1.4 m/z) and MS2 scans (R = 17 500 at 400 m/z, AGC = 1 × 10^5^ and maximum IT = 55 ms). A minimum AGC target of 2 000 and dynamic exclusion of 20s was applied and charge states lower than two and higher than seven were rejected for the isolation.

#### Data analysis

Acquired spectra were processed using MaxQuant (v. 2.4.13.0) with the integrated Andromeda search engine^79^ and searched against the human UniProt reference proteome (downloaded on 27.09.2024; 20 427 entries), supplemented with a database of common contaminants. Database searches were performed with full tryptic specificity, allowing up to two missed cleavages. Carbamidomethylation of cysteines was set as a fixed modification, while oxidation of methionine and protein N-terminal acetylation were included as variable modifications. The “match between runs” feature was enabled with a retention time window of ±0.4 minutes. The precursor and fragment mass tolerances were set to 20 ppm. Peptide-spectrum match (PSM) and protein FDR were controlled at 1%. Protein abundances were determined using the MaxLFQ algorithm, and only proteins identified with at least two peptides, including at least one unique peptide, were considered for quantification. In the dataset 2 560 proteins were identified, filtered to 2 189 (proteins must be identified in triplicate in at least one condition). Protein intensities were log_2_-transformed and median-normalized across samples. Missing values (∼15%) were imputed by random sampling from a normal distribution generated for each protein based on the observed intensity distribution across samples. For each protein, the imputation distribution was centered at the minimum observed log_2_ intensity minus 3 and assigned a standard deviation equal to the standard deviation of the observed values. Statistical analysis was performed using analysis of variance (ANOVA) comparing all affinity purifications to the control condition. TDP-43 interactors were defined as proteins that showed significant enrichment in at least one purification with a log_2_FC (fold change) > 1 and *p*value (BH adjusted) < 0.05 (226 proteins, excluding TDP-43 itself). The abundance of TDP-43 interactors was normalized for the abundance of the bait and compared to their abundance in the TDP-43 WT condition. For cluster definition, the average of normalized abundance across replicates was calculated for each interactor and z-score–transformed. Interactors were subsequently clustered based on the similarity of their abundance profiles using Pearson correlation and the interactors were displayed in the network (Cytoscape v. 5.0.253) based on the interaction retrieved from protein-protein interactions repository (BioGRID v. 5.0.253, considering only physical interactions). GO characterization was performed using EnrichR v.3.4 using as background the human proteome, and only terms comprising at least 5 genes for CC and MF and 10 for BP were considered. To reduce redundancy, GO terms passing a *p*value threshold of 0.05 were subjected to semantic similarity reduction using the reduceSimMatrix approach, which identifies representative parent terms based on a similarity threshold of 0.5. The entire dataset, including raw data, generated tables are available in the PRIDE repository (PRIDE: PXD074566)

#### Identification of high-confidence interactors and generation of protein-protein interactions network

A curated analysis of reported interactors was performed. This includes previously published TDP-43 interactomics datasets with appropriate controls^20,25,31–33,80–82^, integrated with protein-protein interactions annotated in BioGRID (v. 5.0.253 downloaded in January 2026, including only the following interactions types “physical association”, “direct interaction”, “colocalization”)^83^ and manually curated interactions from the literature^84^. Redundancy between datasets was manually removed so that each interaction was reported only once. From the resulting list (2 048 interactors), high-confidence interactors (HCI) were defined as those identified in more than three independent experiments (230 HCI).

The TDP-43 protein–protein interaction network was generated from correlations among interactor profiles. Network clusters were characterized by Gene Ontology cellular component (CC) terms, and proteins were assigned to modules based on annotated interactions from BioGRID (v. 5.0.253) using a guilt-by-association approach and Gene Ontology CC terms.

#### Characterization of TDP-43 interactome features

For functional association of proteins with TDP-43 analysis, protein-protein interaction data were obtained from STRING (v.12, “9606.protein.links.v12.0.onlyAB.txt”)^85^, and interaction confidence scores were converted into edge weight for network analysis. Edge costs were defined as the negative natural logarithm of the confidence score, allowing shortest-path computation to favor high-confidence interactions. For each protein (18 841) the shortest path to TDP-43 was calculated and a score was computed as the product of the confidence values of all edges in the path.

The enrichment for specific features within the TDP-43 interactome was assessed by comparing interactors to a random sampled control group matching the size of the TDP-43 interactors defined as binding in a phase separation-dependent manner (N = 195). Continuous protein features like AlphaFold2 pLDDT confidence scores (downloaded in January 2025)^65^ and predicted secondary structure content, were calculated for each protein and compared between groups by analyzing their distributions using an unpaired non-parametric Wilcoxon test. Enrichment of functional annotations was evaluated using hypergeometric tests. RG, RGG and extended RGG consensus motifs (G(0–3)X(0–1)RG(1– 2)X(0–5)G(0–2)X(0–1)RG(1–2))^86^ were searched across protein sequences from the human proteome to identify motif-containing proteins. Annotation of phase-separation–associated proteins was obtained from PhaSepDB (v. 2.1, LLPS-related entries only; downloaded in December 2022)^87^ and from CD-CODE v. 2 (including only proteins experimentally observed in biomolecular condensates in at least three independent studies; downloaded in January 2025)^88^.

### Co-condensation experiment

#### Sample preparation

HeLa total standardized cell lysates were prepared freshly for each experiment. Parental HeLa Flp-In T-REx host cells were lysed in a mild buffer (50 mM Tris pH 7.5, 150 mM NaCl, 0.5% NP-40, 2.5 mM MgCl_2_, supplemented with a protease inhibitor cocktail (Sigma-Aldrich, PIC0004), phosphatase inhibitors (PhosSTOP, Roche, 04906837001) and Benzonase (50 U/ml)) for 30 minutes at 10°C. Obtained lysate was then centrifuged for 20 min at 20 000 g and protein concentration was measured by BCA. Phase separation reactions were set up by the addition of TDP-43-MBP-His_6_ variants (10 μM final concentration), HeLa cell lysate (2 mg/ml final concentration) and TEV protease (5 µM final concentration) in the following buffer: 50 mM Tris pH 7.5; NaCl 150 mM, 2 mM DTT. Reactions were incubated for 45 minutes at RT.

#### Microscopy analysis

Phase separation reactions were set up as described above using fluorescently labelled TDP43-MBP-His_6_ and imaged in 384-well plates as described in the “Confocal microscopy: imaging of fluorescently labeled TDP-43 condensates” section. Quantification was performed using the Revvity Harmony analysis software. Individual condensates were segmented using the spots segmentation, and condensates on the image borders were omitted.

#### Western blotting analysis

For western blotting and subsequent detection of TDP-43 and UPF1 levels, samples were centrifuged to separate the pellet and supernatant. Proteins were subsequently separated using SDS-PAGE, followed by western blotting using mouse anti-TDP-43 (diluted 1:2 000, Proteintech, 60019-2-Ig) and rabbit anti-UPF1 (diluted 1:1 000, Sigma-Aldrich, HPA019587) antibodies. For TDP-43 signal detection, equal amounts of pellet and supernatant were loaded and quantified. For UPF1 signal detection, pellet was overrepresented 5 times compared to supernatant to increase the detection sensitivity.

#### Sample preparation for MS analysis

Lysate was collected before the addition of TEV protease and supernatant (SN) was collected after the addition of TEV protease. After a 45-minute incubation and centrifugation, they were subjected to sample processing. For quality control, 0.05 µg of BSA was spiked into each sample as an internal standard. Samples were processed using the SP3 protocol^89^. Proteins were reduced (5 mM DTT), alkylated (15 mM iodoacetamide in the dark), and quenched (5 mM DTT). Proteins were digested overnight with trypsin at 37°C. After acidification with formic acid, peptides were purified using C18 StageTips for solid-phase extraction.

#### MS acquisition

LC-MS/MS analysis was performed on an Orbitrap Astral mass spectrometer (Thermo Fisher) coupled to a Vanquish Neo liquid chromatography system (Thermo Fisher). Peptides were separated using a 45 cm reverse phase column (inner diameter: 75 μm; ReproSil-Pur 120 C18-AQ 1.9 μm silica particles, Dr. Maisch GmbH) across a linear gradient from 5 to 40% in 31 minutes (buffer A: 0.1% (v/v) formic acid; buffer B: 0.1% (v/v) formic acid, 80% (v/v) acetonitrile). The mass spectrometer was operated in data-independent acquisition (DIA) with the following parameters: one full FTMS scan in Orbitrap (350 - 1 050 m/z) at 120 000 resolution, 20 ms injection time, and AGC target (%) = 300, followed by 175 fixed windows of 4 Da from 350 to 1 050 m/z, 5 ms injection time, and AGC target (%) = 500 (Astral detector). Precursor ions were fragmented with HCD, Normalized Collision Energy 26%, isolation window 4 m/z.

#### Analysis of the MS data

Acquired DIA spectra were searched using FragPipe computational platform v. 23.0 ^90^, Philosopher v. 5.1.1^91^, DIA-NN v. 1.8.2^92^. Peptide identification from MS/MS spectra was performed with MSFragger search engine against the Human Proteome reference dataset extended with contaminant list (http://www.uniprot.org/, downloaded in July 2025, 20 453 proteins) and reverse decoy sequences. Mass tolerance was set to 10 ppm for precursor and for fragment ions. The search parameters were set to include fully tryptic peptides, allowing for up to one missed cleavage site, carbamidomethylation on cysteine residues as a static modification and up to three variable modifications including methionine oxidation and N-terminal acetylation. Peptide and protein inference was FDR controlled at 1% by PeptideProphet and ProteinProphet respectively. The generated library included entries for 159 647 precursors and 7 487 protein groups. For the DIA analysis, extraction of quantitative data was performed with DIA-NN v. 1.8.2 beta 8 with a maximum FDR of 1% and quantification strategy was set to “Robust LC”. Precursor abundance values were extracted from the file “report.tsv” and only proteotypic precursors with intensities > 0 and a PEP < 0.01 were log_2_-transformed and median-normalized for downstream analysis. For each protein quantified with at least two precursors in every condition, precursor-level filtering was applied. Within each condition, precursor intensities were centered on the median, and precursor variability was quantified as the mean squared error (MSE) across all conditions. Precursors were ranked by their average MSE, and the 50% most variable precursors were excluded, while ensuring that at least two precursors were retained per protein. For the quantification of TDP-43 precursors, the peptides containing variant-specific modified residues were discarded. For each condition, the top 5 most abundant filtered precursors were selected, and their median intensity was used to estimate protein abundance.

To characterize the abundance of depleted TDP-43 in the supernatant, values were normalized to the WT TDP-43 -TEV condition, converted to linear scale, and expressed as percentage relative abundance.

With the stringent filtering criteria applied to the dataset, no imputation of missing values was performed. Differential analysis for the abundance of 5 676 proteins (log_2_-transformed values) was performed with limma package^93^. A linear model without intercept was fitted to protein abundance values across 32 samples representing eight experimental conditions (N = 4 independent replicates per condition). Moderated t-tests were applied using empirical Bayes variance shrinkage, and contrasts were defined to compare each TDP-43 variant against the TDP-43 WT + TEV condition, *p*values were adjusted according to Benjamini-Hochberg method. Proteins were considered significantly downregulated if they met the thresholds: log_2_FC < −0.3 and adjusted *p*value < 0.1. To evaluate condition-specific changes, the protein abundance for each protein across mutant conditions was transformed into z-scores. The entire dataset, including raw data, generated tables are available in the PRIDE repository (PRIDE: PXD076450)

### Dose-response co-condensation experiment

#### Sample preparation for MS analysis

For the dose-response co-condensation profiling, increasing concentrations of WT and 5R TDP-43 (0, 1, 2, 5, 10 µM) were spiked into a standardized lysate (2 mg/ml) and 10 µM of TEV protease was added to the lysate to remove the MBP solubility tag. After 30 minutes at RT the lysates were subjected to centrifugation, and 10 µl (∼20 µg) supernatant was subjected to proteomics preparation. For quality control, 0.05 µg of BSA was spiked into each sample as an internal standard and samples were processed using the SP3 protocol^89^ as described for the co-condensation experiment.

#### MS acquisition

LC-MS/MS analysis was performed on an Orbitrap Astral mass spectrometer (Thermo Fisher) coupled to a Vanquish Neo liquid chromatography system (Thermo Fisher). Peptides were separated using a 45 cm reverse phase column (inner diameter: 75 μm; ReproSil-Pur 120 C18-AQ 1.9 μm silica particles, Dr. Maisch GmbH) across a linear gradient from 5 to 40% in 31 minutes (buffer A: 0.1% (v/v) formic acid; buffer B: 0.1% (v/v) formic acid, 80% (v/v) acetonitrile). The mass spectrometer was operated in the data-independent acquisition (DIA) mode with the following parameters: one full FTMS scan in Orbitrap (350 – 1 050 m/z) at 120 000 resolution, 20 ms injection time, and AGC target (%) = 300, followed by 175 fixed windows of 4 Da from 350 to 1 050 m/z, 5 ms injection time, and AGC target (%)= 500 (Astral detector). Precursor ions were fragmented with HCD, Normalized Collision Energy 26%, isolation window 4 m/z.

#### Analysis of MS data

Acquired DIA spectra were searched using FragPipe computational platform v. 23.0^90^, Philosopher v. 5.1.1^91^, DIA-NN v. 1.8.2^92^. Peptide identification from MS/MS spectra was performed with MSFragger search engine against the Human Proteome reference dataset extended with contaminant list (http://www.uniprot.org/, downloaded in June 2025, 20 453 proteins) and reverse decoy sequences. Mass tolerance was set to 10 ppm for precursor and for fragment ions. The search parameters were set to include fully tryptic peptides, allowing for up to one missed cleavage site, carbamidomethylation on cysteine residues as a static modification and up to three variable modifications including methionine oxidation and N-terminal acetylation. Peptide and protein inference was FDR controlled at 1% by PeptideProphet and ProteinProphet respectively. The generated library included entries for 175 366 precursors and 7 946 protein groups. For the DIA analysis, extraction of quantitative data was performed with DIA-NN v. 1.8.2 beta 8 with a maximum FDR of 1% and the dose-response quantification strategy was set to “Robust LC”. Precursor abundance values were extracted from the file “report.tsv” and only proteotypic precursors with intensities > 0 and a PEP < 0.01 were log_2_-transformed and median-normalized for downstream analysis. For each protein quantified with at least two precursors in every condition, precursor-level filtering was applied to reduce technical variability. Within each condition, precursor intensities were centered on the median, and precursor variability was quantified as the mean squared error (MSE) across all conditions. Precursors were ranked by their average MSE, and the 50% most variable precursors were excluded, while ensuring that at least two precursors were retained per protein. For each condition, protein abundance was calculated from the median of the top five most intense precursors. With the stringent filtering criteria applied to the dataset, no imputation of missing values was performed. For all quantified proteins (N = 5 969), dose-response curve fitting was performed at the protein level using the protti R package^94^. A four-parameter logistic model was applied to protein abundance values (log_2_ scale) as a function of TDP-43 spike-in concentration. Proteins co-precipitating with TDP-43 5R were identified by applying a series of filters to dose–response curve parameters. For each protein, values were retained only if the Hill coefficient was > 0.7, the adjusted ANOVA *p*value was < 0.1, and the correlation coefficient was > 0.7. The area under the curve (AUC) was then compared between wild-type (WT) and TDP-43 5R samples. 403 proteins were classified as co-precipitating with TDP-43 5R based on the following two criteria: i) passed the filtering criteria in 5R but not WT, ii) had an area under the curve ratio 5R versus WT < 1 (Supplementary Fig. 5C). The entire dataset, including raw data and generated tables are available in the PRIDE repository (PRIDE: PXD076467).

#### Characterization of proteins co-condensing with TDP-43 5R

The characterization of features of co-condensing proteins (functional association to TDP-43 and enrichment for proteins associated to phase separation according PhaSepDB and CD-CODE) was performed as described in the “Characterization of TDP-43 interactome features” section.

#### Analysis of molecular grammar

The sequence analysis was performed using localCIDER^95^ following the methodology described by Han et al.^96^. A total of 128 sequence-based parameters were computed for each protein, including 44 global parameters, 14 local parameters (for which maximum, mean, median, and standard deviation were calculated), and 7 patch-based parameters (basic, acidic, charged, polar, aliphatic, aromatic, and proline). For local parameters, a sliding window of 5 residues was used, while patches were defined as contiguous runs of three or more identical or chemically similar residues. Contiguous stretches of residues separated by fewer than two residues were merged, and only patches with at least 5 residues were considered. Sequence parameters were calculated across the human proteome as a reference, and values were z-score normalized relative to the human distribution. For proteins co-precipitating with the TDP-43 5R variant or identified as PS-dependent interactors, an average z-score was computed and compared to the reference proteome. Positive z-scores indicate enrichment, whereas negative values indicate depletion. Statistical significance of enrichment or depletion was assessed using Welch’s two-sample t-test. Features significantly identified in at least one dataset (TDP-43 PS-dependent interactome or coprecipitating with 5R, *p*value BH adjusted < 0.01) and with a coherent enrichment or depletion between the two conditions are shown in the heatmap.

### RNA-Sequencing

#### RNA extraction

Prior to RNA sequencing, the different stable HeLa Flp-In T-REx myc-TDP-43 cell lines were grown and subjected to siRNA treatment (TDP-43-specific or non-targeting control) for 72 hours. Doxycycline induction was performed 18 hours after the siRNA transfection to minimize potential secondary effects caused by long-term TDP-43 depletion. Cells were washed with ice-cold PBS, collected using cell scrapers and centrifuged at 500 g for 5 minutes. Total RNA was isolated from cell pellets using the PureLink RNA Mini-Kit (Invitrogen) according to manufacturer’s instructions, including the DNAse treatment step.

#### Library preparation and RNA sequencing

NGS library preparation was performed with Illumina’s Stranded mRNA Prep Ligation Kit following Stranded mRNA Prep Ligation Reference Guide (April 2021) (Document 1000000124518 v02). Libraries were prepared with a starting amount of 1 000 ng and amplified in 8 PCR cycles. Two post-PCR purification steps were performed to exclude residual primer and adapter dimers. Libraries were profiled in a D5000 Screen Tape on a tape Station 4200 (Agilent technologies) and quantified using the Qubit 1x dsDNA HS Assay Kit, in a QubitFlex Fluorometer (Invitrogen by Thermo Fisher Scientific). All 63 samples were pooled in equimolar ratio and sequenced on 2 NextSeq2000 P4 (100 cycles) flow cells, in single read mode for 1x 116 cycles plus 2 × 10 cycles for the dual index read and 1 dark cycle upfront R1.

### RNA sequencing data analysis

Basic quality controls were done using FastQC (v. 0.12.1) (https://www.bioinformatics.babraham.ac.uk/projects/fastqc/). Prior to mapping, possibly remaining adapter sequences were trimmed using Cutadapt (v. 4.4)^97^ including a NextSeq-specific quality trimming at the 3’ end of reads (--nextseq-trim = 20). A minimal overlap of 5 nt between read and adapter was required and only reads with a length of at least 30 nt after trimming (--minimum-length 30) were kept for further analysis.

Reads were mapped using STAR (v. 2.7.10b)^98^ allowing up to 4% of the bases to be mismatched (--outFilterMismatchNoverReadLmax 0.04 --outFilterMismatchNmax 999) and with a splice junction overhang of 115 nt (--sjdbOverhang 115). Genome assembly and annotation of GENCODE^99^ release 43 were used during mapping. Secondary hits were removed using Samtools (v. 1.17)^100^.

Uniquely mapped exonic reads per gene were counted using featureCounts of the Subread tool suite (v. 2.0.0)^101^ with parameters --donotsort -t exon -s2. For each mutant, genes differentially expressed between conditions were detected using the DESeq2 R package (v. 1.42.0)^102^. The design formula ∼ replicate + condition was used to account for paired replicates and test the effect of the condition. An independent filtering was applied and a significance cutoff of 0.01 was used.

#### Alternative splicing analysis

Alternative splicing events were detected using rMATS (v. 4.1.2)^103^. For each mutant, different conditions were compared pairwise. When running rMATS, the detection of novel splice sites was enabled (--novelSS), clipped alignments were allowed to be used (--allow-clipping) and parameters for reverse-stranded single-end reads were set (--libType fr-firststrand -t single). The read length was set to 116 (--readLength 116), but variable read lengths were allowed (--variable-read-length). The same annotation as during mapping (GENCODE release 43) was used when running rMATS.

After running rMATS, skipped exon events from JCEC results table were filtered based on the following criteria: average counts (of combined inclusion IJC and skipping SJC supporting reads) per condition >= 10, false discovery rate (FDR) < 0.05, and absolute difference in percent spliced in (|ΔPSI|) > 0.1.

#### Motif and RNA binding analysis in alternative splicing flanking sequences

Genomic coordinates of filtered splicing events were extracted and then BSgenome (v. 1.70.2) was used to retrieve sequences from human reference genome (hg38) spanning 500 nucleotides flanking the alternatively spliced exons on both sides.

Motif enrichment analysis was performed using AME (Analysis of Motif Enrichment) ^104^ from the MEME suite, utilizing 6-mer motifs or motifs from the CISBP-RNA database^105^ (CISBP-RNA *Homo sapiens*, DNA-encoded) using the sequences spanning 500 nucleotides flanking not significant alternatively spliced exons (FDR > 0.05) as a background control. Enrichment significance (-log_10_ *p*values BH adjusted) was calculated by hypergeometric test. To quantify motif diversity, Shannon entropy was calculated for each condition based on the adjusted *p*value enrichment (considering only motifs with -log_10_*p*value adj. < 0.05). *p*values were transformed to form a probability distribution across motifs and Shannon entropy was then calculated as 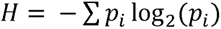. Condition specificity was defined as 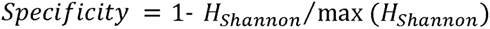, providing a measure of motif specificity.

Protein-RNA binding sites were obtained from publicly available eCLIP datasets provided through the ENCORE Matrix of the ENCODE platform^106^ (experiments: TDP-43: ENCSR720BJU, ENCSR584TCR, ENCSR187VEQ; DDX3X: ENCSR930BZL, ENCSR648LAH; EFTUD2: ENCSR844RVX, ENCSR527DXF; ELAVL1: ENCSR090LNQ; FASTKD2: ENCSR887FHF, ENCSR023UHL; FUBP3: ENCSR486YGP; HNRNPC: ENCSR249ROI, ENCSR550DVK; HNRNPK: ENCSR953ZOA, ENCSR268ETU; ENCSR828ZID; HNRNPL: ENCSR795CAI, ENCSR724RDN; HNRNPM: ENCSR412NOW, ENCSR267UCX; HNRNPU: ENCSR520BZQ, ENCSR240MVJ; MATR: ENCSR440SUX, ENCSR290VLT; PRPF8: ENCSR534YOI, ENCSR121NVA; UPF1: ENCSR456ASB, ENCSR539BEV)(https://www.encodeproject.org/encore-matrix/?type=Experiment&status=released&internal_tags=ENCORE).

File bed or bigBed files from eCLIP datasets were used to extract binding coordinates and overlapped alternative splicing flanking sequences (+/− 500nt). Regions were considered overlapping if at least 10 nucleotides were shared.

#### Identification of TDP-43-dependent and phase separation-dependent alternative splicing events

TDP-43-dependent alternative splicing events were defined as those meeting the following criteria: i) average junction counts (IJC and SJC) ≥ 10 ii) significant changes between KD versus NTC and between Rescue versus KD (FDR < 0.05), iii) have opposite ΔPSI values between KD versus NTC and between rescue versus KD. In total 425 events in 242 genes were identified and classified as either: (i) “rescue of exon inclusion” (185 events), where inclusion levels decreased upon knockdown and increased upon rescue, or (ii) “rescue of exon skipping” (240 events), where inclusion levels increased upon knockdown and decreased upon rescue. For TDP-43-dependent events, the ΔPSI observed in the rescue versus KD condition for WT was subtracted from the corresponding ΔPSI for each variant to illustrate the differences in splicing rescue efficiency of each variant compared to WT (ΔΔPSI). PS-dependent events were defined as those in which the variability within each group (PS-deficient and solid-like) is lower compared to the global variability (total: 75 events).

Sashimi plots were generated using the “rmats2sashimi” pipeline which creates sashimi plot visualizations of the rMATS outputs. The pipeline for Sashimi visualization is available under: https://github.com/Xinglab/rmats2sashimiplot. Since rMATS excludes multi-mapping reads, BAM files were filtered to retain only uniquely mapped reads (-q 255) and subsequently sorted using Samtools (v. 1.17)^107^. The resulting BAM files and events of interest identified earlier using rMATS were used for the visualization.

#### Differential gene expression analysis

The results from the differential gene expression analysis described above were combined, and genes with at least two concordant significant changes in the rescue versus KD condition (|log_2_FC|> 0.5 and *p*value BH adjusted < 0.05) within each group (solid-like and PS-deficient) were selected (447 genes in total), PS-dependent genes were defined as those in which the log_2_FC variability within each group (PS-deficient and solid-like) is lower compared to the global variability (total: 87 genes). Gene abundance levels (log2FC) were converted to z-scores to enhance visualization.

Gene Ontology (GO) enrichment analysis was conducted for PS-dependent up- and downregulated genes using the clusterProfiler package v. 4.10.1 with default background, focusing on Biological Process (BP) terms with q-values < 0.01 and *p*value < 0.05. Redundant GO terms were reduced using the rrvgo package v. 1.14.2, clustering terms based on semantic similarity using the Wang method (threshold = 0.8). The mean log_2_FC (z-score normalized) of genes associated with each GO term was used to represent changes between solid-like and PS-deficient groups.

#### UPF1 gene set enrichment analysis

Rescue samples of solid-like and PS-deficient mutants were compared and ranked based on the Wald statistic as calculated by DESeq2 (v. 1.42.0). Low count genes with a read count < 10 in more than 9 of the 18 rescue samples were removed prior to the analysis. The same analysis was performed for the comparison between rescue vesus KD in WT condition, with removal of low count genes with a read count < 10 in more than three of the six analyzed conditions. A gene set enrichment analysis of a set of UPF1-dependent genes was performed using clusterProfiler (v. 4.10.1). A set of 310 UPF1-dependent genes was composed based on genes identified in at least two of the three published datasets (based on HeLa or HEK293 cell lines) *Imamachi et al.* ^108^ (data source: table S4), *Fritz et al.* ^109^ (data source: table EV2_Supplementary; siUPF1total.vs.NT.log2.fold.change > 1 & FDR < 0.05), and *Colombo et al.* ^110^ (data source: Table S2; UPF1_FDR < 0.01). Gene symbols were mapped to GENCODE release 43 gene identifiers, discarding genes with multiple Ensembl assignments. For genes that could not be mapped directly, updated HGNC gene symbols were retrieved using the HGNC Multi-Symbol Checker and remapped to GENCODE annotations. A final set of 273 genes were identified in the data and tested for enrichment.

### Full proteome characterization

#### Sample preparation

HeLa Flp-In T-REx myc-TDP-43 cell lines (WT or variants) were subjected to TDP-43 KD knockdown (KD) for 72 hours. Doxycycline induction was performed 18 hours after the siRNA transfection.

Cells were harvested and lysed as described in the “Preparation of cell lysates for western blotting and full proteome analysis” section, and the supernatant containing cellular proteins was used to quantify protein levels using a BCA assay. 20 µg of the total proteome were processed using the SP3 protocol^89^. Proteins were reduced (5 mM DTT), alkylated (15 mM iodoacetamide in the dark), and subsequently quenched (5 mM DTT). Proteins were digested overnight with trypsin at 37°C. After acidification with formic acid, peptides were purified using C18 StageTips for solid-phase extraction.

#### MS acquisition

LC-MS/MS analysis was performed on an Orbitrap Astral mass spectrometer (Thermo Fisher) coupled to a Vanquish Neo liquid chromatography system (Thermo Fisher). Peptides were separated using a 45 cm reverse phase column (inner diameter: 75 μm; ReproSil-Pur 120 C18-AQ 1.9 μm silica particles, Dr. Maisch GmbH) across a linear gradient from 5 to 40% in 31 minutes (buffer A: 0.1% (v/v) formic acid; buffer B: 0.1% (v/v) formic acid, 80% (v/v) acetonitrile). The mass spectrometer was operated in data-independent acquisition (DIA) mode with the following parameters: one full FTMS scan in Orbitrap (350 - 1 050 m/z) at 120 000 resolution, 20 ms injection time, and AGC target (%) = 300, followed by 175 fixed windows of 4 Da from 350 to 1 050 m/z, 5 ms injection time, and AGC target (%) = 500 (Astral detector). Precursor ions were fragmented with HCD, Normalized Collision Energy 26%, isolation window 4 m/z.

#### Data analysis

Acquired DIA spectra were searched using FragPipe computational platform v. 23.1^85^, MSFragger v. 4.3, Philosopher v. 5.1.2^91^, DIA-NN v. 1.8.2^92^. Peptide identification from MS/MS spectra was performed with MSFragger search engine against the Human Proteome reference dataset extended with contaminant list (http://www.uniprot.org/, downloaded in December 2025, 20 453 proteins) and reverse decoy sequences. Mass tolerance was set to 10 ppm for precursor and for fragmented ions. The search parameters were set to include fully tryptic peptides, allowing for up to one missed cleavage site, carbamidomethylation on cysteine residues as a static modification and up to three variable modifications including methionine oxidation and N-terminal acetylation. Peptide and protein inference was FDR controlled at 1% by PeptideProphet and ProteinProphet respectively. The generated library included entries for 257 251 precursors and 8 981 protein groups. For the DIA analysis, extraction of quantitative data was performed with DIA-NN v. 1.8.2 beta 8 with a maximum FDR of 1% and quantification strategy was set to “Robust LC”.

Protein group abundances were extracted from the file “report.tsv” and only protein groups with intensities > 0 and a PEP < 0.01 were tested for different normalization approaches^111^. Proteotypic precursors with abundance > 0 were log_2_-transformed and median-normalized. For each condition, the abundance of 8 694 proteins was calculated from the top three most intense precursors. In case of missing values (less than 1% of the entire dataset), missing intensities were replaced by random values sampled from a normal distribution centered below the minimum observed value (min - 1) with a standard deviation matching the observed data. Differential analysis for the abundance of 8 694 proteins was performed with limma package^93^.For WT comparison, differential protein abundance between doxycycline-treated and untreated states was assessed using moderated t-statistics implemented in the limma framework, with empirical Bayes variance shrinkage applied to stabilize gene-wise variance estimates. Differentially abundant proteins were defined based on log_2_FC thresholds (> 0.5 or < −0.5) and *p*value BH adjusted < 0.05.

For the comparison of the variants, a limma linear model without intercept was fitted to protein abundance values across 48 samples representing 12 experimental conditions (12D, 3W, ΔCR, 5R, 12A, G376V plus/minus doxycycline, N = 4 independent replicates per condition). Moderated t-tests were applied using empirical Bayes variance shrinkage, and contrasts were defined to compare each variant rescue versus KD condition, *p*values were BH adjusted.

Proteins with at least 2 concordant and significant changes per condition (PS-deficient and solid-like) (|log_2_FC| > 0.5 & *p*value BH adjusted < 0.05) (total 286 proteins) were considered for further analysis. PS-dependent proteins were defined as those in which the log_2_FC variability within each group (PS-deficient and solid-like) is lower compared to the global variability (total: 99 genes). Proteins were stratified based on the direction of regulation of the WT condition into upregulated (log_2_FC rescue versus KD > 0) and downregulated (log_2_FC rescue versus KD < 0). For visualization purpose, log_2_FC were standardized to z-score. GO enrichment was performed using clusterProfiler using the full proteome as a background (q < 0.01, *p*value < 0.05). Redundant terms were reduced using semantic similarity (Wang method) via GOSemSim/rrvgo (similarity > 0.9 or 0.85 respectively for up- and downregulated proteins), retaining representative terms based on significance. The entire dataset, including raw data, generated tables are available in the PRIDE repository (PRIDE: PXD076485)

### Network analysis and multi-omics integration

Proteomic datasets of PS-dependent upregulated and downregulated proteins were integrated with transcriptomic and alternative splicing data. For the network analysis, four datasets were utilized: i) PS-dependent interactors of TDP-43 (195 proteins), ii) 75 PS-dependent alternative splicing events, iii) 87 PS-dependent RNA expression changes, and iv) 99 PS-dependent protein abundance changes.

Functional relationships between PS-dependent protein abundance changes and PS-dependent interactors were established by integrating RNA-protein interaction data from the RNAInter database^112^. RNA-protein interaction data were sourced from RNAInter and filtered to include only human interactions supported by experimental evidence (either strong or weak, excluding predicted interactions) with interaction scores greater than 0.25. Interactions were further refined to include only those with PS-dependent proteins identified in the dataset as mRNA target (Interactor1.Symbol) and PS-dependent interactors as mRNA binder (Interactor2.Symbol).

Proteins regulated by TDP-43 phase separation in a splicing-dependent manner were defined as those present in both the PS-dependent protein abundance dataset and the PS-dependent alternative splicing dataset (n = 7). Proteins regulated through protein–protein interactions were defined based on the presence of curated functional links in the RNAInter database connecting them to phase separation–dependent interactors.

The resulting PS-dependent proteins, together with their connections to PS-dependent alternative splicing events and PS-dependent TDP-43 interactors, were visualized as an interaction network using Cytoscape (v. 5.0.2). In this network, experimentally validated targets of UPF1-mediated degradation, as reported by *Imamachi et al.*^108^., *Fritz et al.* ^109^, and *Colombo et al.* ^110^, were integrated into the RNAInter-derived interaction landscape to highlight the functional role of UPF1.

## Results

### *In silico* and *in vitro* analysis of the designed TDP-43 variants establishes a panel of “phase separation-deficient” and “solid-like” TDP-43 variants

To establish whether the phase separation state of TDP-43 influences its functionality, we designed a panel of TDP-43 variants which, based on literature knowledge, we predicted to be either condensation-deficient or condensation-prone. We reasoned that placing selected mutations that change, for example the charge, hydrophobicity or aromaticity in different positions of the C-terminal LCD would enable us to tune TDP-43’s condensation behavior in cells, eventually allowing us to study TDP-43 functionality across different condensation states while avoiding potential mutation-specific effects. Specifically, we designed a panel of six TDP-43 variants, aiming to obtain three TDP-43 variants with reduced condensation properties and three variants with enhanced condensation properties (Fig. 1A). In the 3W variant, three tryptophans (W334G, W385G, W412G) that are essential for LCD phase-separation^44^ were replaced with glycines, which tend to have spacer properties in IDRs^73^. In the ΔCR variant, we removed the α-helix-forming conserved region (CR, amino acids 321-340), which is a known driver of TDP-43 phase separation^42,43,60–62,113^. We also utilized a phosphomimetic 12D variant, for which we previously reported a reduced condensation/more liquid-like behavior; and the corresponding 12A variant, which we previously showed to yield more amorphous/aggregate-like condensates^71^. In the 5R variant, we combined five disease-associated mutations (G295R, Q343R, G348R, G357R, G384R), all of which introduce an arginine^114^, since arginine-specific interactions were shown to increase the viscosity of TDP-43 condensates^45^. Finally, we introduced the recently reported disease-linked point mutation G376V, which causes TDP-43 to form more amorphous, aggregate-like condensates^70^.

**Figure 1:**
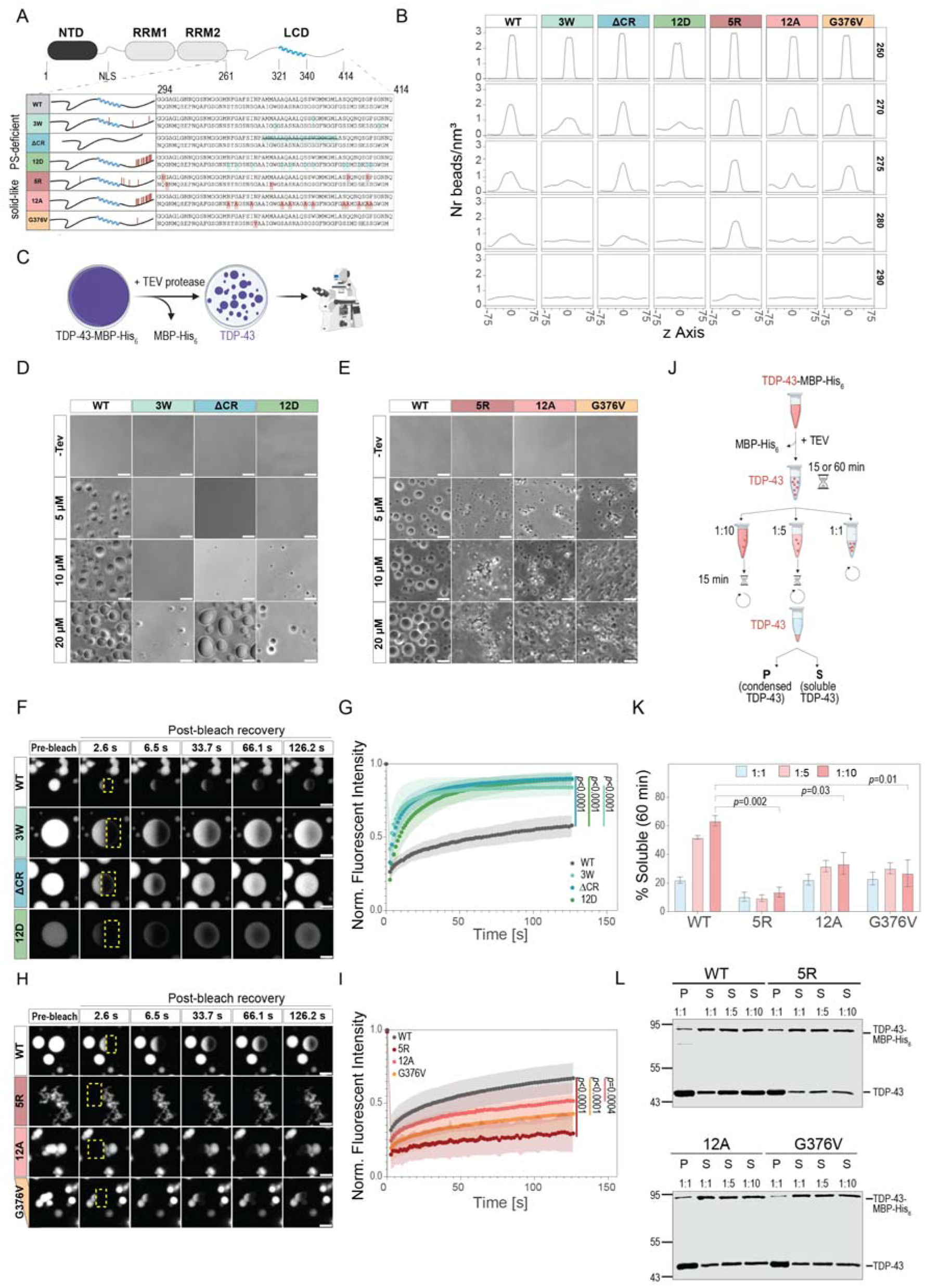
*In silico* and *in vitro* analysis of the designed TDP-43 variants establishes a panel of “phase separation-deficient” and “solid-like” TDP-43 variants. 1A. TDP-43 variants characterized in this study. Schematic diagram of the full-length TDP-43 with its N-terminal domain (NTD), RNA-recognition domains (RRM1, RRM2) and disordered LCD (aa. 261-414), containing the conserved region (CR, aa. 321-340) is shown. Variants designed to be phase separation-deficient are highlighted in turquoise, while variants designed to be more condensation-prone than WT are shown in red. Created in part with BioRender.com. 1B. Density profiles for condensates of the indicated TDP-43 variants calculated from CALVADOS3 coarse-grained simulations at different temperatures (in K, shown on the right). 1C. Scheme illustrating the workflow of the *in vitro* condensate microscopy assay. TDP-43-MBP-His_6_ is incubated with TEV protease to proteolytically remove the MBP solubility tag; reactions are incubated in phase separation buffer for 45 minutes and resulting condensates are imaged using brightfield microscopy. Created in part with BioRender.com. 1D, E. Representative brightfield microscopy images of condensates formed by the variants predicted to be PS-deficient (D) or condensation-prone (E) in comparison to TDP-43 WT at 5, 10 and 20 μM. Imaging was done 45 minutes post TEV addition. Negative controls without TEV protease addition (-TEV) contain TDP-43-MBP-His_6_ at 20 μM. Scale bar: 5 µm. 1F, H. Representative confocal fluorescence microscopy images for the FRAP experiments. Condensates from PS-deficient TDP-43 variants and WT as control were formed at 40 μM protein concentration (F), while for the amorphous mutants and corresponding WT control (H), a protein concentration of 15 μM was used. Bleached regions of interest (ROI) are indicated in yellow. Scale bar: 2 µm. 1G, I. Quantification of FRAP analysis illustrating a faster and higher fluorescence recovery for the PS-deficient variants compared to WT (G) and a lower fluorescence recovery of the amorphous variants compared to WT (I). An average of 14 condensates were analyzed per condition. Mean values are shown, with standard deviation indicated in the shaded regions. Significance was determined by quantifying areas under the curve for individual condensates followed by one-way ANOVA with a Dunnett’s multiple comparisons test to WT. *p*values are indicated. 1J. Workflow illustrating the condensate reversibility assay. Condensate formation was induced using TEV protease cleavage, followed by a 15- or 60-min incubation and a centrifugation step for undiluted (1:1) samples, or dilution (1:5 or 1:10) in phase separation buffer and an additional incubation for 15 minutes. Samples were centrifuged to separate supernatant (S, soluble TDP-43) and pellet (P, condensed TDP-43) fractions. Created in part with BioRender.com. 1K. Quantification of condensate reversibility. Dilution was performed at 60 minutes post condensate induction. % soluble values were determined by quantifying the signal coming from the soluble (S) fraction as a percentage of total signal (P+S fractions) for a given mutant. Error bars indicate standard deviation across three independent experiments. Significance was determined for the signal obtained in 1:10 dilutions using a one-way ANOVA with a Dunnett’s multiple comparisons test to WT. *p*values are indicated. 1L. Representative western blot images for TDP-43 WT, or the less dynamic 5R, 12A and G376V variants tested for condensate reversibility after 60 minutes of incubation time. Pellet (P) and supernatant (S) fractions were loaded for undiluted (1:1) samples, while for 1:5 and 1:10 samples only supernatant (S) fractions are shown. Blots were incubated with a TDP-43-specific antibody (80001-1-RR, Proteintech).

We employed molecular dynamics (MD) simulations with an implicit-solvent, residue-level coarse-grained model^64^ to predict the phase separation behavior of these six variants in comparison to TDP-43 wild-type (WT). MD simulations allowed us to trace molecule trajectory and to probe condensate stability as a function of the thermal energy provided to the system (. 1B, Extended Data Fig. 1A). As expected, the molecular mobility progressively increased with increasing temperature, leading to enhanced diffusion; hence at 290K all TDP-43 variants were fully dispersed, and no condensed phase was observed (Fig. 1B). Analysis of the density profiles revealed that condensates formed from the 5R variant were most resistant to increasing temperature. G376V, 12A and the ΔCR variant were similar to WT and showed a tendency to form condensates across temperatures, with condensate dispersal starting at 280K. In contrast, the 3W and 12D variants showed a much lower tendency for condensate formation than WT, with a higher tendency to distribute in the dilute phase (Fig. 1B). Encouraged by the *in silico* analysis, which for the majority of our designed variants validated the anticipated behavior, we advanced to experimentally evaluate the PS properties of the designed variants.

To assess the PS behavior of the designed variants *in vitro*, we purified bacterially expressed TDP-43 variants with an MBP solubility tag^115^, induced their condensation by TEV protease-mediated removal of the MBP tag in a physiological buffer and monitored condensate formation by brightfield microscopy (Fig. 1C). In comparison to wild-type (WT), TDP-43, 3W, ΔCR and 12D showed a clear reduction in phase separation propensity, with visible condensate formation starting only at 10 μM for ΔCR and 12D, and 20 μM for 3W (Fig. 1D). At high protein concentration (20 µM), the 3W and 12D variants formed only few small, round condensates, while ΔCR assembled into much larger condensates that readily fused (Fig. 1D). On the other hand, 5R, 12A and G376V formed small, amorphous or aggregate-like structures, which did not fuse and tended to form agglomerates, instead of round condensates seen for TDP-43 WT (Fig. 1E). Using TDP-43 variants sparsely labeled with a small fluorescent dye, Alexa Fluor 488 (AF488), we recapitulated these condensate phenotypes, while also being able to visualize very small condensates formed by the 3W, ΔCR and 12D variants at 5 µM protein concentration (Extended Data Fig. 1B), which were undetectable in assays with unlabeled protein.

As the above-described TDP-43 variants led to striking changes in condensate morphology, we proceeded to assess condensate dynamics using fluorescence recovery after photobleaching (FRAP) analysis. To this end, we formed condensates of TDP-43 WT and the 3W, ΔCR and 12D variants sparsely labeled with AF488 at 40 µM protein concentration, performed half bleaches and measured the fluorescence recovery over the timecourse of ∼2 minutes. All three variants showed a faster and higher degree of fluorescence recovery in the bleached region compared to TDP-43 WT (Fig. 1F, G). 3W, ΔCR and 12D condensates readily fused, consistent with their larger size compared to WT condensates. We did the same comparison for TDP-43 WT and the more amorphous 5R, 12A and G376V variants at 15 µM protein concentration. All three variants showed slower fluorescence recovery, with 5R showing the strongest and 12A the most moderate reduction (Fig. 1H, I). To address whether any of the three less dynamic TDP-43 variants show a lower saturation concentration (C_sat_) for condensate formation than TDP-43 WT, we quantified the TDP-43 concentration in the dilute phase by sedimentation analysis followed by SYPRO Ruby staining. This revealed a significantly decreased C_sat_ for 5R and a weaker reduction of C_sat_ for G376V compared to WT, while no difference in C_sat_ was seen for the 12A variant (Extended Data Fig. 1C, D). To further corroborate the lower dynamicity phenotype of the 5R, 12A and G376V variants, we tested the reversibility of condensates formed by these mutants. To this end, 15- or 60-minute-old condensates were either centrifuged to separate soluble (supernatant) or condensed (pellet) fractions, or diluted 1:5 or 1:10, followed by an additional 15- or 60-minute incubation and the same centrifugation step (see scheme in Fig. 1J). For TDP-43 WT, increasing levels of protein were detected in the soluble fraction upon dilution, indicating condensate reversibility. In contrast, the 5R, 12A and G376V variants all showed similar amounts of TDP-43 in the soluble fraction even after the dilution steps, indicating lower condensate reversibility and/or accelerated condensate ageing (Fig. 1K, L, Extended Data Fig. 1E, F).

Collectively, our *in vitro* characterization of the designed TDP-43 variants (Fig. 1A) demonstrates that by introducing rationally designed mutations in the LCD, we were able to successfully tune TDP-43 phase separation in both directions, creating a panel of phase separation-impaired, more dynamic and liquid-like TDP-43 condensates (3W, ΔCR, 12D), and a panel of amorphous, less dynamic and solid-like TDP-43 condensates (5R, 12A, G376V). For simplicity, we will refer to the first panel (3W, ΔCR, 12D) as “PS-deficient”, and to the second panel (5R, 12A, G376V) as “solid-like” TDP-43 variants.

### Rationally designed LCD mutations tune TDP-43 phase separation behavior in cells

As a next step, we wanted to test whether our rationally designed TDP-43 variants tune TDP-43’s phase separation behavior also in cells. As phase separation is a highly concentration-dependent process, we utilized a cellular system where TDP-43 variant levels can be well-controlled. We integrated the different TDP-43 variants into HeLa Flp-In T-REx cells, in which a single copy of a gene-of-interest can be stably integrated and then expressed in a doxycycline-inducible manner. To study the different TDP-43 variants in a physiological environment, we depleted endogenous TDP-43 using siRNA and then expressed TDP-43 WT or the different TDP-43 variants with an N-terminal EGFP tag for 24 hours (Fig. 2A).

**Figure 2.**
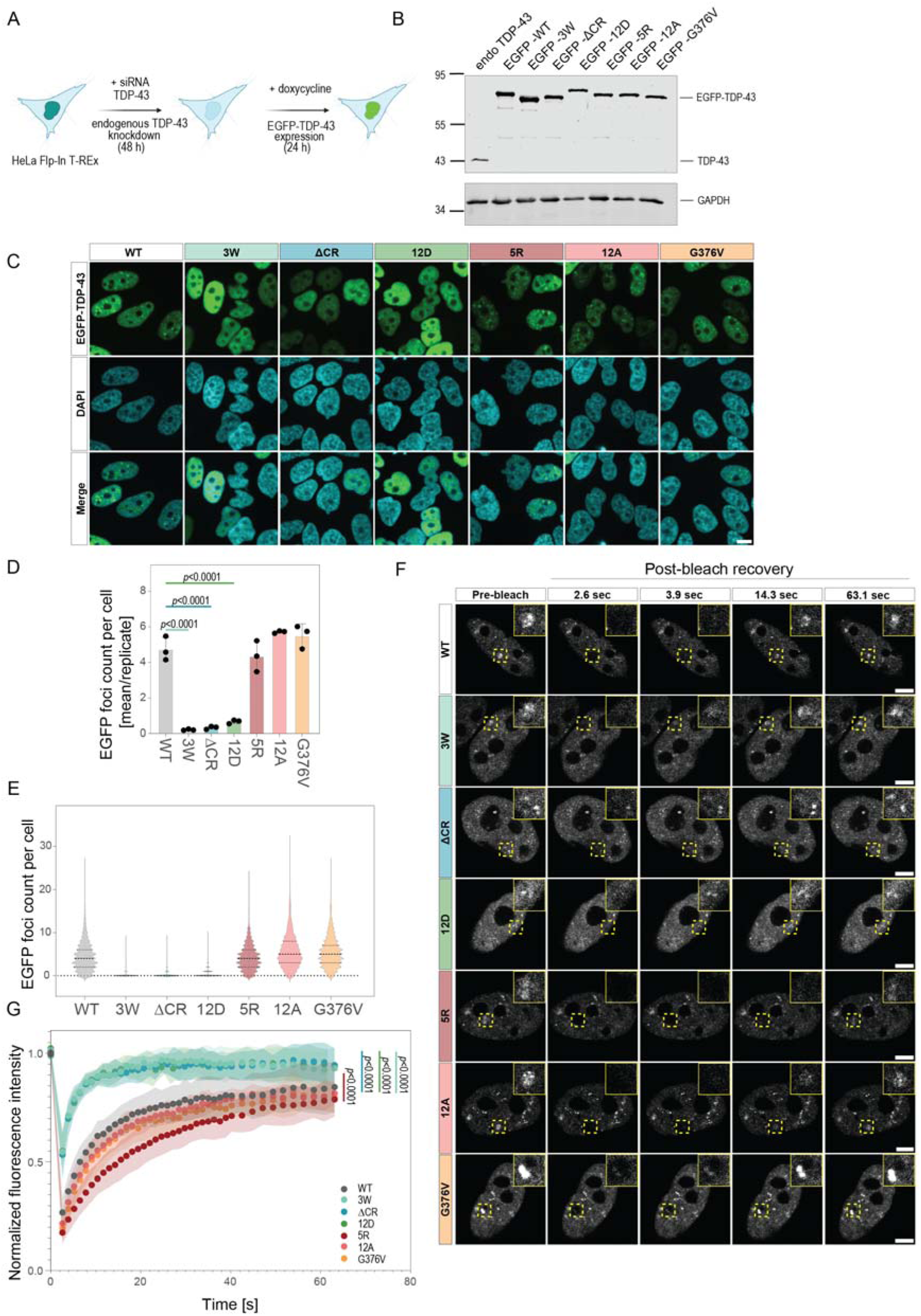
Rationally designed LCD mutations tune TDP-43 phase separation behavior in cells. 2A. Scheme illustrating the experimental setup in all cellular experiments with EGFP-TDP-43 variants. Silencing of endogenous TDP-43 by siRNA was done for 48 hours, followed by doxycycline-induced expression of EGFP-tagged TDP-43 variants for 24 hours. Created in part with BioRender.com. 2B. Western blot showing EGFP-TDP-43 expression levels in HeLa Flp-In T-REx cells expressing the indicated EGFP-TDP-43 variants in the absence of endogenous TDP-43 and after doxycycline induction. Untreated HeLa Flp-In T-Rex EGFP-TDP-43 WT cells without TDP-43 KD or doxycycline induction were used to illustrate endogenous TDP-43 levels (“endo TDP-43”). GAPDH blot is shown as a loading control. Antibodies used: mouse anti-TDP-43 (60019-2-Ig, Proteintech), rabbit anti-GAPDH (10494-1-AP, Proteintech). 2C. Representative confocal images of HeLa Flp-In T-REx cells expressing the indicated EGFP-TDP-43 (green). Nuclei were stained with DAPI (blue). Maximum intensity projections are shown. Scale bar: 10 μm. 2D. Quantification of nuclear TDP-43 foci from confocal high-throughput images shown in (C). Single points show the mean counts of EGFP-TDP-43 foci per cell in each experiment (∼15 fields of view were quantified per replicate). Mean values of three independent experiments are shown in a barplot. Error bars indicate standard deviation. Significance was determined using a one-way ANOVA with a Dunnett’s multiple comparisons test to WT. *p*values are indicated. 2E. Distributions of foci counts per cell. The median is indicated by a thick dashed line, and the 1^st^ and 4^th^ quartiles are shown in dotted lines. ∼2 000 cells were quantified across three replicates. 2F. Representative FRAP images in HeLa Flp-In T-REx cells expressing the EGFP-tagged TDP-43 variants. Bleached nuclear regions are indicated in yellow dashed squares. Top right quadrants show a zoomed-in image of the bleached region. Scale bar: 5 μm. 2G. Quantification of fluorescence recovery from FRAP experiments shown in (F). Regions of interest in nuclei were bleached and their fluorescence recovery was measured for ∼1 minute. ∼13 cells were analyzed per cell line across three biological replicates, mean values of all fluorescence recovery curves are plotted, with standard deviation indicated in the shaded region. Significance was determined by quantifying areas under the curve for individual condensates followed by one-way ANOVA with a Dunnett’s multiple comparisons test to WT. *p*values are indicated.

Western blots demonstrated that the TDP-43 knockdown was highly efficient, and by titrating doxycycline amounts, we were able to achieve similar levels with an approximately two-fold overexpression of all EGFP-TDP-43 variants (Fig. 2B).

TDP-43 is known to enrich in nuclear foci, such as Cajal bodies, gems and paraspeckles, but also foci of unknown identity and function^40,116^. When we examined TDP-43 localization by high-throughput confocal microscopy in the above-described cellular model system under knockdown of endogenous TDP-43, we observed that the different TDP-43 PS variants showed varying tendencies to localize to such foci. All three PS-deficient variants (3W, ΔCR, 12D) remained more diffuse than TDP-43 WT, showing fewer nuclear TDP-43 foci per cell (Fig. 2C, D, E). On the other hand, solid-like TDP-43 variants (5R, 12A, G376V) showed similar nuclear foci numbers as TDP-43 WT, with some cells having more TDP-43 puncta compared to WT (Fig. 2C, D, E). Notably, all TDP-43 variants displayed physiological nuclear localization, which enables us to compare TDP-43 functionality in the nucleus.

TDP-43 is also responsive to various types of cellular stress and is recruited into large stress-induced nuclear foci^117,118^. To test how TDP-43’s recruitment into stress-induced nuclear foci is affected by the PS-altering mutations, we exposed cells to heat stress (1 hour at 43°C) followed by 3-hour recovery at 37°C (Extended Data Fig. 2A). Also in this stress paradigm, all three PS-deficient variants (3W, ΔCR, 12D) showed fewer nuclear foci than TDP-43 WT, whereas the solid-like TDP-43 variants (5R, 12A, G376V) showed similar numbers of stress-induced TDP-43 foci (Extended Data Fig. 2B, C, D).

Additionally, we sought to determine whether the dynamics of TDP-43 in nuclear foci are altered in cells expressing PS-deficient or solid-like TDP-43 variants. To this end, we performed FRAP experiments by bleaching a region in the nucleus containing TDP-43-enriched foci and measuring fluorescence recovery. All PS-deficient TDP-43 variants showed significantly faster and more efficient fluorescence recovery than TDP-43 WT (Fig. 2F, G), indicating a higher dynamicity of the 3W, ΔCR, and 12D variants in cells. In contrast, all three solid-like variants showed slower fluorescence recovery than TDP-43 WT, which was significant for the 5R variant (Fig. 2F, G). We also conducted FRAP experiments in the above-described heat stress recovery paradigm (Extended Data Fig. 2A), by performing point bleaches on the larger, stress-induced nuclear TDP-43 foci. Also in these stress-induced nuclear TDP-43 assemblies, all three PS-deficient variants remained significantly more dynamic than TDP-43 WT, whereas a trend towards reduced fluorescent recovery was seen for all three solid-like variants (Extended Data Fig. 2E, F).

To be able to study TDP-43 behavior and functionality in the absence of the large EGFP-tag, we generated HeLa Flp-In T-Rex cells with inducible expression of N-terminally myc-tagged TDP-43 variants at near endogenous levels (Extended Data Fig. 3A, B). We reasoned that using a small epitope tag and close to physiological expression levels should best preserve TDP-43’s native phase separation behavior and functionality. All myc-TDP-43 PS variants properly localized to the nucleus and showed varying tendencies to form nuclear foci, similar to those observed in EGFP-tagged variants: PS-deficient variants exhibited fewer foci compared to TDP-43 WT, and solid-like variants formed similar or higher numbers of nuclear foci (Extended Data Fig. 3C, D). This cellular model was used for subsequent functional analysis.

In summary, altered TDP-43 condensation behavior that we observed *in vitro* for our rationally designed TDP-43 variants (Fig. 1A) translates into phenotypic changes in cells, affecting the distribution and dynamicity of TDP-43 in physiological and stress-induced nuclear foci.

### The phase separation behavior of TDP-43 governs its protein interactome

To evaluate whether the differential phase separation behavior of our rationally designed TDP-43 variants affects TDP-43 protein interactions in the cellular environment, we performed affinity purification coupled to mass spectrometry (AP-MS) (Fig. 3A). In the experimental setup, TDP-43 WT, the three PS-deficient TDP-43 variants (3W, ΔCR and 12D) and the three solid-like variants (5R, 12A and G376V) were expressed with a myc-tag in a doxycycline-inducible manner using the above-described HeLa Flp-In T-REx system in the absence of endogenous TDP-43 (see Supplementary Fig. 1A for expression levels of the different TDP-43 variants). After mild cell lysis, TDP-43 interactors were purified using an anti-myc antibody, and the co-purified proteins were analyzed by mass spectrometry (Fig. 3A, Supplementary Table 2). To filter out protein contaminants, we performed a negative-control purification using non-induced TDP-43 WT cells and the same anti-myc antibody. The number of proteins identified and their abundance were consistent across all conditions (Supplementary Fig. 1B, C), and TDP-43 was the most significantly enriched protein, with similar amounts being co-purified in all samples, except for the non-induced control cells (Supplementary Fig. 1D, E, F). A threshold of log_2_FC > 1 and *p*value adjusted < 0.05 compared to the control purification in at least one TDP-43 variant condition led to the identification of 226 TDP-43 interactors. Gene ontology (GO) analysis identified that TDP-43 interactors were characterized by a strong enrichment in proteins that bind to RNA and regulate RNA metabolism (e.g. mRNA splicing via spliceosome, mRNA transport and gene expression (Fig. 3B). Cellular component analysis showed an enrichment for nucleus-associated compartments, in agreement with TDP-43’s primary localization in the nucleus, as well as membrane bound organelles (e.g. mitochondria) (Fig. 3B). Most of the enriched GO terms were associated with the solid-like TDP-43 variants, and in particular 5R exhibited the highest number of interactors involved in RNA-regulatory pathways (Supplementary Fig. 1G, H, I). We benchmarked the interactors identified in the generated dataset against a repository of protein-protein interactions (BioGRID v. 5.0.253)^83^ integrated with TDP-43 interactomics datasets (AP-MS, BioID and APEX) ^20,25,31–33,80–82^ not available in BioGRID (see materials and methods). We defined high-confidence interactors (HCIs) as proteins detected in more than three independent interactomics datasets, to distinguish them from proteins identified less consistently. Our acquired dataset largely recapitulates previous TDP-43 interactome studies, with over half of the proteins having been previously reported as TDP-43 interactors. Among these, 39 proteins were identified as HCIs (Fig. 3C).

**Figure 3:**
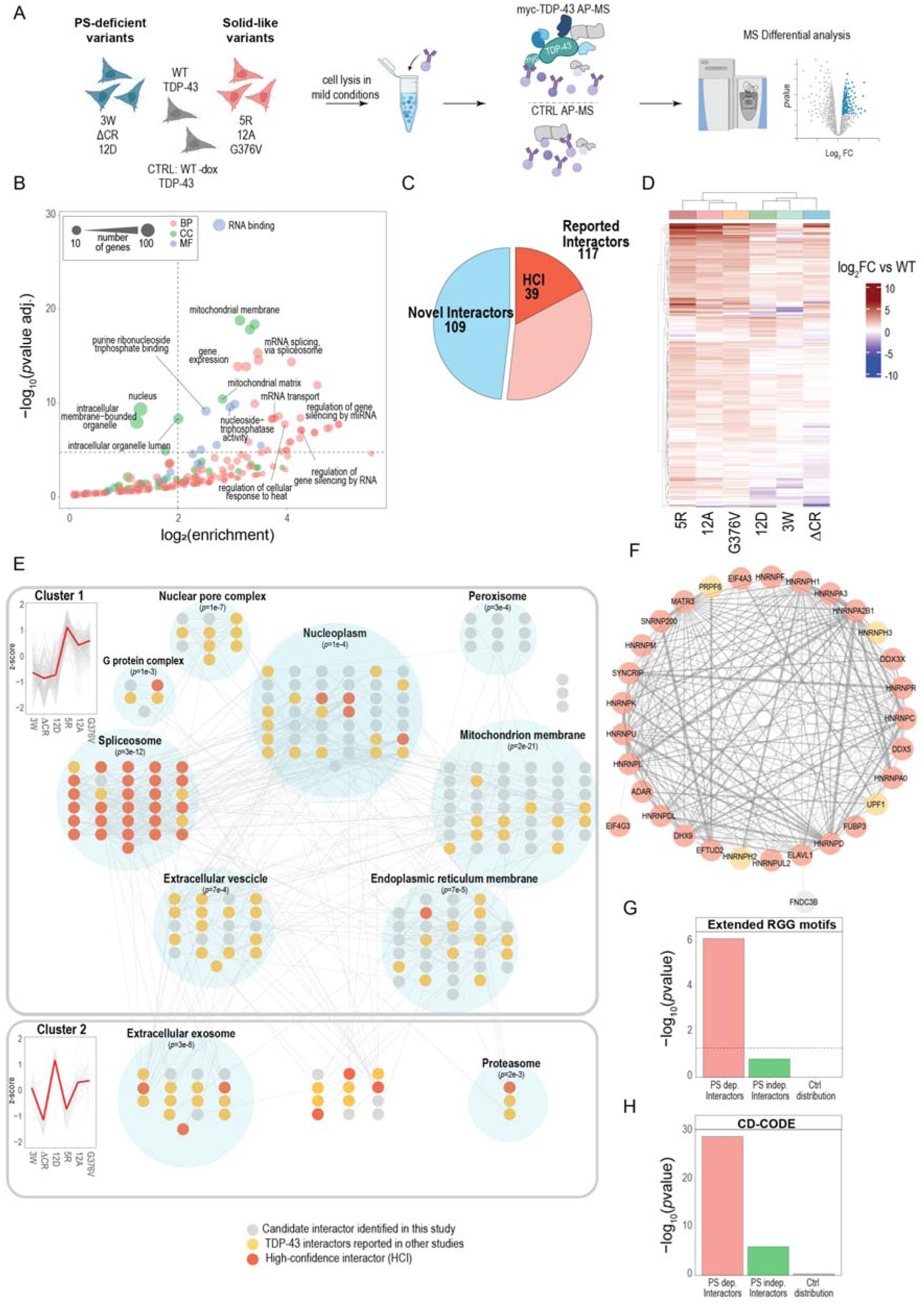
The phase separation behavior of TDP-43 governs its protein interactome. 3A. Workflow for the identification of TDP-43 interactors by AP-MS. Phase separation-deficient variants (3W, ΔCR, 12D), solid-like variants (5R, 12A, G376V) or TDP-43 WT were expressed with a myc tag to enable co-purification of protein interactors under mild lysis conditions (N = 3). Co-purified proteins were subjected to MS identification and quantification. To filter out non-specific interactors, the purification was also performed for a mock condition without the expression of TDP-43. All purifications were done in the absence of endogenous TDP-43 (after siRNA-mediated depletion). Created in part with BioRender.com. 3B. GO enrichment analysis of TDP-43 interactors. GO terms with at least 10 genes associated, a log_2_FC enrichment > 1, and with a *p*value < 1 × 10^−5^ are labeled. To reduce redundancy, terms were grouped based on similarity and only parental terms were shown. Circle corresponds to the number of genes per term, and color indicates the GO category (red: Biological Process (BP); green: Cellular Component (CC); blue: Molecular Function (MF)). 3C. Overlap of TDP-43 interactors identified in this dataset with interactors reported from other TDP-43 interactomics experiments. Novel interactors are shown in blue, reported interactors are shown in shades of red. HCIs: high-confidence interactors, defined as TDP-43 interactors identified in at least three independent interactomics datasets (shown in dark-red). 3D. Unsupervised hierarchical clustering of TDP-43 interactor levels identified for solid-like and PS-deficient variants of TDP-43. Values reported in the heatmap show the average (three independent replicates) intensity of interactors as log_2_FC compared to TDP-43 WT. Proteins with an enhanced interaction intensity are shown in red, while those with weaker interactions are shown in blue. 3E. Protein-protein interaction network of TDP-43. Proteins identified as TDP-43 interactors are clustered based on the abundance profile across all TDP-43 purifications (plot profile on the left). Proteins are organized in modules according to reported protein-protein interactions and their localization. The edges (connection between two nodes) indicate interactions identified in at least two independent experiments (BioGRID v. 5.0.253)^83^. Novel, previously reported, as well as high-confidence interactors are shown as nodes in different colors (gray, yellow and red respectively). Functional enrichment analysis was performed using STRING, and statistical significance was assessed using a hypergeometric test. 3F. Protein-protein interaction network for the spliceosome module. The width of the edge between the nodes is proportional to the number of independent studies describing the interactions. Novel interactors, reported interactors, and high-confidence interactors are indicated in gray, yellow, and red, respectively. 3G, H. Enrichment analysis for TDP-43 interactors (PS-dependent/PS-independent) and for the control group. Enrichment was evaluated for proteins containing an extended RGG motif G(0-3)X(0,1)RG(1,2)X(0,5)G(0,2)X(0,1)RG(1,2)^86^ in (G), and for proteins annotated as associated to phase separation from the CD-CODE database (only proteins reported to participate in biomolecular condensates in at least three independent experiments were considered) in (H). Reported values correspond to enrichment calculated using a hypergeometric test.

Additionally, we aimed to validate these TDP-43 interactors by verifying their functional association with TDP-43, as well as their potential for direct binding to TDP-43. In the first case, we generated a network of proteins using the STRING database (v.12)^85^ including direct and indirect functional relationships to calculate minimal-cost paths distances between identified interactors and TDP-43. Proteins identified as interactors in the AP-MS dataset ranked among the top 20% of TDP-43 functionally associated proteins (Extended Data Fig. 4A). Secondly, we tested with MD simulations whether the protein pairs composed of one chain of TDP-43 and one chain of an identified interactor are more likely to bind to each other compared to a control group composed of TDP-43 and proteins that did not pass the interactor filtering criteria^68,119^. Pairs consisting of TDP-43 and its interactors showed a significantly higher fraction of frames in a predicted bound state than the control group, corresponding to lower *in silico* Kd values (*p*value = 0.004; Extended Data Fig. 4B, C). In parallel, TDP-43 and its interactors exhibited significantly more contact events than the control group (*p*value = 5.4 × 10⁻), indicating a stronger propensity for direct interaction relative to the control (Extended Data Fig. 4D).

To evaluate whether the interacting proteins were differentially purified based on the phase separation properties of TDP-43 and to ensure that variations in interactor abundance were not due to differences in the bait purification yield, the abundance of the identified interactors was normalized for the bait amount, and the resulting levels were compared to the WT condition (Fig. 3D). Unsupervised hierarchical clustering, as well as principal component analysis and cluster correlation of the analyzed purifications indicated two clear and distinct groups which coincided with the phase separation properties of TDP-43 (solid-like and PS-deficient) (Fig. 3D, Supplementary Fig. 2A, B). Notably, the solid-like TDP-43 variants showed a massive co-purification of interactors compared to the PS-deficient variants (Fig. 3D). Clustering profile of proteins across all analyzed conditions led to the identification of two groups: one containing interactors whose abundance is dependent on TDP-43 phase separation behavior with stronger affinity to solid-like TDP-43 variants (cluster 1, “PS-dependent TDP-43 interactors”, 195 proteins) and another group of interactors, whose abundance is independent of TDP-43 phase separation (cluster 2) (Supplementary Fig. 2C). Subsequently, we generated a network in which we overlapped all identified TDP-43 interactors with annotated interactions (BioGRID v. 5.0.253). We then compared this network to 100 randomized control networks sampled from the BioGRID database. TDP-43 interactome network shows more interactions, more connections per protein, and greater transitivity (the tendency of proteins to be highly interconnected) than the controls (Supplementary Fig. 2D, E, F). These findings indicate that the identified TDP-43 interactors are highly interconnected and can be organized into distinct groups (modules) based on physical and functional associations ^120,121^ (Fig. 3E). Among the modules that bind with high affinity to solid-like variants we identified several components of the nuclear pore complex, the spliceosome (Fig. 3F), the nucleoplasm, and various membrane-bound organelles (ER, mitochondrial membrane, extracellular vesicle and peroxisome) (see Supplementary Fig. 2G for the abundance profile of the modules and Supplementary Fig. 3A for the protein organization within the modules). Network centrality analysis (eigenvector) revealed that proteins forming the spliceosome module occupy central positions within the generated network (Supplementary Fig. 2H). The spliceosome complex is functionally linked to TDP-43 due to the protein’s known roles in splicing regulation^122–124^, and indeed we identified the majority of the proteins belonging to this group to be high-confidence interactors of TDP-43 (Fig. 3F). Taken together, our findings indicate that numerous splicing regulators bind to TDP-43 and that the affinity of these interactions is governed by the phase separation properties of TDP-43.

To identify specific features among the PS-dependent TDP-43 interactors, we compared PS-dependent and independent interactors to a randomly sampled group of proteins from the human proteome. We found that PS-dependent TDP-43 interactors exhibit a higher protein disorder (*p*value = 1.9 × 10^−4^, Supplementary Fig. 3B), a reduced helix content, and an increased proportion of coiled regions (Supplementary Fig. 3C).

Short linear motifs are short protein regions characterized by low amino acid complexity and high flexibility, which can promote multivalent interactions and drive phase separation^125,126^. Among these, the arginine-glycine-rich RGG motif, is well known to promote protein-RNA binding and protein phase transitions^127^. As several TDP-43 interactors were involved in RNA binding (Fig. 3B), we calculated the enrichment of RG, RGG and extended RGG motifs^86^ among the PS-dependent TDP-43 interactors and identified a significant enrichment of these motifs (*p*value = 6.9 × 10^−7^, Fig. 3G, Supplementary Fig. 3D, E).

The lower pLLDT score, the shift in secondary structure composition and the enrichment in RG/RGG motifs support the hypothesis that the interactions between this group of proteins and TDP-43 might be governed by phase separation. Hence, we sought to analyze to which extent the interactors themselves are prone to undergo condensation. We found that PS-dependent interactors are significantly enriched in proteins annotated to be phase-separating in the CD-CODE^88^ and PhaSepDB^87^ databases (Fig. 3H and Supplementary Fig. 3F).

Collectively, we identified common features among the PS-dependent TDP-43 interactors, e.g. a high degree of disorder, enrichment in RG/RGG motifs and high PS propensity. This suggests that their association with TDP-43 might come from a multivalent network of interactions driven by phase separation, where solid-like TDP-43 variants form a stronger network of interactions, thus selectively binding certain interactors more efficiently than TDP-43 WT or PS-deficient TDP-43 variants.

### Phase separation properties govern co-condensation of proteins with distinct sequence features in the cellular lysate

To investigate whether proteins that preferentially interact with the solid-like TDP-43 variants in AP-MS experiments selectively co-condense with solid-like TDP-43 variants, we modified and further implemented the experimental approach used by *Freibaum et al.* for reconstituting nuclear and cytoplasmic condensates in total HeLa cell lysates^80^. We spiked recombinant TDP-43-MBP-His_6_ variants into a standardized HeLa cell lysate (2 mg/ml) and induced TDP-43 condensate formation by the addition of TEV protease in the crowded, multicomponent environment resembling the cellular milieu. Using fluorescently labelled TDP-43 variants, condensates were visualized through confocal microscopy. Alternatively, using reactions with unlabeled TDP-43 variants combined with centrifugation, we quantified the abundance of TDP-43 and co-depleted proteins in the supernatant by western blotting and mass spectrometry (Fig. 4A).

**Figure 4:**
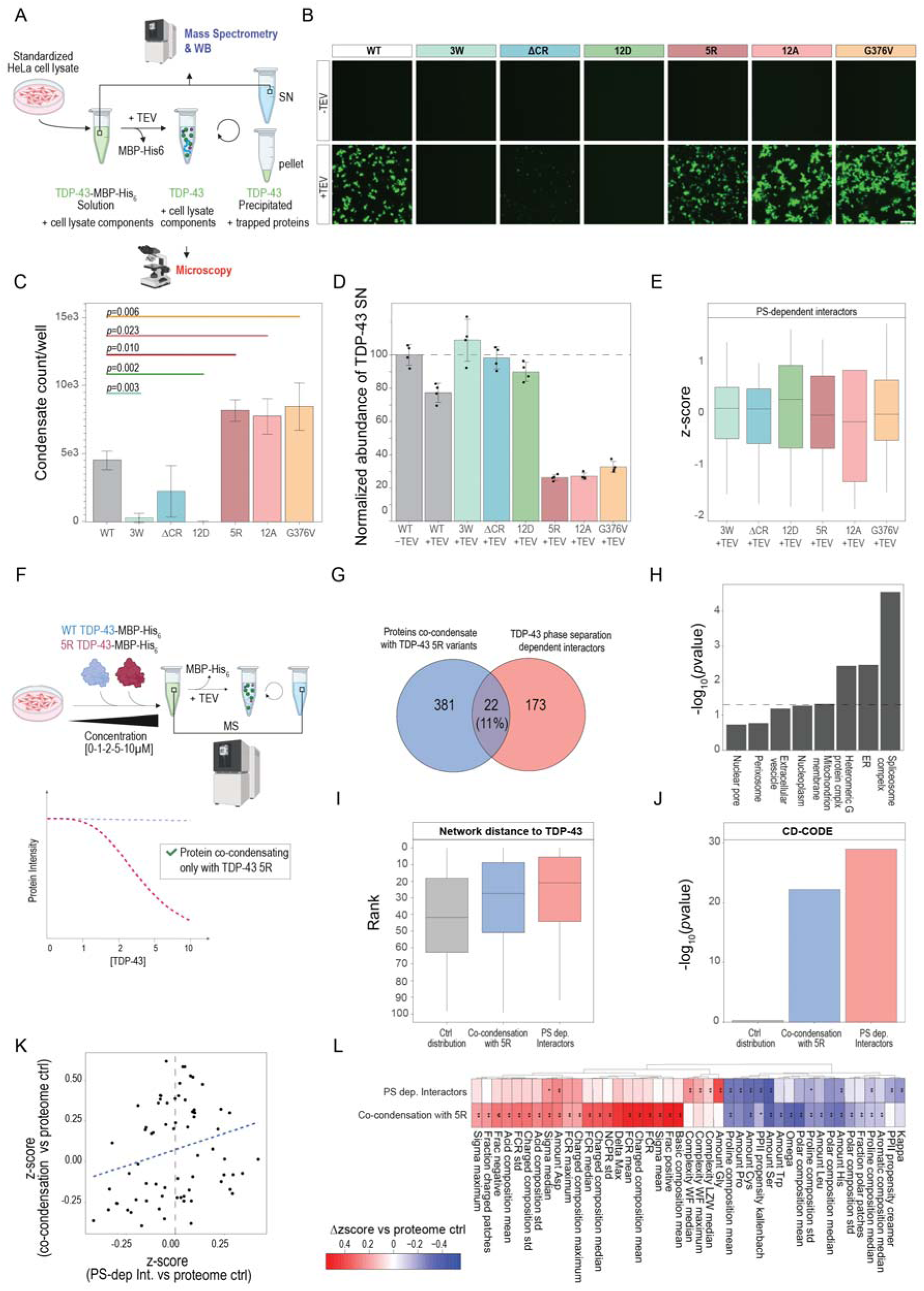
Phase separation properties govern co-condensation of proteins with distinct sequence features in the cellular lysate. 4A. Experimental scheme for the co-condensation experiment. Purified TDP-MBP-43-His_6_ was spiked into standardized HeLa cell lysate (2 mg/ml) and formation of condensates was triggered by TEV protease addition. Formed condensates were examined by confocal fluorescence microscopy or pelleted by centrifugation. The supernatant and pellet fractions were examined by western blotting (WB), and soluble proteins in the supernatant were quantitatively analyzed by mass spectrometry. Created in part with BioRender.com. 4B. Representative confocal microscopy images of TDP-43 condensates formed at 10 μM protein concentration in the presence of HeLa cell lysate (2 mg/ml) upon TEV cleavage (45 minutes). TDP-43-MBP-His_6_ proteins were labeled with Alexa Fluor 488. Maximum intensity projections are shown. Scale bar: 10 µm. 4C. Quantification of the number of condensates per well for the images shown in (B). 12 fields of view were quantified per condition. Mean values of 3 experimental replicates are shown. Error bars indicate standard deviation. Significance was determined using one-way ANOVA with a Dunnett’s multiple comparisons test to WT. *p*values are indicated. 4D. Abundance of soluble TDP-43 measured in the supernatant fraction (SN) by mass spectrometry. Abundance values were normalized to the corresponding TDP-43 levels in the absence of TEV (first column). The bar plot represents the mean values, error bars indicate the standard deviation, and individual points correspond to biological replicates (N = 4). 4E. Abundance levels of TDP-43 PS-dependent interactors (N = 43). Only interactors whose abundance changed significantly under at least one condition were considered (*p*value adjusted < 0.05). The level of endogenous soluble proteins in the supernatant after the centrifugation is reported as z-score, calculated from the mean protein value for each condition. Boxplot reports the distribution of the values with the median (central line), the interquartile range (1^st^ and 3^rd^ quartile) and whiskers (1.5XIQR, interquartile range). 4F. Experimental scheme for the dose-response co-condensation profiling upon the titration of TDP-43 WT and 5R. Increasing amounts of purified protein (0, 1, 2, 5, 10 µM) were added to the total HeLa cell lysate (2 mg/ml) and protein levels were measured by mass spectrometry before and after TEV protease addition and centrifugation. 403 proteins displaying a dose-dependent depletion from the supernatant following increasing concentrations of spiked-in TDP-43 5R variant, but not WT, were classified as proteins co-precipitating with solid-like mutants (example profile shown below). See materials and methods and Fig. S9C for details on the data analysis procedure. Created in part with BioRender.com. 4G. Venn diagram showing the overlap between the proteins identified as PS-dependent interactors by AP-MS (Fig. 3) and proteins co-precipitating specifically with the 5R variant. Dose-response co-condensation profiling confirmed co-precipitation of 22 proteins with the 5R variant, representing ∼11% of the 195 proteins previously identified in the AP-MS dataset. 4H. Enrichment analysis for TDP-43 modules identified in the AP-MS dataset and proteins co-condensing with 5R. Reported values correspond to enrichment calculated using a hypergeometric test. 4I. Functional association of proteins with TDP-43 was analyzed across three groups: control (403 randomly sampled proteins from the identified dataset), co-condensing with 5R (403 proteins), and PS-dependent interactors (195 proteins). For each group, the shortest path to TDP-43 within the STRING network was calculated, and a path score was determined by multiplying the interaction confidence values along the path. Proteins were then ranked based on their respective path scores. Boxplot reports the distribution of the values with the median (central line), the interquartile range (1^st^ and 3^rd^ quartile) and whiskers (1.5XIQR, interquartile range). 4J. Enrichment analysis for proteins annotated to undergo phase separation from the CD-CODE database for PS-dependent interactors, proteins co-condensing with 5R and random control proteins. Only proteins reported to partition into biomolecular condensates in at least three independent studies were considered. Reported values correspond to enrichment calculated using a hypergeometric test. 4K. Correlation between significant grammar sequence features (N = 74, *p*value adjusted < 0.01, Welch test) of TDP-43 PS-dependent interactors and proteins co-condensing with 5R. Sequence parameter values for each protein were calculated using CIDER^95^ and expressed as z-scores relative to the mean and standard deviation of the human proteome. Each point represents a grammar sequence feature deviation from the proteome average, with positive and negative values indicating enrichment or depletion, respectively. 4L. Conserved grammar sequence features for proteins identified as PS-dependent interactors and co-condensing with 5R. A scale from blue (negative values) to red (positive values) displays respectively the depletion and enrichment of properties compared to the mean properties of the human proteome. The significance of the deviation from the human proteome is reported (Welch test, adjusted *p*values are as follows: ** < 0.01 and * < 0.1). Among the 128 grammar features calculated, this heatmap only reports features that show significant differences compared to the human proteome in at least one condition and a coherent enrichment between the two conditions.

Microscopy analysis revealed that TDP-43 WT at 10 µM concentration efficiently formed condensates in the HeLa cell lysate (Fig. 4B). In line with our homotypic *in vitro* phase separation experiments (Fig. 1D, E), 3W, ΔCR, and 12D showed a strong deficiency in condensate formation, whereas 5R, 12A, and G376V formed more condensates compared to WT (Fig. 4B, C), with 12A and G376V showing a larger area covered by the condensed fraction (Supplementary Fig. 4A). In parallel, we quantified the abundance of TDP-43 in the supernatant and pellet fractions by western blotting and used mass spectrometry to measure the depletion of proteins from the supernatant. Both approaches showed that the major part (∼70%) of solid-like TDP-43 variants was predominantly lost from the supernatant and partitioned into the pellet fraction after lysate centrifugation, while PS-deficient TDP-43 variants remained largely soluble and barely pelleted (Supplementary Fig. 4B, Fig. 4D).

To identify proteins that co-condense and thereby co-pellet with the different TDP-43 variants, we monitored the depletion of endogenous proteins from the supernatant by mass spectrometry (see Supplementary Fig. 4C, D, E for quality control for the proteomics analysis). In total, 371 proteins were depleted (log_2_FC < −0.3, *p*value adjusted < 0.1) in at least one TDP-43 variant (Supplementary Fig. 4G, Supplementary Table 3). Most of the depletion events were observed upon the addition of solid-like TDP-43 variants (in particular 12A and 5R) (Supplementary Fig. 4F). Among the identified depleted proteins, 13 were previously classified as PS-dependent TDP-43 interactors in our AP-MS analysis (Fig. 3). Although the abundance changes for most endogenous proteins were very modest, proteins identified as PS-dependent interactors displayed a distinct trend: they were more abundant in the supernatant (more soluble) in presence of the PS-deficient TDP-43 variants compared to solid-like variants (Fig. 4E).

To validate that the observed co-depletion of endogenous proteins is a result of altered phase separation properties of TDP-43, we set up a dose-response experiment, reasoning that the co-pelleting/co-depletion of proteins should follow a dose-response curve and be dependent on the concentration of added TDP-43. Thus, we set up phase separation reactions with increasing amounts of TDP-43 WT and 5R and probed protein levels in the supernatant by mass spectrometry (Fig. 4F). In this dataset, we selected proteins that either showed a 5R-specific dose-response pattern or had a lower fitted area under the curve in 5R than in WT, indicating a stronger co-depletion with 5R than WT (see Supplementary Fig. 5A, B, C for quality control and methodology). We focused specifically on 5R due to its prominent solid-like condensation phenotype and lower C_sat_, enabling us to capture differentially co-condensing proteins more efficiently. Dose-response co-depletion profiling retrieved 403 proteins that showed enhanced co-pelleting with 5R variant compared to WT. Using this orthogonal approach, we validated ∼11% of the interactors that were bound with higher affinity to solid-like TDP-43 variants in our AP-MS experiment (22 proteins; *p*value = 5.18 × 10^−11^, hypergeometric test) (Fig. 4G, Supplementary Table 4). None of the PS-independent interactors were identified, and spliceosome components were the most enriched proteins co-pelleting with 5R condensates (Fig. 4H). The depletion profiles of proteins belonging to this cluster (DDX3X, FUBP3, HNRNPA3, HRNPR, HNRNPUL2), as well as of other nucleus-related proteins (NUP160 and SMARCA5) are reported in Supplementary Fig. 5D - J.

Several lines of evidence suggest that our co-condensation profiling experiment is a complementary interactomics approach to identify proteins interacting via phase separation. First, although a cell lysate is devoid of native cellular architecture, potentially allowing TDP-43 to interact with proteins it might not encounter under physiological conditions, we identified a functional association between TDP-43 and the proteins that co-precipitate with 5R condensates. The functional association (calculated as the rank position from a minimal cost path to TDP-43) is lower compared to the PS-dependent interactors but higher compared to a group of random proteins sampled from the identified proteins in the dataset (Fig. 4I). Second, interactors that co-partition into 5R condensates were enriched in proteins known to undergo phase separation (Fig. 4J and Supplementary Fig. 5K). Third, PS-dependent interactors and proteins co-condensing with 5R exhibit shared molecular sequence features, suggesting a common mechanism of interaction. We calculated 128 sequence parameters^95^ (see Materials and Methods) and evaluated whether these features were significantly enriched or depleted in the AP-MS or co-condensation datasets using the human proteome as a control reference (Supplementary Fig. 6). We focused on parameters that were significantly enriched or depleted in at least one of the two datasets and identified a modest yet significant correlation between them (*p*value = 0.04) (Fig. 4K). This correlation reflected a convergence of sequence features that underlie PS-dependent interactions and co-condensed proteins. Specifically, these proteins were enriched in acidic and basic amino acids resulting in highly unbalanced charged sequences (high FCR and sigma) typical of strong polyelectrolytes dominated by long-range repulsion (Fig. 4L). At the same time, they displayed reduced features that promote order, such as structured interaction propensity, hydrophobicity, proline content, aromaticity (tryptophan), and polar residues (serine, cysteine) (Fig. 4L).

Integration of these orthogonal approaches shows that AP-MS and co-condensation profiling converge on proteins that are functionally linked to TDP-43, in particular splicing regulators, and are enriched in phase-separating proteins with distinct sequence features. Our data demonstrate that these proteins selectively co-condense with solid-like TDP-43 condensates, thereby possibly causing their sequestration and alterations in functionality.

### The RNA-helicase UPF1 selectively co-condenses with solid-like TDP-43 variants

An interesting disease-linked protein that showed a stronger interaction with solid-like TDP-43 variants (Fig. 5A, Fig. 3) and co-condensed with 5R in a dose-dependent manner (Fig. 5B, Fig. 4), is UPF1. UPF1 is an RNA-dependent helicase involved in non-sense-mediated decay (NMD), which recently was identified as a TDP-43 interactor that becomes dysfunctional in ALS^16,128^. We excluded that UPF1 levels were influenced directly or indirectly by TDP-43, since we observed low variability of UPF1 protein and RNA levels across the KDs and cells expressing TDP-43 variants (Extended Data Fig. 5A, B). Thus, the enrichment of UPF1 with solid-like TDP-43 variants in the AP-MS experiment is unlikely to result from changes in UPF1 expression. To further validate the co-pelleting of UPF1 with solid-like TDP-43 condensates, we performed TDP-43 condensation experiments in the presence of a total HeLa cell lysate, followed by a pellet/supernatant fractionation. We compared TDP-43 variants with distinct phase separation properties and quantified the abundance of UPF1 in the supernatant and pellet fractions by western blotting. Pellet quantification confirmed that UPF1 showed enhanced co-condensation with all solid-like variants compared to the PS-deficient TDP-43 variants (Fig. 5C).

**Figure 5:**
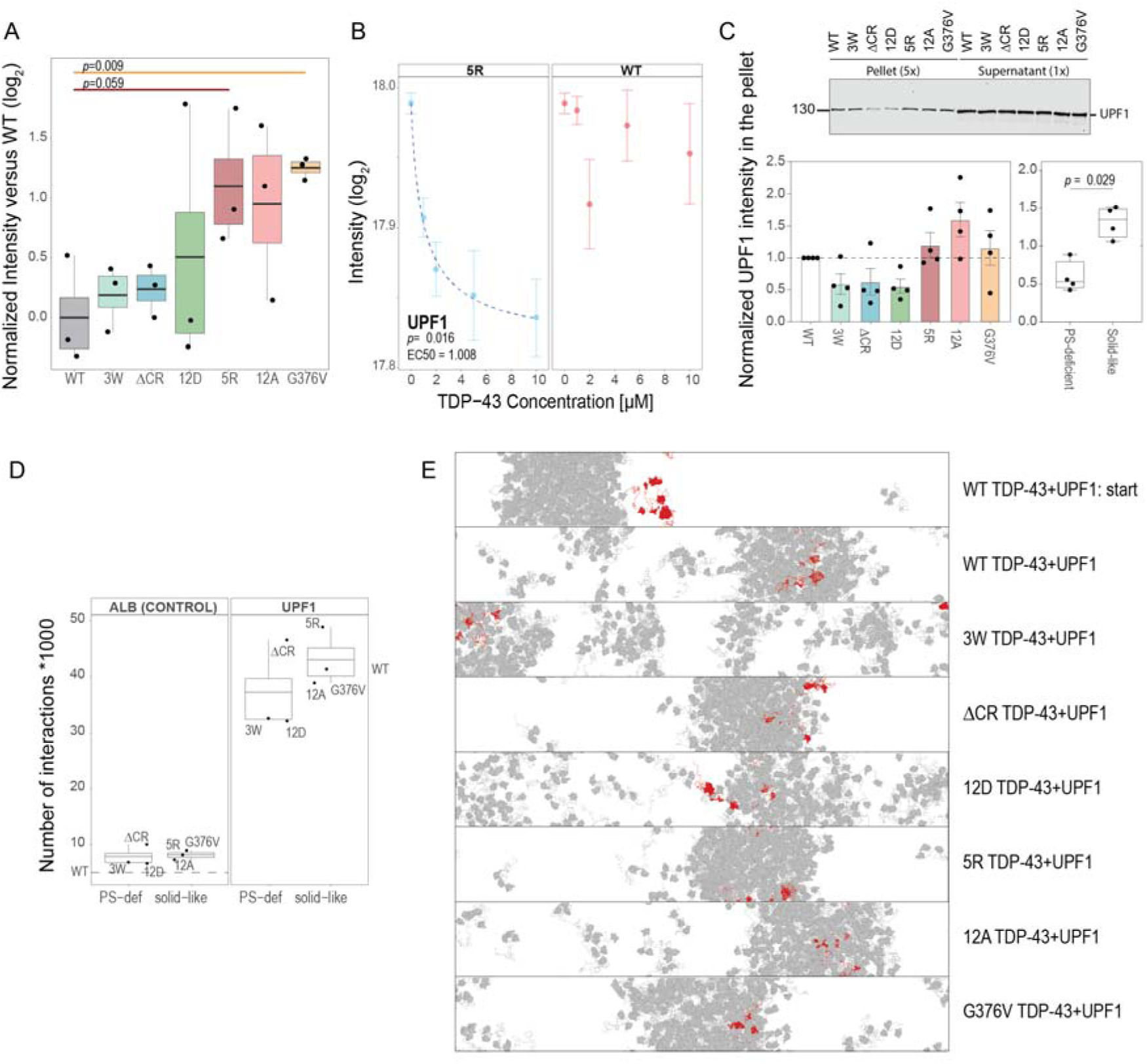
The RNA-helicase UPF1 selectively co-condenses with solid-like TDP-43 variants. 5A. UPF1 abundance in the AP-MS experiment across different TDP-43 variants. The level of UPF1 was normalized to the abundance of TDP-43 purified in each condition and the values are further normalized to the average of the WT condition. Single points represent biological replicates (N = 3), and the boxplot reports the distribution of the values with the mean (central line), the interquartile range (1^st^ and 3^rd^ quartile) and whiskers (1.5XIQR, interquartile range). 5B. UPF1 abundance in a dose-response co-condensation experiment with 5R (blue) and WT TDP-43 (red). UPF1 shows a dose-dependent depletion from the supernatant upon titration of 5R, but not WT TDP-43, into Hela cell lysate. Single points represent biological replicates (N = 4), and the dashed line indicates the fitted dose-response curve. 5C. Western blot illustrating the abundance of UPF1 pellet and supernatant fractions for the co-condensation experiment with TDP-43 variants (10 μM) in the presence of the HeLa cell lysate. The pellet fraction was loaded at five times the volume of the supernatant fraction. Lower panel: quantification of UPF1 in the pellet fraction was performed using four independent replicates, each derived from distinct batches of cell lysate. The values were normalized to set the abundance of UPF1 in TDP-43 WT sample to 1. Mean values shown. Error bars indicate standard error of the mean (SEM). Boxplot on the right illustrates mean values for PS-deficient and solid-like mutants across 4 replicates. Solid line indicates the median, whiskers indicate min and max values. Significance was determined using the Mann Whitney test. 5D. Number of interactions between 1 UPF1/albumin (CTRL) chain and 100 chains of TDP-43 variants, as calculated from MD simulations (averaged over 2 500 frames, 5 µs). Interaction counts are normalized considering protein length. 5E. Representative visualization of *in silico* condensates generated using simulations with the CALVADOS3 coarse-grained model. One chain of UPF1 is shown in red, while 100 chains of TDP-43 are shown in gray. The simulation was performed by adding one chain of UPF1 externally to a condensate generated from 100 molecules of TDP-43 (Start). The other snapshots show the last frame of the simulations (5 µs) for UPF1 with the condensates formed from TDP-43 WT, PS-deficient variants (3W, ΔCR and 12D) and solid-like variants (5R, 12A and G376V).

As a second approach, we analyzed co-condensation of UPF1 with the different TDP-43 variants *in silico*. For this purpose, we used MD simulations and modeled a TDP-43 condensate composed of 100 TDP-43 chains together with a single UPF1 chain or a control protein (i.e. albumin) placed outside of the condensate at the beginning of the simulation. This setup allowed us to assess whether UPF1 preferentially partitions into the condensate compared to the control protein and whether partitioning depends on the phase separation properties of TDP-43. UPF1 showed a higher number of interactions (Fig. 5D) and greater co-localization with TDP-43 density profiles than with albumin (Extended Data Fig. 5C). These interactions were particularly enhanced with solid-like TDP-43 condensates and with ΔCR, which in our MD simulations formed stable condensates (Fig. 1B; Fig. 5D, E). Overall, i*n silico* and biochemical experiments confirms that solid-like variants of TDP-43 selectively co-condense with UPF1, potentially sequestering it in less mobile assemblies.

### TDP-43 phase separation properties modulate alternative splicing and RNA levels

Since we identified many splicing regulators among the PS-dependent TDP-43 interactors and the proteins that selectively co-condensed with solid-like TDP-43, we assessed whether the phase separation propensity of TDP-43 impacts splicing regulation. To this end, we performed mRNA sequencing on RNA extracted from cell lines expressing the different TDP-43 variants. The experimental design included a non-targeting control (NTC), knockdown (KD) of endogenous TDP-43, and a rescue by inducible expression of TDP-43 variants in three different clones of each cell line (Fig. 6A, see Supplementary Fig. 7A for quality control). The rescue restored TDP-43 to slightly higher than endogenous levels but was comparable across the different TDP-43 variants and clones, as confirmed by mRNA levels and western blotting (Supplementary Fig. 7B, C). Due to TDP-43’s long-established role in regulating exon skipping and inclusion ^123,129–131^, we focused specifically on this group of splicing events. We considered splicing events with significant (FDR < 0.05) changes in percent spliced in (PSI) upon TDP-43 rescue compared to KD: a positive value (ΔPSI > 0.1) indicates increased exon inclusion in the rescue, while increased exon skipping is defined by a negative value (ΔPSI < −0.1) (Supplementary Table 5). The number of identified events varied across the analyzed TDP-43 variants from ∼250 (for 12A) to a maximum of ∼1 000 events (for 5R). No clear pattern of the number of splicing events was associated with the phase separation properties (PS-deficient and solid-like groups) (Supplementary Fig. 7D).

**Figure 6:**
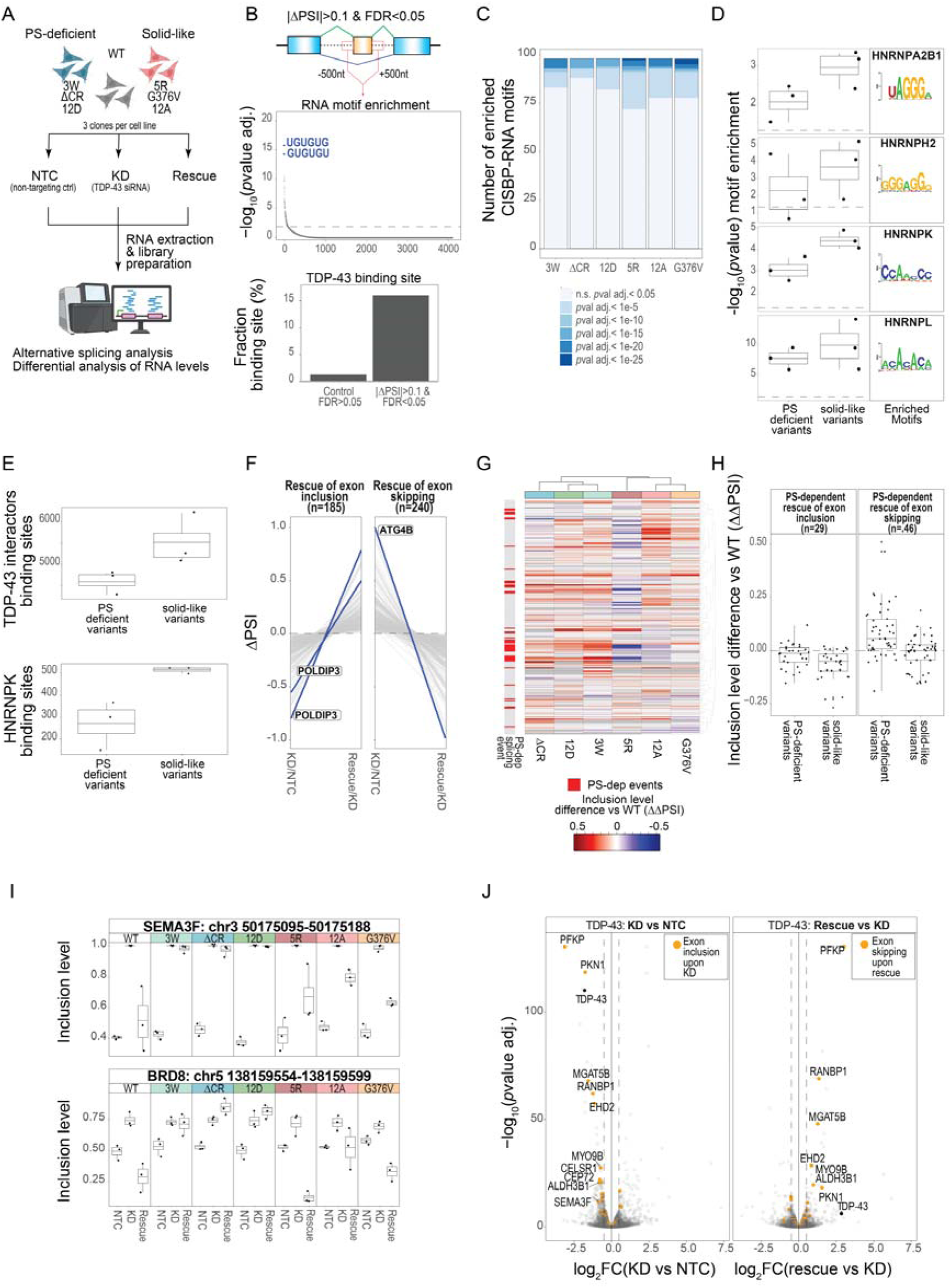
TDP-43 phase separation properties modulate alternative splicing and RNA levels. 6A. RNA sequencing experiment setup to study alternative splicing and gene expression changes in cell lines expressing different TDP-43 PS variants. Created in part with BioRender.com. 6B. Identification of RNA motifs and binding sites. Flanking regions (±500 nt) adjacent to alternatively spliced exons with significant splicing changes (|ΔPSI| > 0.1) in the rescue versus KD comparison were analyzed. Upper panel: enrichment analysis of all 4 096 possible 6-mers; significance was calculated using hypergeometric test with flanking regions of unchanged alternative exons as a background. Lower panel: TDP-43 binding sites present in the flanking regions of alternatively spliced exons (illustrated as percentage of the analyzed region). The analysis was performed using TDP-43 binding regions identified from ENCODE eCLIP datasets (see materials and methods) in flanking sequences of significantly and not significantly (control) alternatively spliced exons. 6C. Number of significantly enriched CISBP-RNA (*Homo sapiens*, DNA-encoded) motifs identified across sample conditions. Enrichment was calculated by hypergeometric test (*p*values adjusted) with flanking regions of unchanged alternative exons as a background. 6D. Significance enrichment (hypergeometric test) for RNA motif of proteins identified as PS-dependent TDP-43 interactors (HNRNPA2B1, HNRNPH2, HNRNPK, HNRNPL). The enrichment for each TDP-43 variant is shown as a single point. The boxplot reports the distribution of the values with the average (central line), the interquartile range (1^st^ and 3^rd^ quartile) and whiskers (1.5XIQR, interquartile range). The motif sequence logo (according to CISBP-RNA is shown in the right box. The RNA motif enrichment for each single TDP-43 variant is shown in Fig. S12F. 6E. Binding sites of TDP-43 interactors in flanking regions of alternatively spliced exons in PS-deficient and solid-like mutant groups. Upper panel: boxplot shows the number of RNA-binding sites of TDP-43 interactors identified within flanking regions (±500 nt) of regulated exons. 13 PS-dependent TDP-43 interactors were considered based on the overlap with the ENCODE eCLIP datasets: DDX3X, EFTUD2, ELAVL1, FASTKD2, FUBP3, HNRNPC, HNRNPK, HNRNPL, HNRNPM, HNRNPU, MATR3, PRPF8, and UPF1. Lower panel: number of HNRNPK binding sites identified within the same flanking regions as in the upper panel. 6F. TDP-43-dependent alternatively spliced exons. A total of 425 events were identified as WT rescue-responsive events (185 exon inclusion and 240 exon skipping). Only events that show opposite changes in the KD versus NTC and rescue versus KD comparisons in WT are considered. Two well-characterized alternative splice targets of TDP-43 (*POLDIP3* and *ATG4B*) are shown as examples. 6G. Unsupervised hierarchical clustering of 240 exon skipping events in the rescue versus KD comparison. Values reported in the heatmap show the differences in inclusion levels normalized to WT (ΔΔPSI), with positive values (weaker skipping compared to WT) in red and negative values (stronger in skipping compared to WT) in blue. In the annotation bar on the left, the events identified as PS-dependent are shown in red (events where the standard deviation within each of the PS groups is lower than the total standard deviation). The unsupervised hierarchical cluster for 185 exon inclusion events is displayed in Fig. S13F. 6H. Inclusion level differences in rescue vs KD comparisons in PS-dependent TDP-43 variants normalized to WT (ΔΔPSI). Individual dots indicate alternative splicing events. The boxplot reports the distribution of the values with the median (central line), the interquartile range (1^st^ and 3^rd^ quartile) and whiskers (1.5XIQR, interquartile range). 6I. Examples of PS-dependent alternatively spliced exons. Inclusion levels of alternatively spliced exons from *SEMA3F* and *BRD8* are shown under NTC, KD and rescue conditions. The levels for each replicate are shown as a single point, while the boxplot shows the average (central line) and the interquartile range (1^st^ and 3^rd^ quartile) and whiskers (1.5XIQR, interquartile range). 6J. Volcano plot showing RNA-level changes following TDP-43 knockdown (left) and subsequent rescue with TDP-43 WT (right). The x-axis represents the log_2_FC, while the y-axis indicates the significance of changes (*p*values adjusted). Exons included upon TDP-43 knockdown and exons skipped upon rescue with TDP-43 WT conditions are highlighted in orange.

We examined the sequence in the flanking regions (+/−500 nt) of the differentially alternatively spliced exons to identify if they are enriched in specific RNA motifs (Fig. 6B). The motif enrichment analysis (AME) ^104^ indicated that 6-mers UGUGUG and GUGUGU were the most enriched motifs, confirming the well-characterized binding preference of TDP-43 for UG-rich sequences, and showing that these motifs are prevalent in the vicinity of the identified alternative splicing events^129^ (Fig. 6B, upper panel). We also analyzed these regions for the presence of previously identified TDP-43 binding sites based on the eCLIP data available on ENCODE^106^ (the datasets are detailed in the materials and methods). When compared to exons without significant splicing changes (control), differentially alternatively spliced exons in rescue versus KD have a higher density of TDP-43 binding sites (Fig. 6B, lower panel).

Next, we wanted to test whether the flanking regions of exons regulated by the TDP-43 variants contain binding motifs for other RBPs as changes in enrichment of particular motifs could indicate altered binding of RBPs in a given region. We utilized the CISBP RNA motif database^105^ and identified that the intronic flanking regions of alternatively spliced exons regulated by sold-like TDP-43 variants were characterized by broader motif specificity (Supplementary Fig. 7E) and a higher number of significantly enriched motifs (Fig. 6C). This finding would be in line with enhanced interactions of certain RBPs with these solid-like TDP-43 variants. This prompted us to analyze the fingerprints of significantly enriched motifs for RBPs that we previously identified as PS-dependent TDP-43 interactors in our AP-MS experiment (Fig. 3E, F). Indeed, we identified motifs associated with HNRNPA2B1, HNRNPH2, HNRNPK and HNRNPL, all of which are implicated in splicing regulation. For three of these, we found a higher enrichment of their RNA binding motifs in the flanking regions regulated by solid-like TDP-43 variants (Fig. 6D, Supplementary Fig. 7F). To determine whether the enrichment of RNA-binding motifs translates into protein-RNA interactions, we analyzed eCLIP datasets from ENCODE^106^ (the datasets are detailed in the materials and methods) for identified PS-dependent interactors (13 proteins identified in the dataset). We found a higher number of eCLIP-enriched binding regions associated with alternatively spliced exons in the solid-like group compared to the PS-deficient group (Fig. 6E, upper panel). Notably, among the interactors identified by motif analysis, HNRNPK showed a markedly greater number of binding sites in the solid-like compared to the PS-deficient group (Fig. 6E, lower panel). Together, these findings suggest that solid-like TDP-43 variants influence alternative splicing by recruiting additional RBPs that contribute to splicing regulation.

Next, we analyzed alternative splicing events regulated by different TDP-43 variants for rescue-associated changes of exon inclusion (rescue versus KD, ΔPSI > 0.1) and exon skipping (rescue versus KD, ΔPSI < −0.1). The overlap of regulated events across conditions was limited, and no distinct group of events could be specifically attributed to the PS properties of different TDP-43 variants (Supplementary Fig. 7G, H). To address this limitation, we focused on events showing opposite changes in the KD versus NTC and rescue versus KD comparisons in WT, meaning that the rescue with WT significantly reversed the PSI changes induced by the KD towards the levels observed in NTC. This approach identified 425 WT TDP-43 rescue-responsive events, composed of 185 exon inclusion and 240 exon skipping events (Fig. 6F). Motif enrichment analysis of their flanking regions revealed significant enrichment of UGUGUG and GUGUGU motifs (Extended Data Fig. 6A). Among the rescue-responsive events, we identified several well-characterized TDP-43 splicing targets, including *AGRN*^132,133^, *ATG4B*^132,134^, *MADD*^129,135,136^, *PFKP*^132,137^, *RANBP1*^132^, *BRD8*^135,138^ and *POLDIP3*^131,137,139^ (see Extended Data Fig. 6B, C for Sashimi plots of alternative splicing events in *ATG4B* and *POLDIP3*). Most TDP-43-dependent alternatively spliced exon skipping/inclusion events were located in protein-coding regions, with fewer events occurring in long non-coding RNAs or 5’UTRs (Extended Data Fig. 6D).

Among the 425 rescue-responsive events, we identified 75 PS-dependent alternatively spliced exons. We defined PS-dependent alternatively spliced exons as those characterized by consistent inclusion patterns within each variant group. For this we filtered events whose inclusion level differences displayed lower variability within each one of the PS-defined groups (PS-deficient and solid-like) compared to the variability observed across all experimental conditions. A summary of the filtering strategy is provided in Extended Data Fig. 6E. Hierarchical clustering of ΔΔPSI values (comparison of ΔPSI in variants versus WT) for events associated with exon skipping upon rescue revealed two distinct groups, corresponding to the PS properties of the TDP-43 variants (Fig. 6G). In contrast, when analyzing events that exhibited exon inclusion upon rescue, we identified fewer PS-dependent events along with a less distinct clustering between the two phase separation groups (Extended Data Fig. 6F). PS-dependent splicing events showed higher levels of exon inclusion upon rescue in the PS-deficient group compared to the solid-like group. This effect was most evident in events with exon skipping upon rescue, where PS-deficient variants were less effective at restoring splicing, whereas solid-like variants performed closer to WT levels (Fig. 6H). Among the PS-dependent alternatively spliced exons, we identified several previously reported TDP-43 splice targets, including *MADD*^135^*, SEMA3F*^123^, *BRD8*^135^, and *PILRB*^129^ (inclusion levels and Sashimi plots for *BRD8*, and *SEMA3F* are shown in Fig. 6I and Extended Data Fig. 6G,H; ΔΔPSI values for rescue versus KD in all PS-dependent alternative splicing events are reported in Extended Data Fig. 6I). In summary, these findings highlight a significant role of TDP-43 phase separation in regulating a subset of alternative splicing events.

Next, we investigated whether TDP-43 alternative splicing affects overall RNA abundance (Supplementary Table 6). Exon inclusion or exclusion can disrupt the reading frame, potentially introducing premature termination codons that trigger RNA degradation. Differential analysis of RNA levels under WT conditions, comparing KD to NTC and rescue to KD, revealed that among the genes with the most significant changes, many harbor TDP-43 rescue-responsive alternative splicing events which we previously identified (Fig. 6F). This phenomenon was specifically observed for exons that were included upon TDP-43 knockdown and skipped upon rescue with WT (Fig. 6J, Extended Data Fig. 6J), pinpointing the role of TDP-43 in the suppression of non-conserved exons^132^.

To verify whether RNA abundance changes could be attributed to altered TDP-43 phase separation properties, we identified transcripts with at least two variants per group (solid-like and PS-deficient) that were coherently and significantly up- or downregulated when comparing rescue to TDP-43 KD. Subsequently, we filtered transcripts for which the standard deviation of rescue versus KD differences within each of the PS-defined groups (PS-deficient and solid-like) was lower than the global standard deviation. This led to the identification of 87 genes whose RNA levels are modulated by phase separation (Extended Data Fig. 6K). Most PS-dependent transcripts map to protein-coding genes (Extended Data Fig. 6L), and upregulated PS-dependent genes are linked to protein quality control pathway and proteostasis (Extended Data Fig. 6M).

Collectively, RNA profiling in cells expressing TDP-43 variants with distinct PS properties reveals differences in RNA abundance and splicing between PS-deficient and solid-like states, establishing phase separation of TDP-43 as an important modulator of its RNA regulatory function. Additionally, our data suggest that TDP-43 PS-mediated interactions with other splicing factors such as HNRNPK, may in part be responsible for the observed PS-dependent changes in alternative splicing.

### TDP-43 phase separation regulates the cellular proteome directly and indirectly

After establishing TDP-43 PS-dependent regulation of alternative splicing and RNA-levels, we investigated whether TDP-43 phase separation impacts the regulation of the cellular proteome. To address this, we performed proteome profiling analysis to evaluate the effects of endogenous TDP-43 depletion, followed by re-expression of myc-tagged TDP-43 WT or the different TDP-43 PS variants (see scheme in Supplementary Fig. 8A). By integrating RNA and proteome datasets, our goal was to define the functional role of TDP-43 phase separation in regulating alternatively spliced exons, RNA and protein levels, as well as to explore the potential contribution of PS-dependent interactors in the regulation of protein levels (Fig. 7A, Supplementary Table 7).

**Figure 7:**
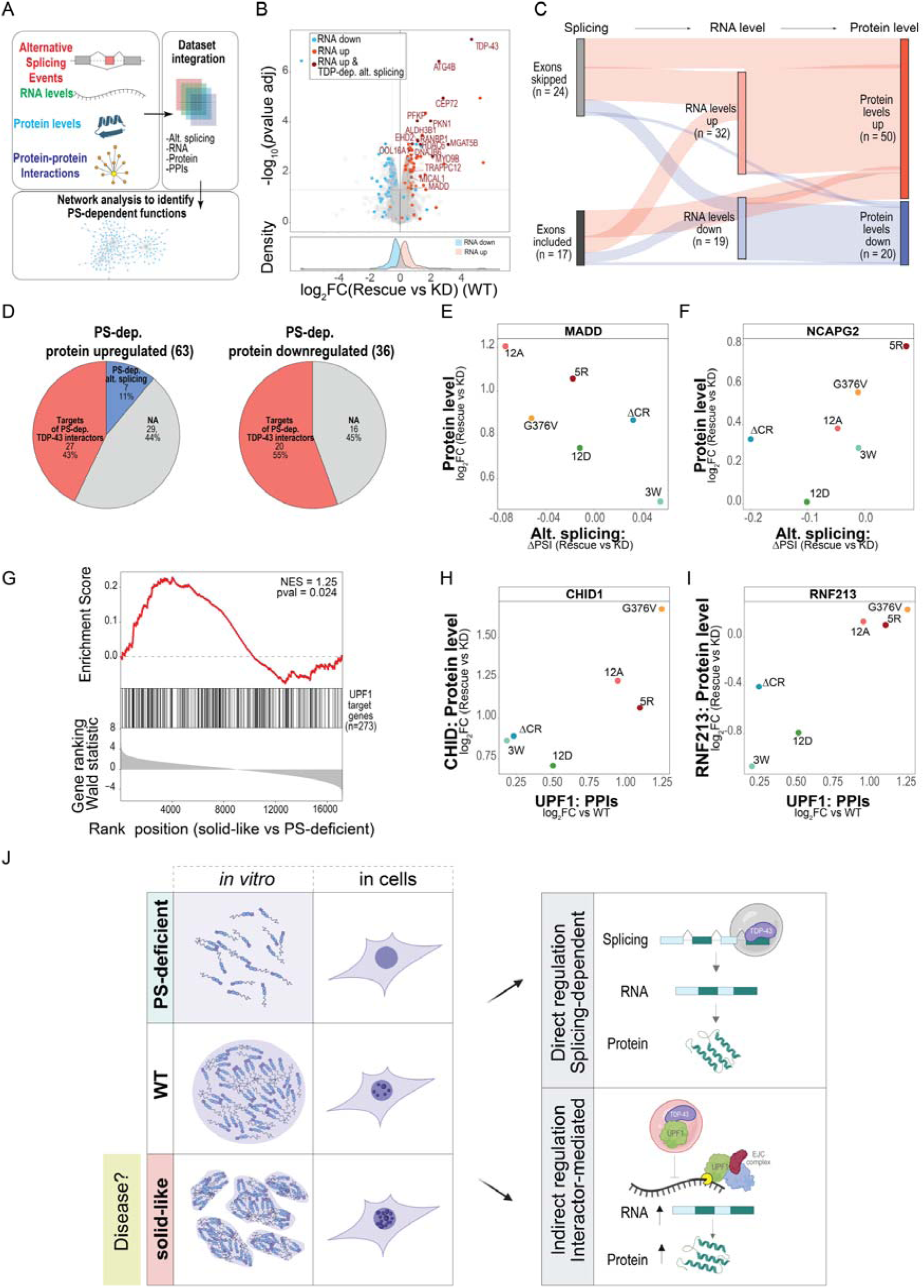
TDP-43 phase separation regulates the cellular proteome directly and indirectly. 7A. Workflow for the identification of PS-dependent events across multiple levels of gene regulation, including alternative splicing, RNA abundance, protein levels, and protein-protein interactions. PS-dependent events were filtered independently in each dataset and subsequently integrated through network analysis to identify regulatory links. Created in part with BioRender.com. 7B. Volcano plot showing differences in protein abundance between TDP-43 knockdown (KD) and rescue with TDP-43 WT. The x-axis represents differential protein abundance, while the y-axis shows statistical significance (adjusted *p*value). Proteins associated with TDP-43-regulated alternative splicing are highlighted in dark red, while genes identified in the RNA-seq dataset as up- or downregulated upon rescue are highlighted in red and cyan. The lower panel shows the density distribution of protein abundance changes for genes that are significantly differentially regulated at the RNA level (|og2FC| > 0.3, adjusted *p*value < 0.05). 7C. Flow diagram displaying events that change upon TDP-43 depletion. The first level represents alternative splicing changes (exon inclusion and skipping upon rescue with WT), the second and third levels represent RNA and protein changes, respectively. The color of the links indicates whether an event is associated with protein up-(red) or downregulation (blue) in rescue versus KD. 7D. Summary of the PS-dependent effects on protein abundance. Proteins linked to PS-dependent alternative splicing events are shown in blue. Proteins associated with PS-dependent TDP-43 interactors are shown in red. Proteins for which no mechanistic link could be established are shown in gray. 7E, F. Impact of TDP-43 PS properties on alternative splicing and protein levels. Correlation plots showing coordinated changes in protein abundance and alternative splicing levels for MADD and NCAPG2 across TDP-43 variants. In the case of MADD, solid-like TDP-43 variants exhibit reduced exon inclusion, which correlates with increased protein abundance. Conversely, for NCAPG2, reduced exon inclusion is associated with decreased protein abundance. 7G. Gene set enrichment analysis (GSEA) comparing RNA levels for solid-like TDP-43 variants (5R, 12A and G376V) and PS-deficient variants (3W, ΔCR and 12D). Gene set: UPF1 target genes in HEK293 and HeLa cells^108–110^. The y-axis displays the running enrichment score (ES) (upper panel), the position of UPF1 target genes (middle panel) and corresponding ranked changes calculated with Wald statistic (lower panel); the x-axis shows the ranked position of each gene in the set. NES (normalized enriched score) and *p*value are shown on the plot. 7H, I. Indirect impact of TDP-43 PS properties on TDP-43 interactors and protein levels. Correlation plots showing coordinated changes in abundance of UPF1 and its targets (CHID1 and RNF213). Solid-like TDP-43 variants sequester mRNA decay factor UPF1, possibly reducing its availability for the RNA degradation machinery and leading to increased protein levels of UPF1 target genes. 7J. Distinct mechanisms though which TDP-43 phase separation regulates the proteome. Alterations in TDP-43 phase separation, which may occur in disease, can have different regulatory effects: (1) a direct effect, spicing-dependent effect, where TDP-43 phase separation modulates alternative splicing of TDP-43 target genes and corresponding protein levels, and (2) an indirect effect, where TDP-43 phase separation changes interactions with RNA regulatory factors (e.g. UPF1 shown as example), which in turn regulate RNA and protein abundance. Created in part with BioRender.com.

We analyzed the abundance levels of 8 694 proteins (filtering steps for the data analysis are shown in Supplementary Fig. 8B), with a good coverage of the proteome dynamic range spanning from hundred pM to µM (according to OpenCell^140^) (Supplementary Fig. 8C, quality control assessments for the proteomics dataset are shown in Supplementary Fig. 8D, E, F). Upon knockdown, TDP-43 protein levels are, on average, 5-fold lower (log_2_ scale) compared to the rescue conditions, which displayed relatively consistent re-expression levels across the different variants (see Supplementary Fig. 8G, H for TDP-43 quantification by mass spectrometry and western blotting).

We first analyzed TDP-43 WT to evaluate the impact of TDP-43 KD and rescue on protein expression levels. After the rescue, 289 proteins were significantly up- or downregulated compared to WT. This finding aligns with the crucial role of TDP-43 in RNA metabolism and gene regulation of hundreds of proteins^141^, including TDP-43 itself^123,142^. The most pronounced changes in protein levels occurred for genes harboring alternatively spliced exons. Notably, genes showing differential regulation at the RNA level also exhibited corresponding changes at the protein levels, in line with observations made by *Kozareva et al.*^141^ (Fig. 7B). This data indicates a robust correlation between RNA-level regulation and protein-level changes (Supplementary Fig. 8I), reinforcing the interconnected relationship between the levels of TDP-43, alternative splicing, RNA expression and protein abundance. An integrated analysis of these layers is illustrated in a flow diagram, which connects alternative splicing to RNA and protein expression levels (Fig. 7C). Notably, most of the exon skipping events upon rescue are linked to elevated RNA and protein levels, underscoring the critical role of TDP-43 in regulating non-conserved exons, which in turn affects protein levels^132^. Overall, 50 out of 172 upregulated proteins were associated with splicing events or RNA changes (Fig. 7C), highlighting the profound influence of TDP-43-dependent RNA regulation on shaping the proteome.

As a next step we sought to investigate how the phase separation properties of TDP-43 influence proteome regulation. We performed a differential analysis comparing protein levels in the rescue versus KD conditions across all analyzed TDP-43 PS variants (Extended Data Fig. 7A). Proteins significantly and coherently regulated in at least two variants per group were classified as PS-dependent if their signal variability within each one of the groups (solid-like and PS-deficient) was lower than the overall variability. As expected, the changes driven by phase separation were more subtle compared to those observed during TDP-43 rescue or knockdown experiments and we identified 63 upregulated and 36 downregulated proteins influenced by phase separation (Fig. 7D). Analysis of relative protein abundance across conditions revealed higher levels of proteins in cells expressing solid-like TDP-43 variants compared to PS-deficient variants (Extended Data Fig. 8), highlighting the role of TDP-43 phase separation in shaping the proteome composition. Among the upregulated proteins, those with increased abundance in solid-like variants were enriched for biological processes related to proteostasis, indicating that solid-like TDP-43 variants might induce an autophagy response (Extended Data Fig. 7B). Conversely, downregulated proteins were enriched for processes involved in axon guidance and neural circuit organization, including the semaphorin-plexin signaling pathway (Extended Data Fig. 7C). Extended Data Fig. 8 provides a detailed overview of the identified PS-dependent proteins, their relative abundance, and their connections to PS-dependent alternative splicing and RNA regulation.

Next, we investigated whether changes in PS-dependent protein abundance are linked to alternative splicing and changes in RNA levels. We identified 7 PS-dependent alternative splicing events (as defined in the previous section) occurring in genes coding for proteins whose levels are PS-dependent. Notably, almost all of these events (6) involve exon skipping upon rescue, which is associated with an increase in protein levels. We define these as proteins regulated by PS-dependent alternative splicing. Fig. 7, E, F illustrates two examples where PS-dependent alternative splicing results in PS-dependent protein level changes (increased and decreased abundance of MADD and NCAPG2 respectively).

However, only a small fraction of the PS-regulated proteins can be explained by phase separation-dependent alternative splicing (Fig. 7D).To address the origin of the remaining PS-dependent protein abundance changes, we followed an approach recently proposed by *Kozareva et al.*^141^ which introduced a *“secondary cascade”* model demonstrating that the mis-regulation of a few genes upon TDP-43 knockdown leads to widespread changes on the proteome level. As our AP-MS dataset revealed that many of the TDP-43 PS-dependent interactions are RGG motif-containing RBPs with roles in gene regulation, we investigated whether there is a functional relationship between these interactors and proteins regulated in a PS-dependent manner. To this end, we integrated the data from RNAInter^112^, a database that catalogs protein-RNA binding interactions. This analysis revealed that the majority of proteins differentially regulated by TDP-43 phase separation are potentially functional targets of PS-dependent TDP-43 interactors (Fig. 7D, Supplementary Fig. 9A, B reports functional links between PS-dependent TDP-43 interactors and PS-dependent differentially expressed proteins). We define these as proteins indirectly regulated by phase separation through TDP-43 interactors. The network in Extended Data Fig. 9, summarizes the organization of PS-dependent proteins showing how these levels are regulated by TDP-43 phase separation both by alternative splicing and indirectly by protein interactions.

Two additional observations support the proposed indirect regulation by TDP-43 PS-dependent interactors. First, RNA motif and binding analysis in the flanking regions of alternatively spliced exons revealed binding signatures of PS-dependent TDP-43 interactors (e.g., HNRNPK, HNRNPH2, HNRNPA2B1) (Fig. 6D). Second, we observed potential downstream effects of altered TDP-43 phase separation on the functionality of its interactors, as exemplified by UPF1. UPF1 is an RNA-dependent helicase involved in mRNA degradation which showed enhanced interaction and co-condensation with solid-like TDP-43 variants in different, orthogonal assays (Fig. 5). We identified UPF1 RNA targets from published studies in HEK293 and HeLa cells^108–110^, focusing on 310 genes reported in at least two independent studies (Extended Data Fig. 7D). Gene set enrichment analysis (GSEA) confirmed that UPF1 target genes are upregulated upon TDP-43 depletion in our cellular system (Extended Data Fig. 7E), in line with recently published data^16^. Next, we examined whether these genes exhibit enrichment, depletion, or no changes when comparing cells expressing solid-like TDP-43 variants to PS-deficient variants. GSEA analysis using UPF1 target genes revealed significant enrichment of UPF1 targets in cells expressing solid-like TDP-43 variants (Fig. 7G). For some UPF1 targets (e.g. CHID1 and RNF213) we observed a correlation between the PS-dependent UPF1 binding to TDP-43, and increased protein levels of CHID1 and RNF213 (Fig. 7H, I). These observations suggest that UPF1 might get trapped in solid-like nuclear TDP-43 condensates, potentially limiting UPF1 cytoplasmic pool and impairing its ability to efficiently perform RNA degradation. This result in increased abundance of UPF1-dependent transcripts and corresponding proteins (Fig. 7J).

Together, our observations indicate that RNA and protein levels are regulated by TDP-43 phase separation properties and reflect a direct effect of TDP-43 on alternative splicing of certain mRNA targets, and the indirect contribution of RNA-binding proteins whose function is modulated by the phase separation properties of TDP-43 (Fig. 7J).

## Discussion

### Tuning TDP-43 phase separation with diverse LCD mutations allows probing condensation-state-dependent functional changes in cells

While the role of phase separation in TDP-43 aggregation in neurodegenerative disease has been widely studied^21,37,143,144^, the relevance of TDP-43 condensation in supporting the protein’s physiological functions in the cell is only beginning to be investigated. As phase separation of TDP-43 might be altered in disease, and targeting its phase transitions is even explored as a therapeutic strategy^20,144–146^, it is of great importance to understand how and to what extent TDP-43 condensation plays a role for its functionality in RNA processing and global RNA/protein homeostasis.

Some efforts have already been made to address the functional relevance of TDP-43 condensation: One study deleted or mutated the alpha-helical conserved region (CR) in TDP-43, which drives TDP-43 condensation^42,43,60^, and showed that these CR perturbations alter TDP-43’s global RNA-binding profile, its autoregulation, and certain TDP-43-mediated 3’-end processing events^60^. A similar approach was adopted in mice, whereby deletion of the CR from the mouse TDP-43 was shown to cause neuronal and behavioral impairments and to enhance translation globally via an increased association of TDP-43 with the translation machinery^62^. Other studies perturbed not only the CR, but also other condensation-relevant sequences in TDP-43^45,61,115^. However, those studies only investigated the effects on a small number of alternative splicing events and either came to discrepant conclusions^45,115^, or confirmed that TDP-43 phase separation is necessary for TDP-43 autoregulation^61^.

In our work, we took a mutation approach to create variants of TDP-43 with PS-deficiency or solid-like condensate properties (Fig. 1A). We made sure to target different regions of the LCD and to utilize different types of mutations to exclude region/mutation-specific effects. Utilizing this panel of TDP-43 variants, where each condensation state is represented by three mutants, we were able to search for PS-dependent rather than sequence-dependent changes in TDP-43 functionality, making it likely that the effects we see come from changes to the condensation state and not the mutations themselves.

Our *in vitro* characterization of the chosen TDP-43 variants revealed two distinct behaviors: PS-deficient phenotypes for the 3W, ΔCR, and 12D variants, and more solid-like properties for the 5R, 12A, and G376V variants (Fig. 1). These findings align with previously published observations on the “molecular grammar” of TDP-43 phase separation, which has highlighted an important role of aromatic and aliphatic residues^44–46,113^ and the alpha-helical conserved region^42,43,60,147^. Our findings furthermore support the notion that charge plays a key role in regulating TDP-43 phase separation, as negatively charged residues in the C-terminus (12D) reduce condensation^71^, while positively charged arginine residues reduce condensate dynamicity^45^. Finally, the highly liquid-like phenotype of 3W (tryptophans substituted by glycines) and the more solid-like behavior of G376V are in line with glycine residues having a liquefying effect, consistent with previous observations in the FET proteins^73^ and glycine substitutions in TDP-43’s alpha-helical conserved region^43^.

Importantly, we observed that the PS behavior observed in homotypic PS assays *in vitro* (e.g. number of condensates, dynamics as measured by FRAP) was largely recapitulated when the same variants were stably expressed in cells at physiological levels and in the absence of endogenous TDP-43 (Fig. 2). This is not always the case for other protein systems, as a previous study reported discrepancies between the *in vitro* PS behavior of the chromosomal passenger complex and its behavior in cells or cytomimetic media^148^. In our system, TDP-43 variants retained their PS behavior both in cells and cellular lysates, supporting the notion that TDP-43 robustly forms condensates under physiological conditions. In healthy cells, where the protein is predominantly nuclear, TDP-43 enriches in multiple nuclear foci. Based on earlier studies, these might represent Cajal bodies, gems and paraspeckles, but also other assemblies of unknown identity^40,116,118^. To what extent the TDP-43 foci of undetermined identity overlap with transcriptional or splicing condensates^58,149^ and are functionally linked to the regulation of transcription and splicing remains to be investigated.

### TDP-43 phase separation modulates protein interactions, e.g. with RNA-binding proteins

Interestingly we found that protein interactome rearrangements were associated with distinct phase separation properties: solid-like TDP-43 variants exhibit enhanced interactions with numerous other proteins, particularly RNA-binding proteins (RBPs) and nuclear pore complex components (Fig. 3). This finding was validated in experiments with recombinant proteins in cellular lysates (Fig. 4). Inspired by previous studies that reconstituted condensates in cellular lysates^80,150^, we developed an experimental framework that enables the characterization of biophysical properties through microscopy as well as the identification of other cellular condensate components by mass spectrometry. A key advantage of this system is that it closely mimics cellular conditions, providing a near-physiological, multicomponent context, yet it allows for easy tuning of experimental conditions (e.g. variation of protein concentration, salt, pH, PTMs etc.) in a way that is normally only feasible in *in vitro* experiments.

Our interactomics dataset corroborates previously reported TDP-43 interactors^20,31–33,71,80–82^ and reveals how the interactome is organized into functionally coherent protein modules^120^ (Fig. 3E). The TDP-43 interactome is highly promiscuous, and TDP-43 ranks among the top 3% of proteins with the most extensive interaction network (BioGRID v. 5.0.253^83^). Its composition varies significantly based on subcellular localization and physiological or pathological conditions^25,151^. Key proteostasis-related proteins, such as HSP70, HSPB1, and co-chaperones BAG2 and BAG3, regulate TDP-43 condensation and inclusion formation^152–154^. Additionally, other RBPs, such as SRRM2, NUFIP2, and HNRNPC, modulate TDP-43 localization and splicing activity, hence they have been suggested as potential therapeutic targets for modulating TDP-43 function^128^. Our work demonstrates that the TDP-43 interactome is not solely dictated by cellular context but is also governed by its PS properties, given that all variants are functionally active and localized in the nucleus. This suggests that phase separation is a key mechanism through which TDP-43 interacts with specific partners to execute distinct functional activities. The interactomics dataset shows that several RBPs involved in splicing regulation interact with TDP-43 in a PS-dependent manner. This observation prompted us to investigate whether the expression of TDP-43 variants with distinct PS properties could lead to global changes in splicing regulation (Fig. 6). Indeed, we found a subset of splicing events that depended on TDP-43’s ability to phase separate or were altered upon expression of solid-like and PS-deficient TDP-43 variants (Fig. 6G). Previous studies largely used specific gene reporters to assess the role of PS in splicing and lack consensus on whether disease-linked or phase separation-altering mutations in TDP-43 affect its regulatory splicing activity^45,71,115^. A possible explanation has been proposed by *Hallegger et al.*^60^, by demonstrating that the effects of TDP-43 condensation are context-specific to particular RNA binding regions (unusually long clusters of motifs of characteristic types and density). While we cannot rule out that there are indeed context-dependent RNA binding preferences among our PS variants, we observed a potential indirect effect of TDP-43 PS on splicing regulation, which we linked to PS-dependent protein interactors. Analysis of sequences flanking alternatively spliced exons showed that exons differentially spliced by solid-like TDP-43 are enriched for RNA-binding motifs and binding sites associated with PS-dependent RBP interactors, particularly HNRNP proteins that interact more strongly with solid-like TDP-43 (Fig. 6C, D, E).This pinpoints that the recruitment of other RBPs into TDP-43 condensates may modulate splicing events, which we further linked to abundance changes on the protein level. Thus, a significant proportion of the splicing events regulated by solid-like TDP-43 variants might be driven by their altered interactomes.

Additionally, we observed that the PS-deficient TDP-43 variants showed a lower efficiency in correcting some of the alternative splicing changes (e.g. in *SEMA3F* or *BRD8*) (Fig. 6H, I). This could be attributed to a partial loss of interactors by these variants as we have observed in the AP-MS experiment, which might lead to deleterious effects on splicing of specific genes by these TDP-43 variants. Additionally, as PS-deficiency has been reported to compromise TDP-43 binding to long RNA-binding regions of specific motifs^155^, inability of our PS-deficient variants to correct splicing of a subset of genes could potentially be attributed to the specific sequence features of those transcripts.

### Phase separation modulates TDP-43 functional activity by alternative splicing and by re-arranging the interactome

Functional changes within the cell depend on the modulation of the cellular proteome at various levels, including alterations in protein abundance, reshaping of the interactome, and post-translational modifications^156^. In this study, we demonstrate that phase separation of TDP-43 is a critical mechanism that governs the reorganization of the cellular proteome. To uncover the global role of TDP-43 PS, we employed a multi-layered approach^141,157^ combining interactomics, RNA-Seq and proteomics data. By integrating these results, we describe how altered TDP-43 condensation, which might occur in diseases such as ALS or FTD, regulates alternative splicing, RNA and protein levels. Notably and not surprisingly, the effects observed in our study are more subtle and nuanced compared to those observed when studying the consequences of TDP-43 depletion^141^. However, our findings nicely complement the observation of a “cascade effect” reported upon TDP-43 knockdown, with almost half of the differentially expressed proteins being functional linked to PS-dependent interactors. In our study, we find that TDP-43 PS can regulate the splicing and abundance of RNAs, and consequently the proteome, both via PS-dependent alternative splicing of TDP-43 target genes, and indirectly via protein interactors that are modulated by TDP-43 PS properties (Fig. 7J). Our data reveal that numerous RNA-and protein-level changes arise when TDP-43’s PS behavior is altered, as might be the case in disease, for example through disease-linked mutations. We speculate that disease-relevant transcriptome and proteome rearrangements might be driven through alterations in TDP-43’s PS state and could begin long before TDP-43 mislocalization and the associated nuclear loss of function.

A notable example of the indirect phase separation-mediated regulation is UPF1. While UPF1 has been reported to be neuroprotective in models of TDP-43 dysfunction^158,159^, a more direct pathological link has been identified recently as TDP-43 and UPF1 were found to interact and co-aggregate in disease^16^. UPF1 functions in non-sense-mediated decay (NMD) and a set of UPF1-dependent mRNA targets normally undergo degradation via NMD^108–110^. Our data show that solid-like TDP-43 variants interact more strongly with UPF1, and UPF1 co-condenses with solid-like TDP-43 variants. Sequestration of UPF1 into undynamic TDP-43 condensates may compromise UPF1 function, in line with the observed increase in the levels of UPF1 RNA targets specifically in cell lines expressing solid-like PS variants. We exemplify this with CHID1 and RNF213, two UPF1 targets whose RNA and protein abundance linearly correlate with the abundance of UPF1 interacting with TDP-43.

Taken together, our study offers evidence that the cellular proteome is regulated via TDP-43 phase separation through the combination of two mechanisms. These comprise TDP-43-mediated alternative splicing as well as an indirect regulation involving global TDP-43 interactome rearrangements caused by altered TDP-43 PS properties.

### Limitations of the study

Our PS-dependent protein-protein interactions are based on AP-MS and co-condensation assays, which both were performed on cell lysates. These approaches, unlike proximity-based labeling approaches that preserve cellular architecture, cannot capture transient interactions in living cells or identify which interactions are specific to the native cellular environment, e.g. nucleus versus cytoplasm.

The integration of large-scale datasets is inherently constrained by differences in sensitivity, variability, definition of thresholds and data quality, which can introduce bias into multi-layer analyses. For instance, changes in protein levels may not be detectable for certain alternative splicing events (e.g., SEMA3F). These challenges are further exacerbated by additional regulatory mechanisms that TDP-43 is involved in that were not accounted for in the analysis. Specifically, we did not consider PS-dependent regulation of alternative polyadenylation at the 3′UTR, which can influence mRNA stability^60^ or PS-dependent regulation of mRNA translation^62^. More in-depth analysis of these individual TDP-43 functions would be needed to define the exact molecular mechanisms by which TDP-43 PS regulates the proteome.

Additionally, the datasets acquired are high-dimensional, characterized by a large number of variables but relatively few observations (6 analyzed TDP-43 variants). This imbalance can limit statistical power, make it more challenging to identify consistent patterns, and increase the possibility of correlations that may not accurately represent causal relationships.

Although we made every effort to control the levels of rescued TDP-43 across all our experiments, it was not always possible to achieve identical protein abundance. This variability added complexity to distinguishing the specific effects of phase separation from those related to TDP-43 abundance.

Finally, our analysis was performed in HeLa cells, which do not reflect the gene expression, RNA splicing and degradation patterns or regulatory dynamics of neuronal cells^160–162^. Consequently, our analysis is limited to studying TDP-43 phase separation effects on gene expression in general, without addressing neuron-specific processes. Future studies should validate these findings in neuronal models to better understand the physiological relevance of TDP-43 phase separation in neuronal gene regulation.

## Acknowledgements

The authors thank all members of the Dormann lab (in particular Emre Pekbilir and Sára Varga), Julian König and Christian Münch for critical discussion. We thank all lab members and Marc-David Ruepp for comments on the manuscript. We acknowledge the IMB Flow Cytometry Core Facility, especially Stefanie Möckel and Stephanie Nick, for their assistance with experiment setup and analysis. Cell sorting was performed on the Invitrogen Bigfoot (funded by the Deutsche Forschungsgemeinschaft (DFG), project number 511658729). Support by the Microscopy, Genomics, Bioinformatics, Proteomics and Protein Production Core Facilities of IMB Mainz is gratefully acknowledged. The authors acknowledge Fridolin Kielisch for the consulting on statistical analysis. The authors especially thank the Proteomics Core Facility for assistance with MS experiments and the Microscopy Core Facility for expert advice in imaging setup and analyses. We also acknowledge the internal service projects Z01 (Biopolymer Engineering and Bioanalytics) and Z02 (Advanced Biopolymer Imaging) of CRC1551 for their support. Funding by the DFG supported the Revvity Opera Phenix (project ID 316215830), the Leica Stellaris 8 FALCON (project ID 497669232) and the ORBITRAP ASTRAL (project ID 524805621). Funding by ReAlity (Landesinitiative des Landes Rheinland-Pfalz) supported purchase of the ZEISS Axio Observer wide field microscope.

This work was funded by the European Union (EU) through the ERC CoG TDP-ASSEMBLY (Project 1011258709), the DFG within the Heisenberg Programme (project ID 442698351), the CRC1551 “Polymer Concepts in Cellular Function” (project ID 464588647) and the SPP2191 “Molecular Mechanisms of Functional Phase Separation (project ID 419139133). F.U. acknowledges funding by a Marie Curie Individual Fellowship (Grant agreement number 101207537)).

## Extended Data Figures

**Extended Data Fig. 1.**
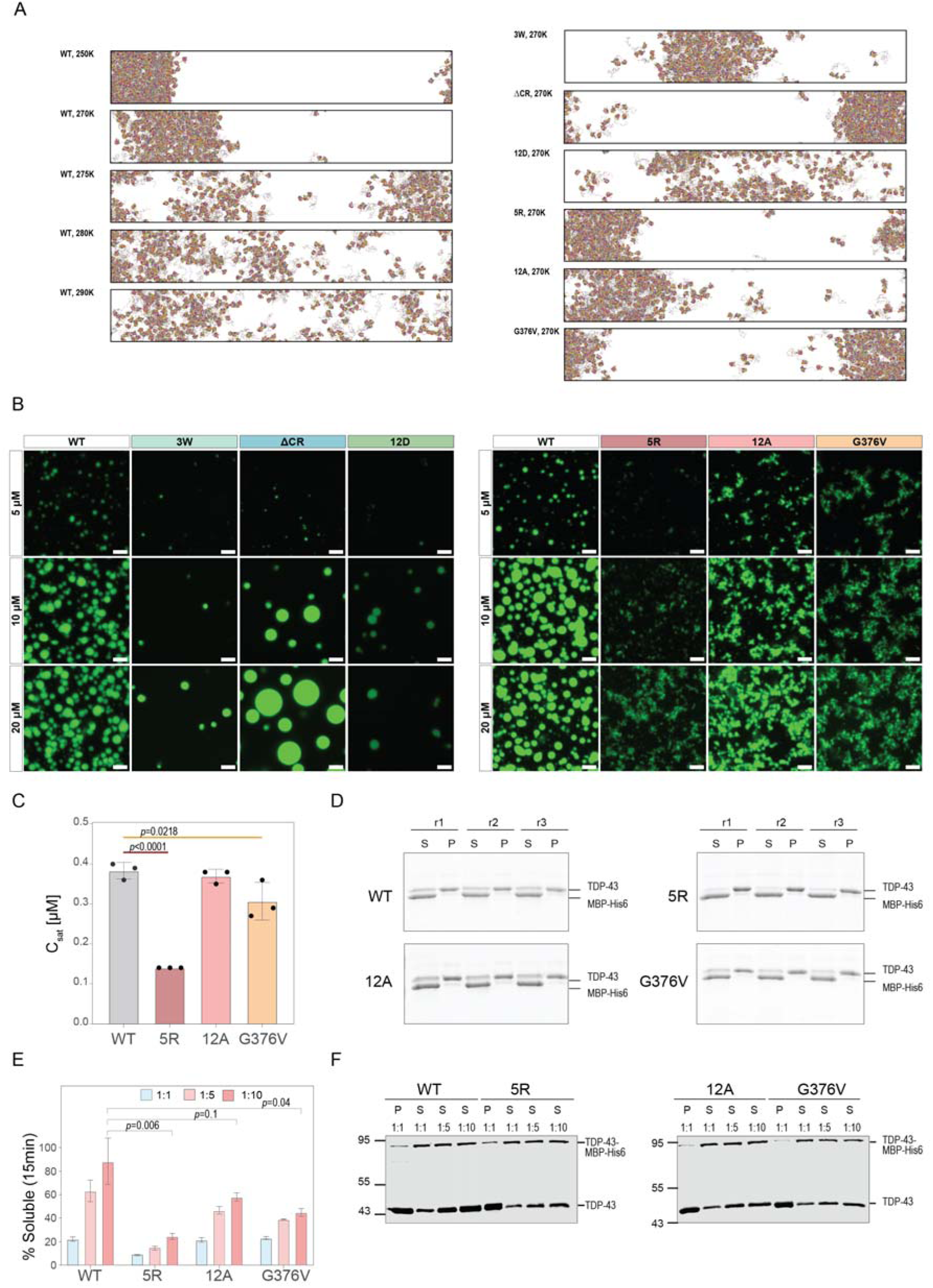
A. Representative visualization of *in silico* condensates generated using simulations with the coarse-grained CALVADOS3 model. For the WT, the last frame of the simulation at different temperatures is shown (250 – 290 K, left panel), while for the other variant simulations the last frame at 270 K is shown (right panel). B. Representative confocal fluorescence microscopy images of condensates formed by TDP-43 variants in comparison to TDP-43 WT at 5, 10 and 20 μM. Imaging was performed 45 minutes after TEV addition. Proteins were fluorescently labeled with Alexa Fluor 488 C5 malemide. Scale bar: 5 µm. C. Saturation concentration (C_sat_) values obtained for condensates formed by TDP-43 WT and solid-like TDP-43 variants. SYPRO Ruby signal from S and P fractions was quantified to obtain a total signal corresponding to 1 μM protein. Afterwards, protein concentration corresponding to the amount of signal in S fraction was quantified and assigned as C_sat_. Three independent experimental replicates were used for the quantification and each replicate corresponds to a C_sat_ value quantified from three phase separation reactions processed on the same day. Error bars indicate standard deviation. Significance was determined by a one-way ANOVA with a Dunnet’s multiple comparisons test to WT. *p*values are indicated. D. Representative images showing SYPRO Ruby-stained gels with TDP-43-containing supernatant (S) and pellet (P) fractions used to estimate C_sat_ values. Three reactions were performed for each C_sat_ estimation replicate and loaded side by side (r1 - 3). E. Quantification of condensate reversibility. Dilution was performed at 15 minutes post condensate induction. % soluble values were determined by quantifying the signal coming from the soluble fraction (S) as a percentage of total signal (P + S fractions) for a given mutant. Error bars indicate standard deviation across three independent experiments. Significance was determined for the signal obtained in 1:10 dilutions using a one-way ANOVA with a Dunnett’s multiple comparisons test to WT. *p*values are indicated. F. Representative western blot images for TDP-43 WT, or the less dynamic 5R, 12A and G376V variants tested for condensate reversibility after 15 minutes of incubation time. Pellet (P) and supernatant (S) fractions were loaded for undiluted (1:1) samples, while for 1:5 and 1:10 samples only supernatant (S) fractions are shown. Blots were incubated with a TDP-43-specific antibody (80001-1-RR, Proteintech).

**Extended Data Fig. 2.**
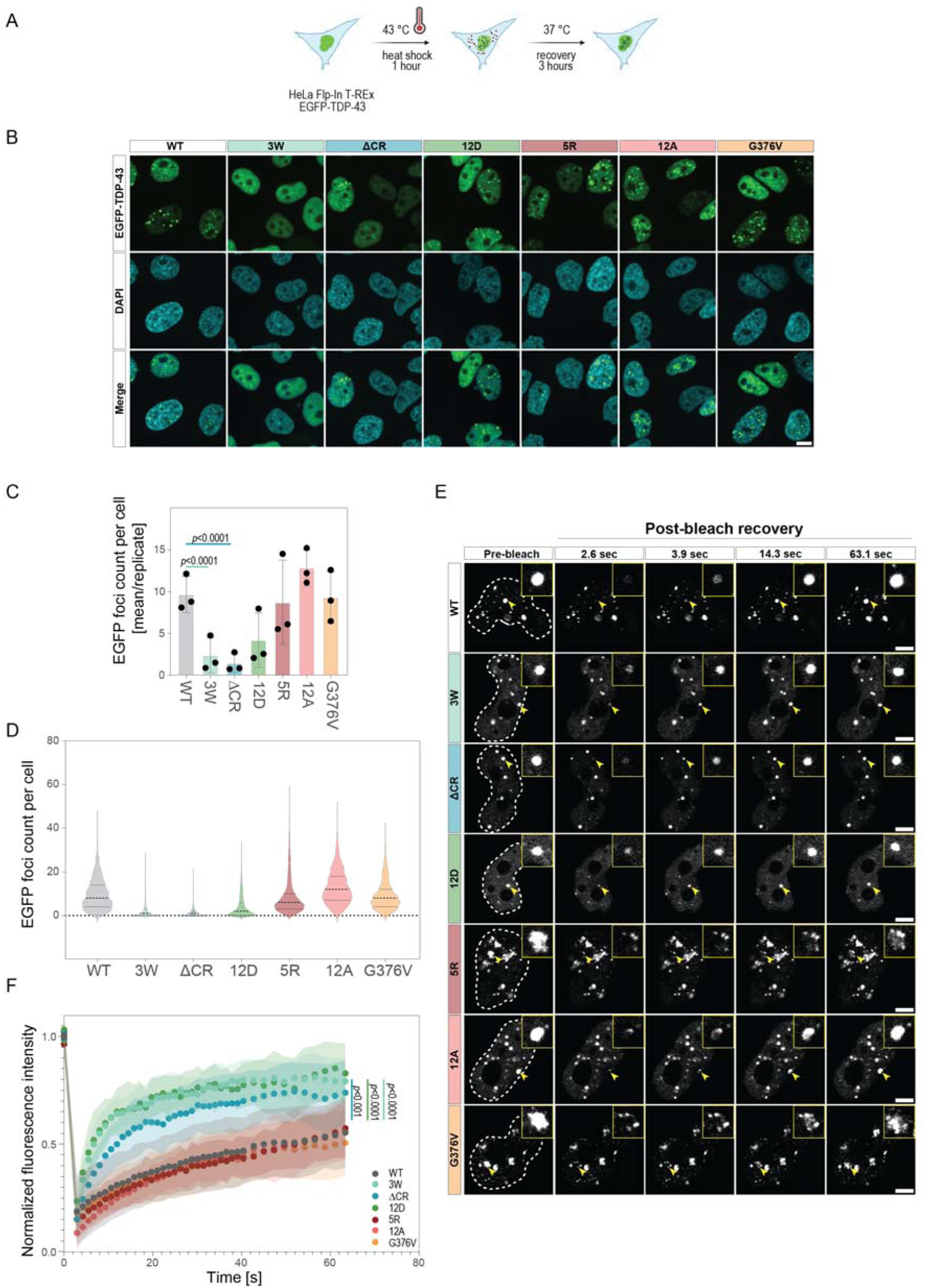
A. Scheme illustrating the heat shock treatment and recovery procedure. Silencing of endogenous TDP-43 with siRNA was done for 48 hours, followed by the doxycycline-induced expression of EGFP-tagged TDP-43 variants for 24 hours. Heat stress treatment was performed for 1 hour at 43°C, followed by a 3-hour recovery period at 37°C). Created in part with BioRender.com. B. Representative confocal images of HeLa Flp-In T-REx cells expressing the indicated EGFP-TDP-43 variants subjected to heat stress as illustrated in (A). EGFP-TDP-43 variants are shown in green. Nuclei were stained with DAPI (blue). Maximum intensity projections are shown. Scale bar: 10 μm. C. Quantification of nuclear stress-induced TDP-43 foci formed during heat shock recovery from confocal high-throughput images shown in (B). Single points show the mean counts of EGFP-TDP-43 foci per cell in each experiment (∼15 fields of view were quantified per replicate). Mean values of three independent experiments are shown in a barplot. Error bars indicate standard deviation. Significance was determined using a one-way ANOVA with a Dunnett’s multiple comparisons test to WT. *p*values are indicated. D. Distributions of foci counts per cell. The median value is indicated by a thick dashed line, 1^st^ and 4^th^ quartiles are shown in dotted lines. ∼2 000 cells were quantified across three replicates. E. Representative FRAP images in HeLa Flp-In T-REx cells expressing the indicated EGFP-tagged TDP-43 variants during heat stress recovery. Point bleaches were performed on individual TDP-43 condensates and are indicated with yellow arrows. Top right quadrants show a zoomed-in image of the bleached condensate. Scale bar: 5 μm. F. Quantification of FRAP in HeLa Flp-In T-REx cells during heat stress recovery. Stress-induced TDP-43 condensates were bleached using a point bleach and their fluorescence recovery was monitored for ∼1 minute. ∼11 cells were analyzed per cell line across three biological replicates, mean values of all fluorescence recovery curves are plotted, with standard deviation indicated in the shaded region. Significance was determined by quantifying areas under the curve for individual condensates followed by one-way ANOVA with a Dunnett’s multiple comparisons test to WT. *p*values are indicated.

**Extended Data Fig. 3.**
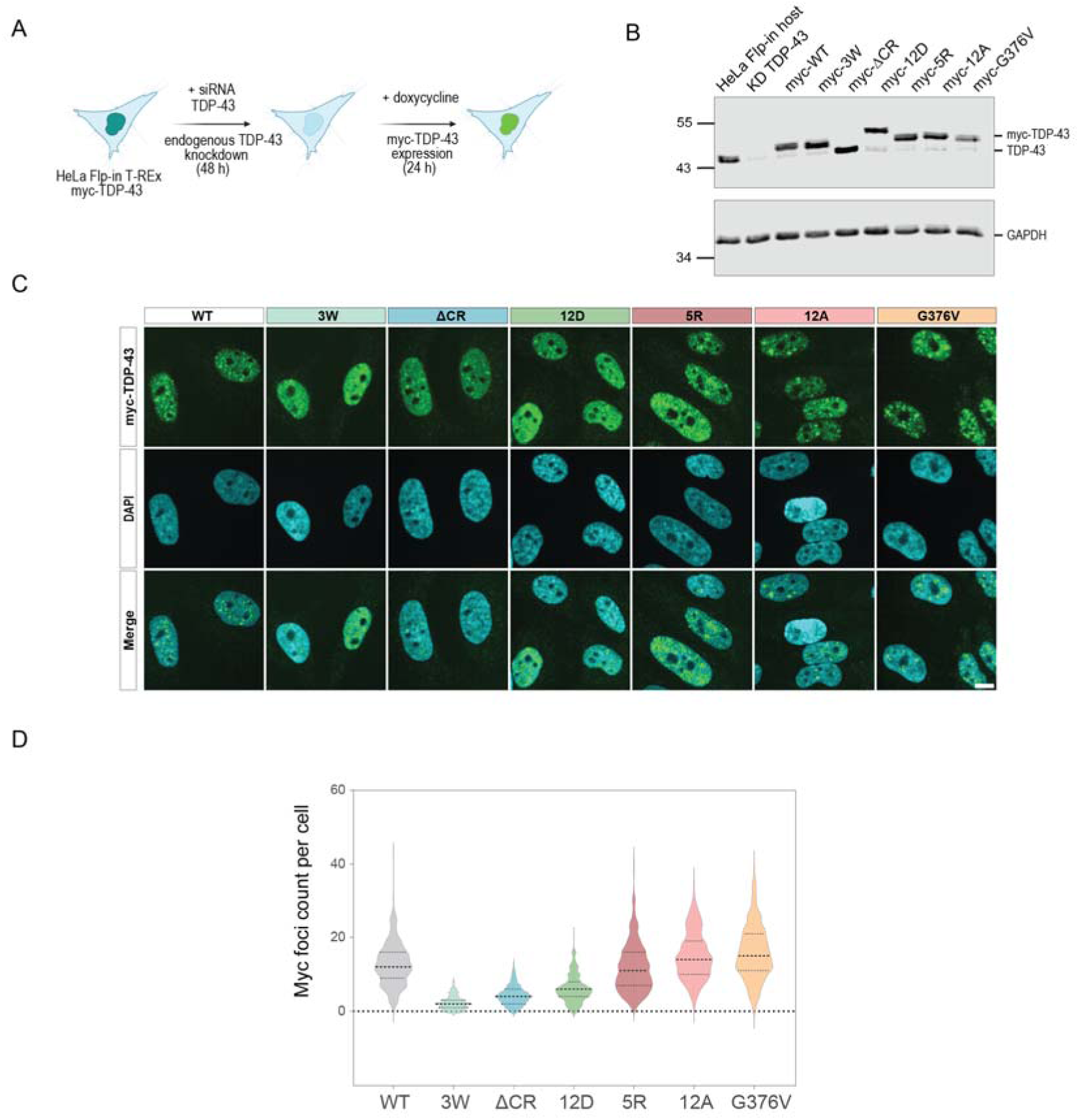
A. Scheme illustrating the experimental setup in all cellular experiments with myc-TDP-43 variants. Silencing of endogenous TDP-43 with siRNA was done for 48 hours, followed by the doxycycline-induced expression of myc-tagged TDP-43 variants for 24 hours. Created in part with BioRender.com. B. Western blot showing myc-TDP-43 expression levels in HeLa Flp-In T-REx cells expressing the indicated myc-TDP-43 variants in the absence of endogenous TDP-43 and after doxycycline induction. Parental HeLa Flp-In T-REx host cells were used to show endogenous TDP-43 levels as well as its depletion after knockdown (KD TDP-43). GAPDH blot is shown as a loading control. Antibodies used: mouse anti-TDP-43 (60019-2-Ig, Proteintech), rabbit anti-GAPDH (10494-1-AP, Proteintech). C. Representative confocal images of HeLa Flp-In T-REx cells expressing the indicated myc-TDP-43 variants stained with an anti-myc antibody (clone 9E10, IMB PPCF, green). Nuclei were stained with DAPI (blue). Maximum intensity projections are shown. Scale bar: 10 μm. D. Quantification of nuclear TDP-43 foci from confocal high-throughput images shown in (C). Violin plots show the distributions of foci counts per cell. The median value is indicated by a thick dashed line, and the 1^st^ and 4^th^ quartiles are shown in dotted lines. Between 86 and 467 cells were quantified.

**Extended Data Fig. 4.**
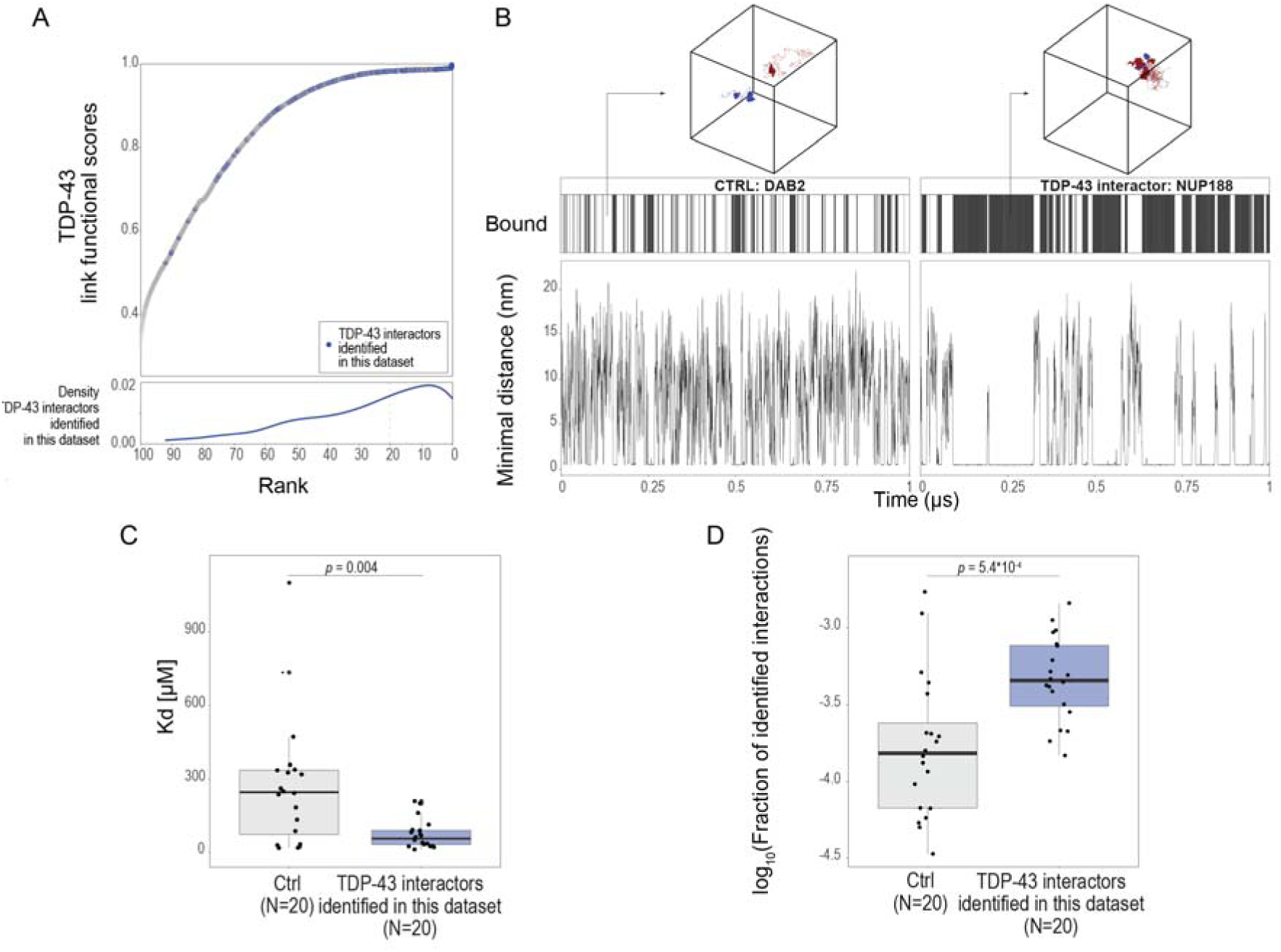
A. Functional association of proteins with TDP-43. 18 841 proteins were ranked based on their functional association with TDP-43. For each protein present in the STRING network (v. 12), the shortest interaction path to TDP-43 was calculated, and a path score was derived as the product of the interaction confidence values along the path. Lower panel shows the distribution of the functional score for all the identified TDP-43 interactors (Fig. 3E). B. Prediction of interactions between TDP-43 and its interacting proteins compared to the control group. Representative examples of interactions are shown between a control protein (DAB2) and a TDP-43 interactor (NUP188). The lower panel displays the minimal distance between the two protein pairs for each frame. The upper panel includes a tile plot, showing the fraction of frames in which interactions are identified. On top is a snapshoot of the simulation between DAB2 (red) and TDP-43 (blue) and between NUP188 (red) and TDP-43 (blue). C, D. Prediction of interactions between TDP-43 and its interacting proteins compared to control pairs. The analysis included 20 control pairs, defined as TDP-43 paired with proteins detected in the AP-MS dataset but not classified as interactors, and 20 interaction pairs consisting of TDP-43 and proteins identified as interactors in the AP-MS dataset. Molecular dynamics simulations for each protein pair were performed using the CALVADOS3 coarse-grained force field model. *In silico* dissociation constant (Kd) was estimated for each protein pair from the ratio of the probability of bound versus unbound states. Statistical significance between groups was assessed using a Wilcoxon rank-sum test (N = 20 per group) (C). The number of contacts normalized to the total number of possible contact pairs was estimated for each protein pair. Statistical significance between groups was assessed by an unpaired two-sided t-test (N = 20 per group) (D).

**Extended Data Fig. 5.**
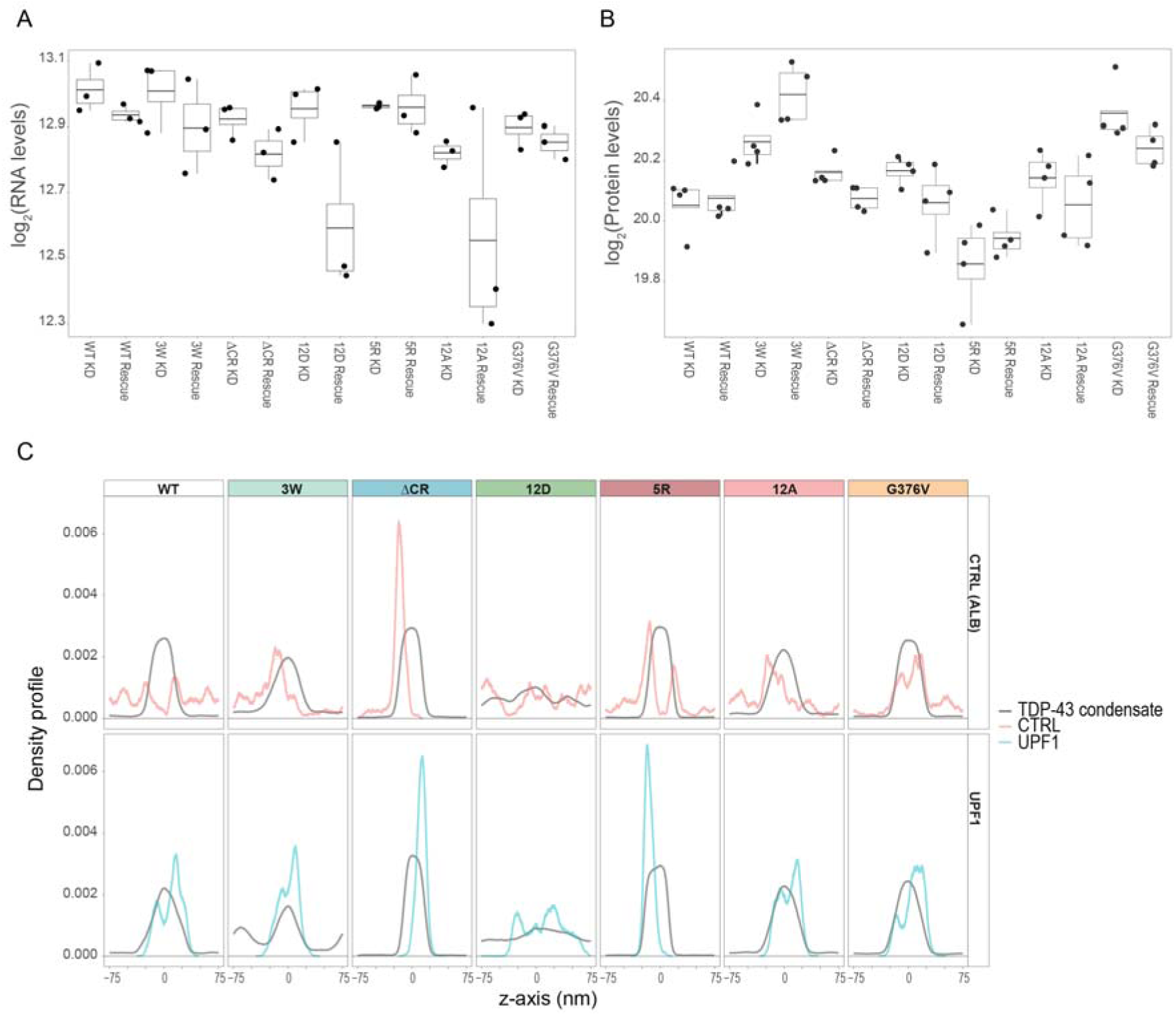
A, B. Quantification of RNA (A) and protein (B) levels of UPF1 across the described conditions. TDP-43 was depleted using siRNA (KD), followed by re-expression of myc-tagged WT, PS-deficient or solid-like TDP-43 variants (rescue). Single points represent biological replicates, and the boxplot reports the distribution of the values with the mean (central line), the interquartile range (1^st^ and 3^rd^ quartile) and whiskers (1.5XIQR, interquartile range). C. Density profiles from molecular dynamics simulations showing co-localization between TDP-43 variants (100 chains, gray) and UPF1 (single chain, cyan) or albumin (control, single chain, red). Profiles were computed from 5 µs trajectories (2 500 frames) and each protein/condensate was normalized to the number of beads and chains to enable comparison of UPF1, albumin and TDP-43 condensates.

**Extended Data Fig. 6.**
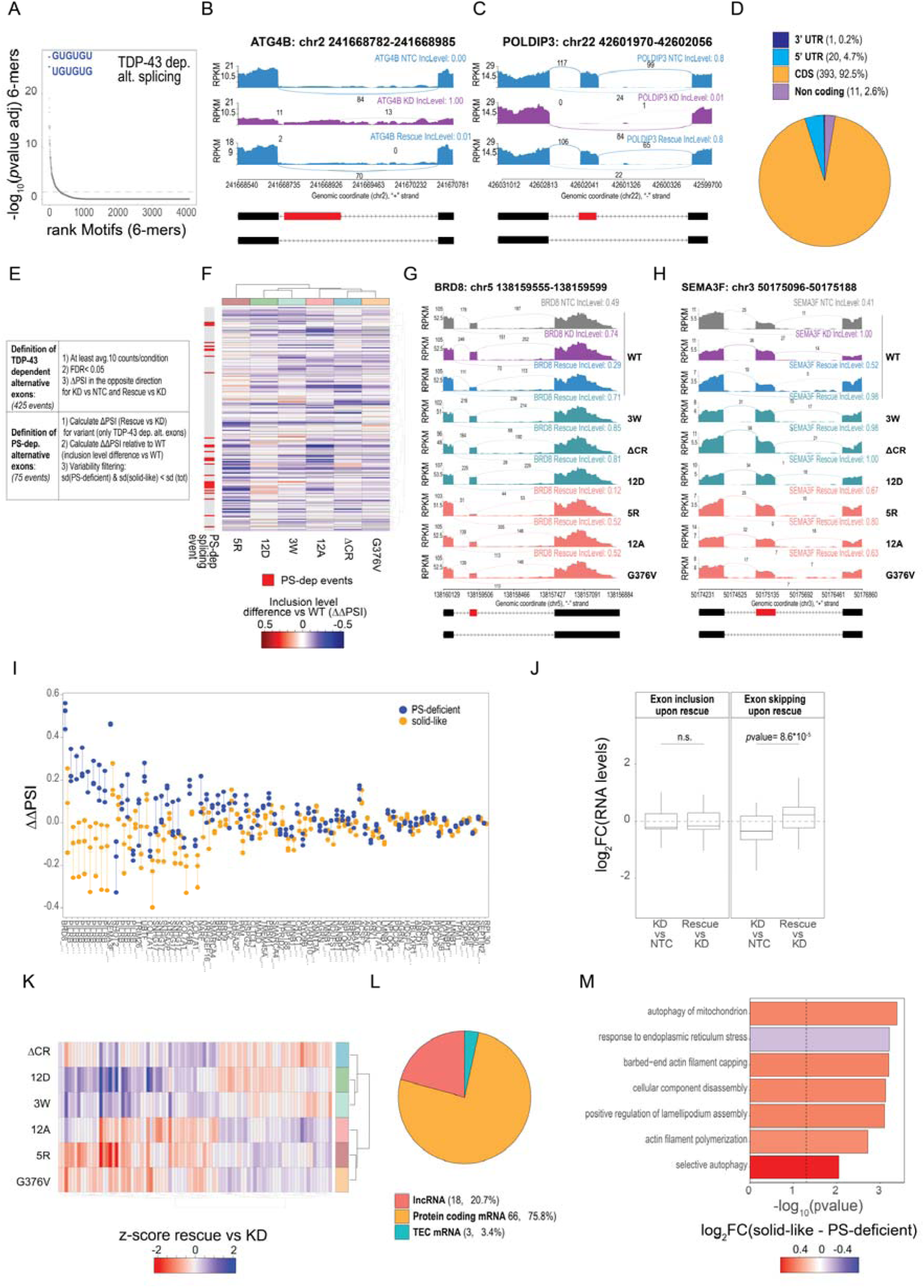
A. Motif enrichment analysis for 4 096 6-mers in TDP-43-dependent alternatively spliced exons (425 events). Significance was calculated using hypergeometric test with flanking regions of unchanged alternative exons as a background. B, C. Sashimi plots showing alternatively spliced exons in *POLDIP3* and *ATG4B*. The plot shows exon coverage and splice junction usage for NTC, KD and rescue in TDP-43 WT cells. D. Annotation of TDP-43 dependent alternatively spliced exons. CDS = coding sequence. E. Summary of the data analysis approach for identifying TDP-43- and PS-dependent alternatively spliced exons. F. Unsupervised hierarchical clustering of 185 exon inclusion events in the rescue versus KD comparison. Values reported in the heatmap show the differences in inclusion levels normalized to WT (ΔΔPSI), with positive values (stronger inclusion compared to WT) in red and negative values (weaker inclusion compared to WT) in blue. In the annotation bar on the left, the events identified as PS-dependent are shown in red. G, H. Sashimi plots showing alternatively spliced exons in *SEMA3F* and, *BRD8*. The plot shows exon coverage and splice junction usage for NTC, KD and rescue in TDP-43 WT cells, and the rescue for all variants analyzed. I. Visualization of 75 PS-dependent alternatively spliced exons. ΔΔPSI values are shown for alternatively spliced exons grouped by PS-deficient (blue) and solid-like conditions (orange). J. Distribution of expression level changes (log₂FC) for genes containing alternatively spliced exons that are included (left panel) or skipped (right panel) upon rescue. Each panel compares KD versus NTC and rescue versus KD conditions. Boxplot reports the distribution of values with the median (central line), the interquartile range (1^st^ and 3^rd^ quartile), and whiskers (1.5XIQR, interquartile range). Statistical significance was assessed using an unpaired two-sided t-test. K. Unsupervised hierarchical clustering of 87 PS-dependent genes following TDP-43 rescue. The heatmap represents z-score differences in gene expression (log_2_FC), with red and blue indicating respectively positive and negative deviations from the mean expression levels of each gene. L. Annotation of PS-dependent genes. M. GO enrichment analysis (Biological Process) for upregulated PS-dependent genes. GO terms significantly enriched (*p*value < 0.01) are grouped based on semantic similarity. Red color represents elevated expression levels of genes associated with the GO term in the solid-like condition, whereas blue indicates higher expression levels in the PS-deficient condition.

**Extended Data Fig. 7.**
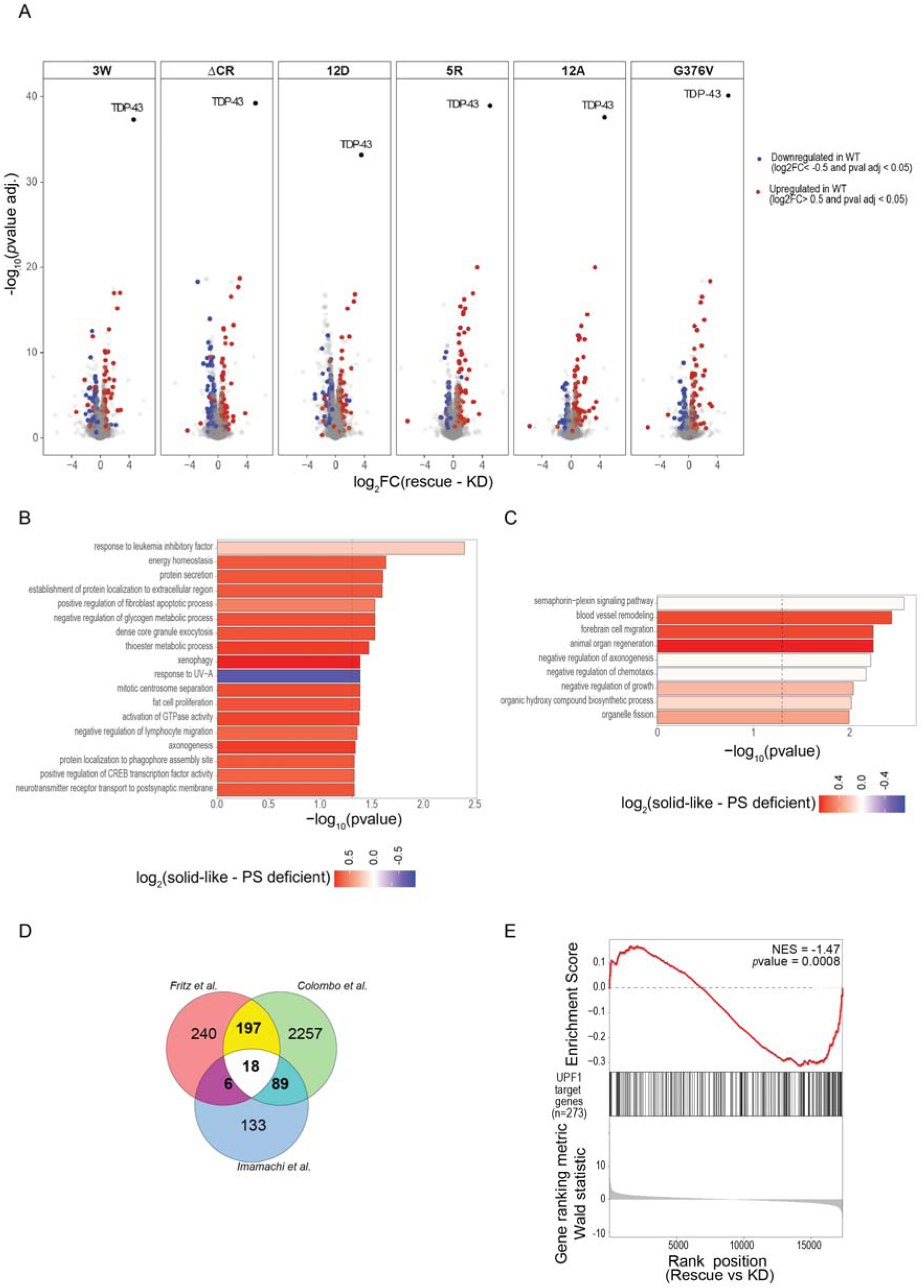
A. Volcano plots showing differences in protein abundance of the TDP-43 variants with distinct PS properties between rescue and KD conditions. The x-axis represents differential protein abundance, while the y axis shows statistical significance (*p*value adjusted). Proteins that were up- or downregulated in WT are shown in red and blue respectively. B, C. GO enrichment analysis (Biological Process) for upregulated (B) and downregulated (C) PS-dependent proteins. GO terms significantly enriched (*p*value < 0.01) are grouped based on semantic similarity. Red color represents elevated expression levels of proteins associated with the GO term in the solid-like condition, whereas blue indicates higher expression levels in the PS-deficient condition. D. Venn diagram showing the overlap of UPF1 target genes from HEK293 and HeLa cells based on the dataset generated by *Fritz et al*., *Colombo et al.*, and *Imamachi et al.* ^108–110^. In bold: the number of genes identified in at least two datasets and used as “UPF1 target genes” in this study. E. Gene set enrichment analysis (GSEA) comparing RNA levels for TDP-43 rescue and knockdown conditions. Gene set: UPF1 target genes in HEK293 and HeLa cells defined in Fig. S15D. The y-axis displays the running enrichment score (ES) (upper panel), the position of UPF1 target genes (middle panel) and corresponding ranked changes calculated with Wald statistic (lower panel); the x-axis shows the ranked position of each gene in the set. NES (normalized enriched score) and *p*value are shown on the plot.

**Extended Data Fig. 8.**
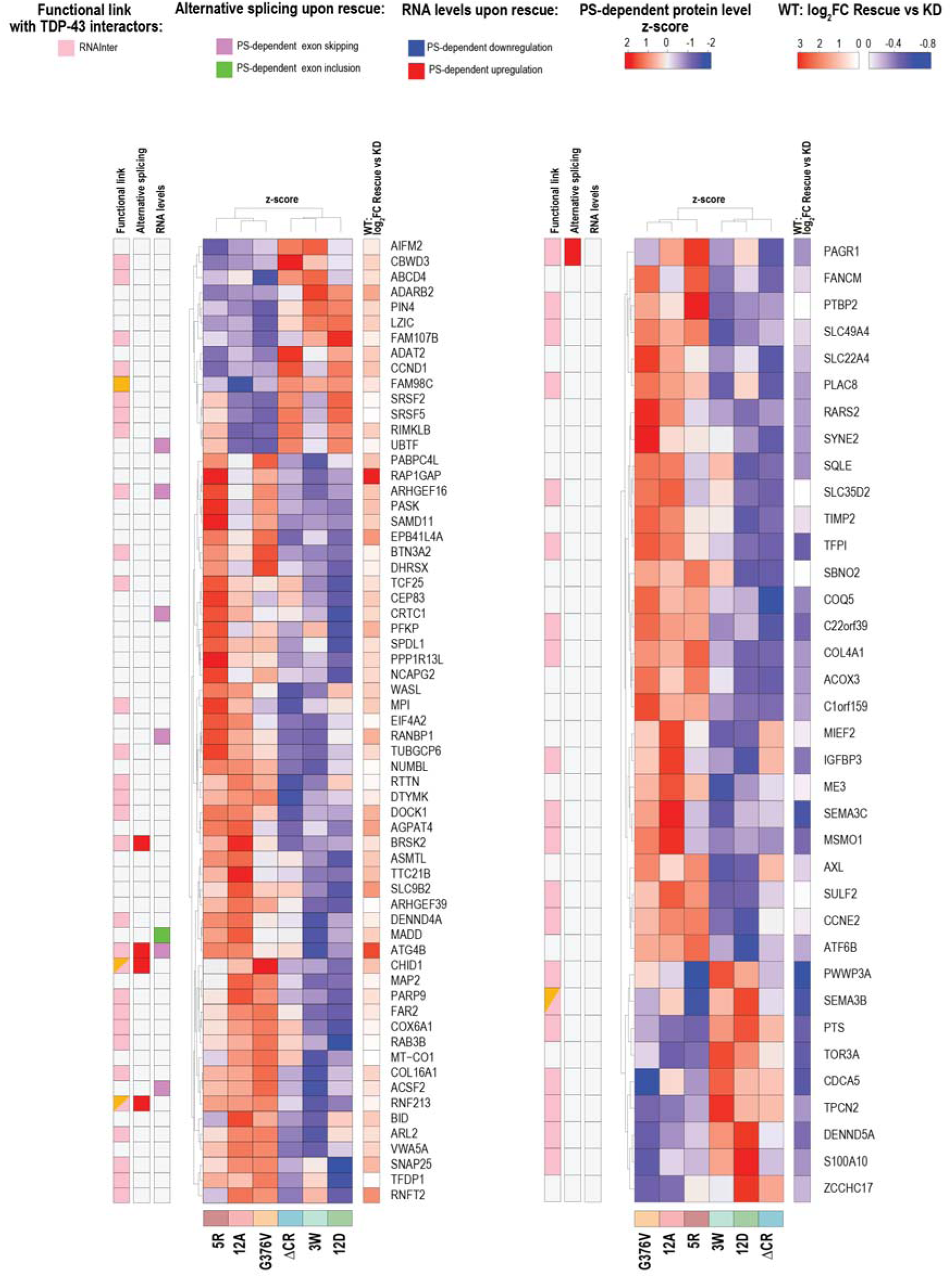
Summary of proteins with abundance levels regulated by PS. The heatmap categorizes proteins as upregulated (left) or downregulated (right) upon rescue with TDP-43 WT. The following layers of information are displayed from left to right:

• Functional link between PS-dependent proteins and PS-dependent TDP-43 interactors based on RNAInter dataset.
• Link between PS-dependent proteins and PS-dependent RNA abundance changes highlighting whether RNA abundance is upregulated (red) or downregulated (blue) upon rescue.
• Link between PS-dependent proteins and PS-dependent alternatively spliced events highlighting whether the protein is associated with exon inclusion or skipping upon rescue.
• Z-scores of protein abundance changes (rescue versus KD). Red and blue color coding indicates higher and lower abundance compared to the average value.
• Differential analysis for the WT condition comparing protein abundance changes between rescue versus KD.

**Extended Data Fig. 9.**
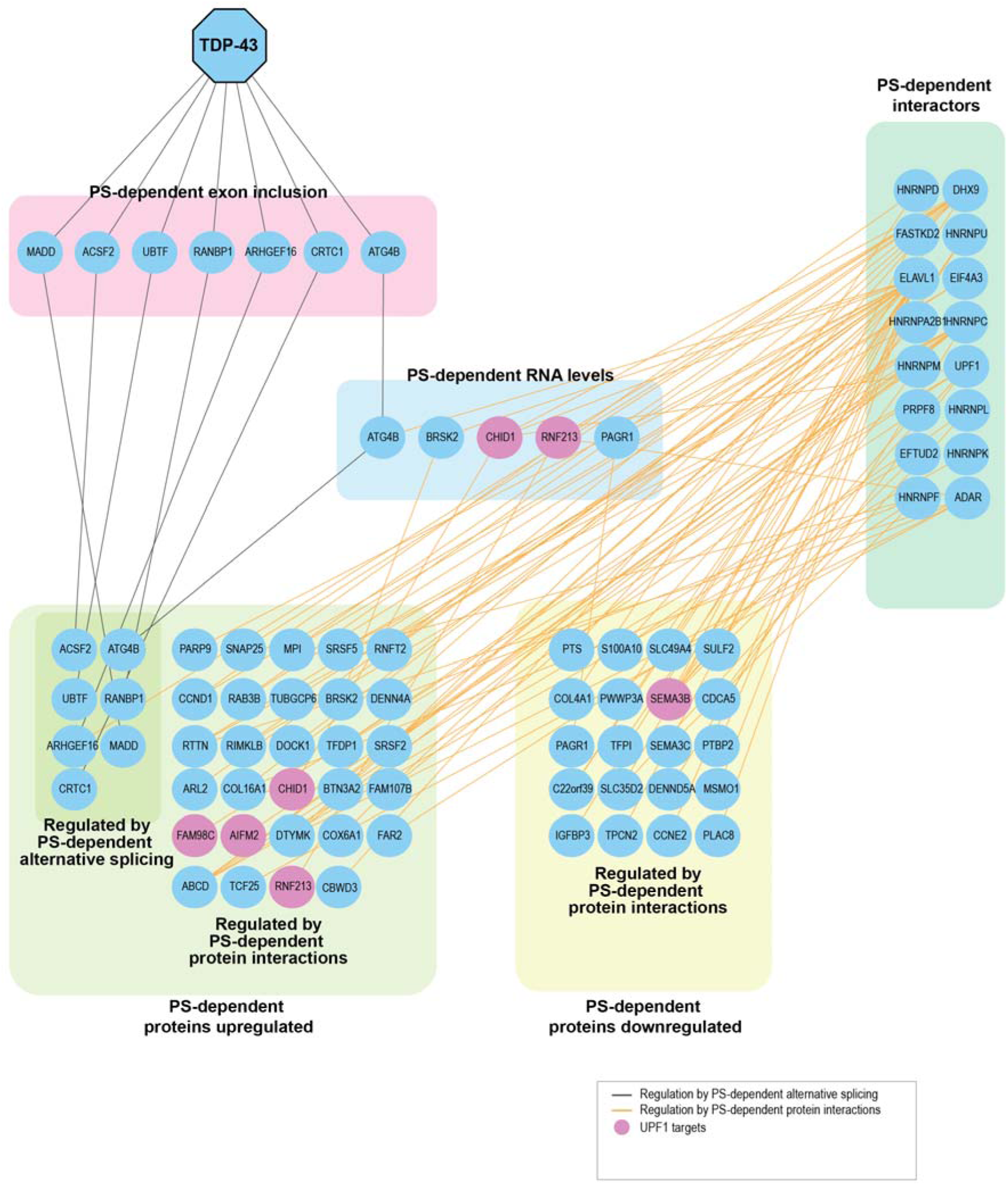
Network representation of PS-dependent events across different levels of gene regulation for proteins with identified links to either PS-dependent alternative splicing changes and/or to PS-dependent TDP-43 interactors (from AP-MS dataset). Splicing-dependent regulation mechanisms are depicted with black lines, while functional associations derived from RNAInter are shown with yellow lines. UPF1 target genes are highlighted in pink.

