## Supplementary Table 1 for "Phase separation behavior of TDP-43 governs its protein interactome and regulation of alternative splicing"

**Table S1**. Mutations introduced in the TDP-43 sequence for each TDP-43 variant.

| TDP-43 variant | Mutations |
|---|---|
| 3W | W334G, W385G, W412G |
| ΔCR | aa 321-340 deletion (AMMAAAQAALQSSWGMMGML) |
| 12D | S373D, S375D, S379D, S387D, S389D, S393D, S395D, S403D, S404D, S407D, S409D, S410D |
| 5R | G295R, Q343R, G348R, G357R, G384R |
| 12A | S373A, S375A, S379A, S387A, S389A, S393A, S395A, S403A, S404A, S407A, S409A, S410A |
| G376V | G376V |
